# Uncertainty-Aware Deep Learning for Multi-Metric and Dose-Specific Prediction of Drug Synergy

**DOI:** 10.1101/2025.11.05.686895

**Authors:** Muhammad Javad Heydari, Bryan Lye, Parvin Mansouri, Thomas Marsland, Kavya Pamulapati, Robert Vary, Hermione Allen, Kaylene J Simpson, John G Lock, James McKenna, Fatemeh Vafaee

## Abstract

Accurately predicting drug synergy is critical to accelerate the development of combination therapies for cancer and other complex diseases. Yet, the vast combinatorial drug and dose space poses a substantial challenge, even for modern deep learning approaches. Existing approaches often lack generalisability, collapse rich dose–response surfaces into single dose-averaged synergy scores, and fail to quantify predictive uncertainty. Here, we introduce AlgoraeOS, a biologically informed, attention-aware deep neural network designed to address these challenges. Trained on the largest harmonised dataset of experimentally tested drug combinations, AlgoraeOS simultaneously predicts multiple synergy metrics, while preserving their empirical correlations and accurately estimating both aleatoric and epistemic uncertainty. The model achieves state-of-the-art performance and strong out-of-distribution generalisability across diverse tissues and drug mechanisms, including rigorous zero- and few-shot evaluations. Notably, AlgoraeOS predicts the entire dose-response surface, providing dose-specific inhibition profiles with high precision and scalability to multi-million–point datasets. Prospective *in vitro* validation of dose-specific inhibition was performed using an anchor compound entirely absent from the training corpus, tested in combination with 24 mechanistically diverse partner drugs across three molecularly distinct cancer cell lines, yielding 2,592 dose–combination measurements. This evaluation demonstrated consistent rank-order fidelity (Spearman’s ρ = 0.51–0.59, p < 0.001) and strong directional agreement (Kendall’s τₐ = 0.863), confirming reliable prediction under out-of-distribution conditions. Model-derived uncertainty estimates further stratified predictions by expected reliability, with lower-uncertainty predictions showing higher concordance with experimental outcomes. By integrating uncertainty-aware, multi-metric, and dose-resolved prediction into a single unified framework, AlgoraeOS offers a powerful solution for drug-combination discovery and establishes a new standard for model development and validation in the field.

## Introduction

Combination therapy has become a foundational strategy in modern oncology, reflecting the clinical necessity of targeting the complex and adaptive nature of cancer ^1,2^. While monotherapies can be initially effective, their success is often transient due to intrinsic intra-tumour heterogeneity and the rapid evolution of acquired resistance through compensatory signalling pathways ^3^. The clinical rationale for using drug combinations extends beyond merely overcoming resistance; it aims to achieve synergistic effects by targeting distinct, complementary mechanisms to improve response rates and extend the duration of patient benefit ^4^. The success of this approach is evident in numerous standards of care, such as the combination of BRAF and MEK inhibitors in melanoma, which have significantly improved patient outcomes ^5^. This clinical imperative is reflected in the trial landscape, where single-agent studies have dropped from ∼70% to 20–30% (2000 – 2021). ^6^ Consequently, the number of clinically validated combinations is substantial, with resources like the OncoDrug+ database documenting over a thousand responsive combinations from clinical sources ^7^. Despite these clinical successes, the preclinical discovery pipeline for identifying new, effective combinations remains a critical bottleneck. The combinatorial space is immensely large, screening a modest library of a few thousand compounds in a pairwise fashion would require millions of experiments, a scale that is prohibitive for even the most advanced high-throughput screening (HTS) platforms ^8^. To address this combinatorial complexity, computational approaches are now critical for prioritising promising combinations and guiding efficient experimental validation.^1,9^

The availability of large-scale datasets, such as DrugComb ^10^, has catalysed a surge in deep learning models for synergy prediction ^11,12^. However, as our comprehensive literature review reveals (**Supplementary File 1**), despite promising reported performance, often reflected in low aggregated error rates or high accuracy, many contemporary approaches exhibit critical limitations that hinder their translational impact. A primary challenge is model generalisability. Many methods are trained on relatively small or homogenous datasets (e.g., the Merck screen ^13^) ^14–16^, which constrains their applicability to broader classes of drugs or diverse cell lines. Moreover, such models can be particularly prone to learning dataset-specific patterns or “shortcuts” rather than capturing the underlying chemical and biological mechanisms of drug interactions ^17^. Consequently, these models often exhibit poor generalisation to out-of-distribution (OOD) data, such as unseen drugs or cell lines, an issue not adequately examined through rigorous zero-shot or few-shot validation settings ^18,19^. This highlights a critical gap: the need for more extensive OOD validation prior to committing to costly experimental follow-up.

Beyond the challenge of generalisability, the predictive frameworks of existing models are often incomplete. First, they are typically designed to predict a single, specific synergy score (e.g., HSA, Bliss, Loewe, or ZIP). However, these metrics are based on distinct mathematical assumptions about drug independence and can lead to differing interpretations of synergy for the same experiment ^20^. Second, most studies overlook the prediction of inhibition rates. While synergy scores quantify the *interaction*, inhibition provides the necessary context on *potency*. A combination may be highly synergistic yet achieve only modest inhibition, making it biologically insignificant, or conversely, display strong inhibition but no true synergistic interaction ^21^. A robust model must therefore predict synergy and inhibition to ensure that prioritised combinations are not only interactive but also therapeutically potent.

Additionally, a major limitation is the general lack of uncertainty quantification: existing methods are often not designed to assign a confidence level to their predictions, failing to estimate the uncertainty arising from noisy data (aleatoric) or from model limitations (epistemic) ^22^. This information is essential for rationally prioritising candidates for preclinical validation, where resources are limited and the cost of failure is high.

Perhaps the most fundamental limitation, however, lies in how these models represent the dose-response relationship itself. Most existing models are trained to predict a single synergy value averaged across a range of dose concentrations. This approach fundamentally misrepresents the dynamic, dose-dependent nature of drug interactions, where a combination may be synergistic at one dose ratio and antagonistic at another. By collapsing the complex dose-response surface into a single metric, these models discard critical information essential for designing clinically effective therapeutic combinations ^23^.

Together, these limitations underscore the need for next-generation computational models that are generalisable, calibrated in their confidence, and capable of predicting the entire dose-response landscape to provide a holistic assessment of both synergy and inhibition.

Here, we introduce **AlgoraeOS**, a biologically informed attention-aware deep neural network developed in partnership with Algorae Pharmaceuticals Ltd to comprehensively address these limitations. Trained on DrugComb, the largest harmonised datasets to date and validated through a rigorous framework that includes out-of-distribution zero-shot and shortcut learning assessments, AlgoraeOS provides a multi-faceted predictive output. Specifically, it is designed to: **i)** precisely predict a full spectrum of synergy scores (e.g., Loewe, Bliss, ZIP, HSA) simultaneously to preserve their empirically observed relationships; **ii)** reliably quantify both aleatoric (data) and epistemic (model) uncertainty to provide a confidence score for each prediction; and **iii)** deliver inhibition predictions in a dose-specific manner. Through this comprehensive approach, AlgoraeOS not only represents the state-of-the-art in synergy prediction from high-throughput screening data but also establishes a new, more rigorous standard for the validation and development of future models in the field.

## Results and Discussion

### Overview of the AlgoraeOS Platform

To predict drug combination efficacy, we developed AlgoraeOS, a lightweight deep neural network (>25 layers, ∼3.5M parameters) that integrates multi-modal features to generate multi-metric synergy predictions with associated uncertainty. The end-to-end architecture of AlgoraeOS is illustrated schematically in **Figure 1**. The platform takes as input representations for two drugs and a cell line. Drug features are derived from both their physicochemical properties (n=1,024) obtained from RDKit ^24^ and contextual SMILES embeddings (n=384) learned by a ChemBERTa model ^25^. The cellular context is represented by normalised and batch-corrected gene expression data from the Cancer Cell Line Encyclopedia ^26^ (CCLE; n=16,720 genes).

**Figure 1.**
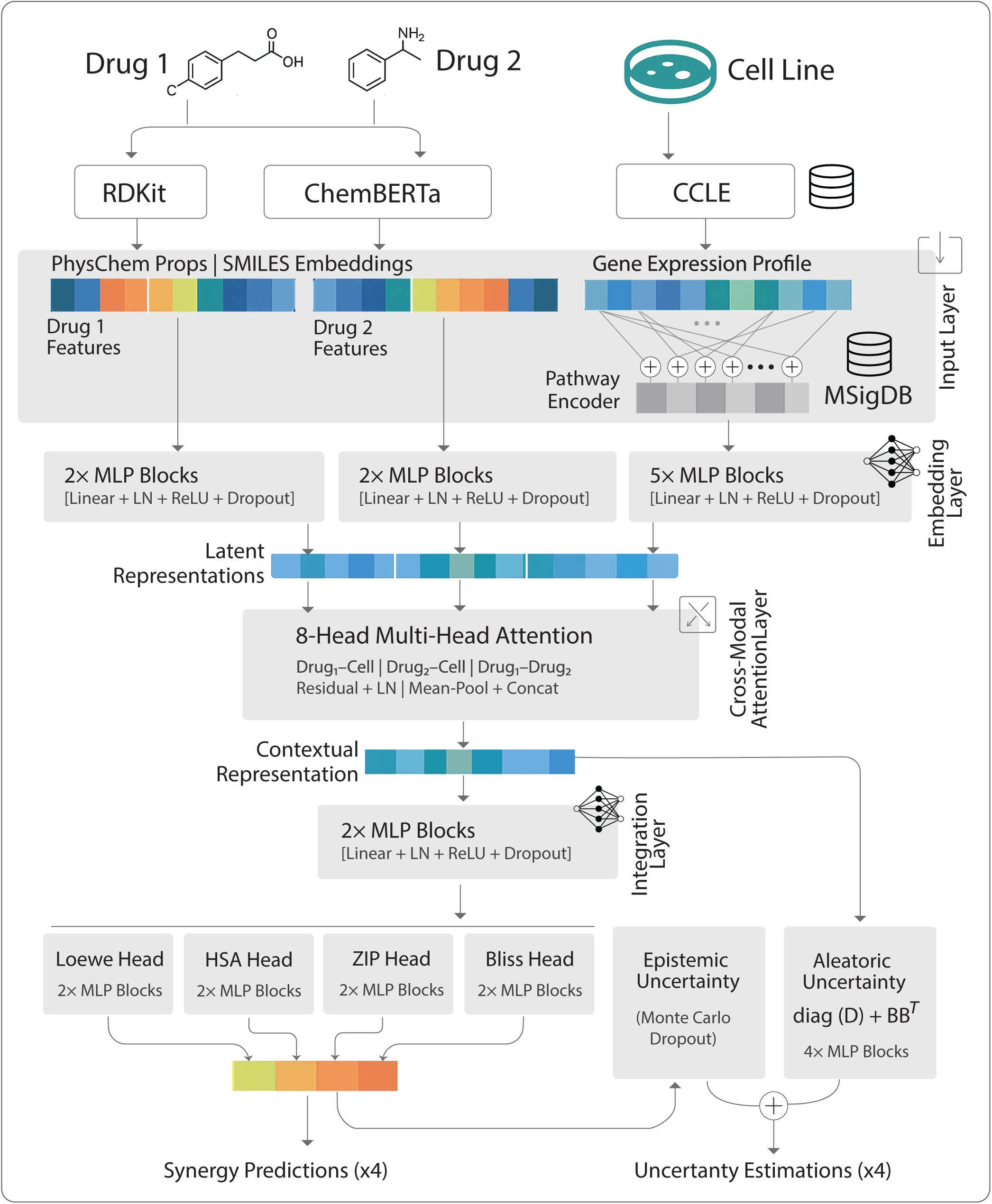
Schematic overview of the AlgoraeOS platform architecture. The end-to-end deep neural network integrates drug features (RDKit properties and ChemBERTa embeddings) with cell line gene expression. The biologically-informed input layer, which aggregates gene expression according to MSigDB pathways, before a multi-head cross-attention mechanism models drug-cell interactions. The architecture branches into two parallel output heads to simultaneously predict four distinct synergy scores (Loewe, Bliss, HSA, ZIP) and their corresponding aleatoric and epistemic uncertainty, providing multi-metric and confidence-aware therapeutic synergy predictions.

AlgoraeOS incorporates a biologically informed input layer for processing gene expression data, whose architecture is explicitly constrained by biological pathways and curated gene signatures from the Molecular Signatures Database (MSigDB) ^27^. In this layer, each neuron represents a specific biological gene set, and the edges encode the inclusion of an input gene within that set. This design reduces the high-dimensional gene expression data into a biologically meaningful, lower-dimensional space. Following this pathway-constrained aggregation, the representations for each input modality are mapped into a 256-dimensional latent space using a series of feed-forward layers equipped with Layer Normalisation, ReLU activation, and dropout regularisation.

To model the complex interdependencies between the drugs and the cellular context, we implemented an 8-head cross-attention architecture. This module captures complementary information through three pairwise configurations: Drug 1 ↔ Cell, Drug 2 ↔ Cell, and Drug 1 ↔ Drug 2. The Drug–Cell attentions encode how each compound’s effect depends on the cellular background, while the Drug–Drug attention facilitates direct interaction between the two drug embeddings, capturing potential synergistic or antagonistic relationships. The outputs from all attention heads are pooled and concatenated to form a unified, contextually-enriched representation. This representation is then branched into two parallel heads: one for synergy prediction and one for uncertainty estimation. The synergy prediction head uses a feed-forward network to fuse the features before parallel MLP towers simultaneously predict four standard synergy metrics (Loewe, Bliss, HSA, and ZIP).

A distinctive feature of AlgoraeOS is its ability to quantify the confidence of its predictions by estimating both aleatoric and epistemic uncertainty ^28^. Aleatoric uncertainty, which reflects inherent data noise, is estimated by a dedicated head trained to model a sample-specific noise covariance matrix across the four synergy scores. In parallel, epistemic uncertainty, which captures the model’s confidence given the training data, is estimated via Monte Carlo Dropout ^29^ during inference. This dual mechanism allows AlgoraeOS to report not only *what* it predicts but also *how confident* it is in that prediction, providing an essential metric for prioritising candidates for experimental validation.

### Exploratory Data Analysis and Drug Feature Selection

Our model was trained and evaluated on a comprehensive dataset curated from DrugComb ^10,30^, the largest publicly available repository of high-throughput drug combination screening data. This resource integrates and harmonises results from multiple large-scale studies, such as ALMANAC ^31^, Merck ^13^, FORCINA ^32^, and CLOUD ^33^. Starting with DrugComb V1.5, we applied a rigorous curation pipeline to ensure data quality and consistency. We focused exclusively on dual therapy records, retaining only those whose cell lines were present in the Cancer Cell Line Encyclopedia (CCLE) DepMap dataset for complete molecular annotation. To maintain data integrity, entries with discrepancies in drug SMILES across repositories or missing values for any of the four primary synergy metrics were excluded. This curated dataset comprised 368,123 unique experimental records, spanning 3,139 drugs and 170 cell lines. To ensure the model’s predictions are insensitive to the input order of the drugs, we performed reverse augmentation, doubling the final dataset size to 736,246 records. This augmentation was performed after the initial train-test splits for all validation strategies to prevent any information leakage from the test set into the training data.

DrugComb quantifies drug–drug interactions using four widely adopted reference models: ^34^ Bliss independence, Highest Single Agent (HSA), Loewe additivity, and Zero Interaction Potency (ZIP). Each defines a different baseline expectation of non-interaction. The Bliss model assumes that drugs act independently at the effect level, combining their inhibitory probabilities. HSA offers a conservative criterion, defining synergy only when the observed combination effect exceeds the maximum inhibition of either single agent. Loewe additivity models the scenario in which two agents behave as dilutions of the same compound, quantifying deviation from additive behaviour. Finally, the ZIP model integrates the Bliss and Loewe principles, evaluating deviations in both potency and shape from an expected non-interactive dose–response surface. By training AlgoraeOS to predict all four complementary metrics, we enable a comprehensive and mechanism-agnostic evaluation of synergy, capturing a richer view of the drug interaction landscape than single-metric models.

**Figure 2a–d** illustrates the empirical distributions of the four synergy metrics across all drug combinations. All metrics are approximately bell-shaped around zero, reflecting predominantly additive interactions. Loewe and, to a lesser extent, HSA display left-shifted distributions, indicating a bias toward antagonism, while ZIP and Bliss are more symmetrically distributed. The narrower, sharply peaked shapes relative to their Gaussian fits suggest leptokurtic tendencies, reflecting a predominance of near-additive interactions with relatively few extreme synergy or antagonism events. **Figure 2e** presents pairwise correlations among the metrics. Strong concordance is observed between ZIP and Bliss (PCC = 0.846) and between HSA and Bliss (PCC = 0.713), consistent with their shared dose–response additivity assumptions. In contrast, Loewe correlates weakly with the others (PCC = 0.224 – 0.311), reflecting its distinct probabilistic independence framework. Together, these results highlight the non-interchangeability of synergy models and the importance of integrating multiple metrics for balanced interpretation.

**Figure 2.**
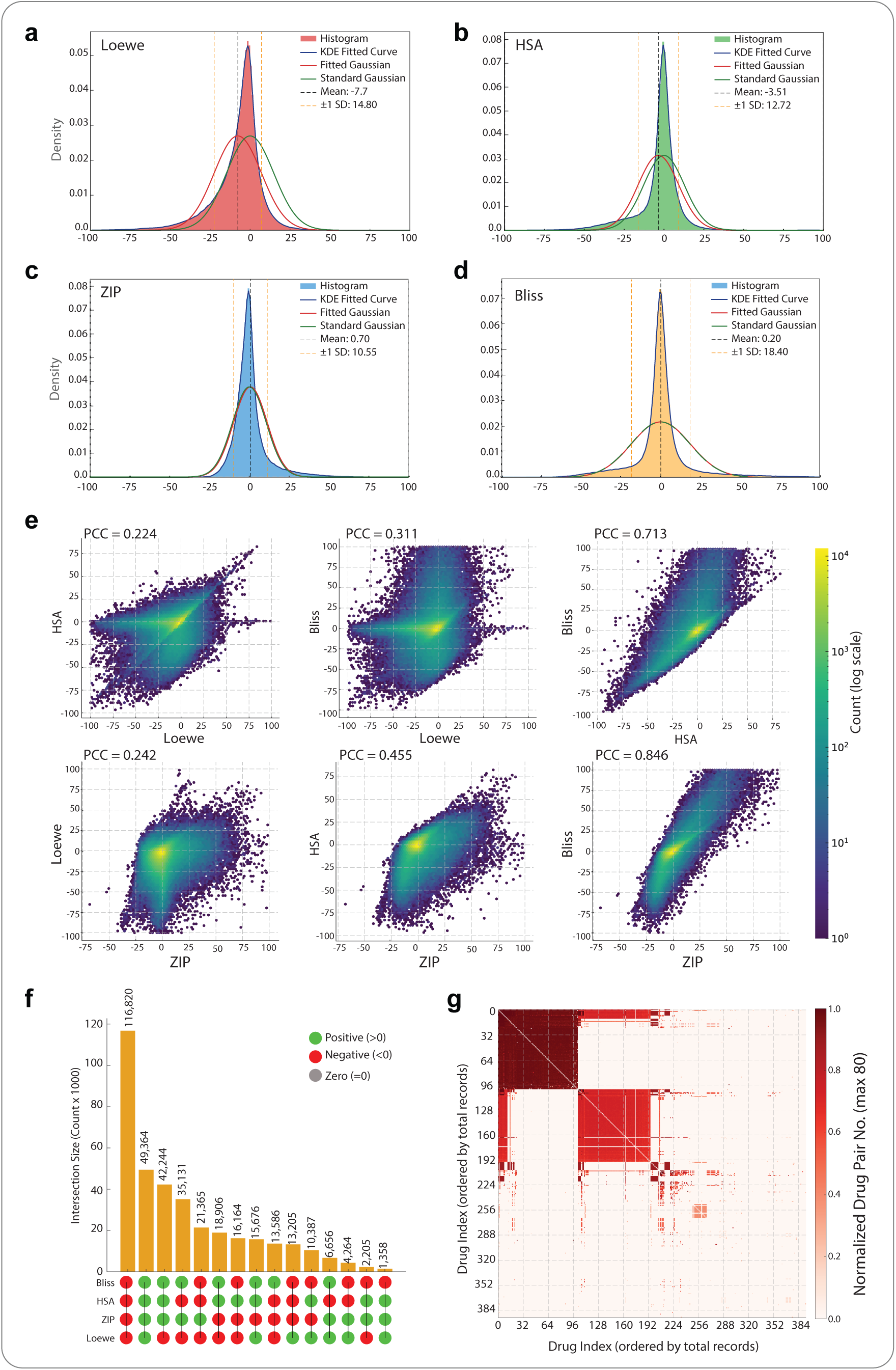
(a–d) Distributions of synergy scores under four reference models. Panels show Bliss (a), HSA (b), Loewe (c), and ZIP (d). Coloured bars depict the histogram of scores. The blue curve is the Kernel Density Estimation (KDE) fit (non-parametric density). The red curve is the fitted Gaussian (normal density with mean and SD estimated from the data). The green curve is the standard Gaussian (mean = 0, SD = 1) shown for baseline comparison. The black dashed line marks the sample mean; orange dashed lines indicate ±1 SD around the mean. **(e)** Pairwise correlations among the four major synergy scoring models (Bliss, HSA, Loewe, and ZIP) computed across all drug combination experiments. Each panel shows a density scatter plot with the corresponding Pearson correlation coefficient (PCC). **(f)** UpSet plot showing the top 15 categories of synergy score concordance across metrics when classified as synergistic (> 0, green) or antagonistic (< 0, red). Grey denotes additive interactions (= 0), which were not among the top 15 categories and therefore do not appear in the plot. **(g)** Heatmap showing dataset coverage across all drug pairs with more than 10 experimental records. Each axis represents drugs ordered by their total record count, and colour intensity reflects the number of experiments (normalised by a maximum of 80) performed for each drug pair.

**Figure 2f** summarises categorical overlaps of synergy classifications (> 0 as synergistic, < 0 as antagonistic) across metrics. The largest intersection corresponds to consensus antagonistic predictions (116,820 combinations), followed by full concordance on synergistic outcomes (49,346). The next major group (42,244) represents combinations deemed synergistic by Bliss, HSA, and ZIP but antagonistic by Loewe, underscoring Loewe’s distinct tendency toward antagonistic classification. Although the > 0 and < 0 thresholds are coarse and values near zero are effectively additive, this pattern reflects the stringency of Loewe’s dose additivity assumption, where drugs are treated as functionally interchangeable. Violations of this assumption, particularly for combinations involving mechanistically distinct and non-interchangeable agents, can lead Loewe to classify effects as sub-additive or antagonistic even when other models detect synergy. These results reinforce the non-interchangeability of synergy models and highlight the importance of integrating multiple reference frameworks for robust interpretation.

Beyond the analyses of synergy metric behaviour, **Figure 2g** shifts focus to the experimental landscape itself, illustrating how comprehensively drug pairs are represented in the dataset. The heatmap visualises the coverage of the curated dataset across all drug pairs with more than 10 experimental records. Each axis represents drugs ordered by their total record count, and the colour intensity indicates the normalised number of experiments conducted for each pair. The heatmap reveals a markedly uneven distribution: a subset of drugs has been extensively profiled across numerous combinations and cell lines (dense dark-red blocks), while most potential pairs remain sparsely sampled or entirely untested. This non-homogeneous coverage demonstrates that, although the dataset constitutes the most comprehensive resource currently available, it still represents only a limited and uneven fraction of the possible combinatorial drug space.

### AlgoraeOS Performance Across Synergy Metrics, Tissues, and Mechanisms

The predictive performance of AlgoraeOS was evaluated using 5-fold cross-validation. The curated dataset was randomly partitioned into five folds, with each fold serving once as a test set while the model was trained on the remaining four. This procedure was repeated five times, and performance metrics were averaged across all folds. Model training employed the Adam optimiser with a Mean Squared Error (MSE) loss function for 600 epochs (base learning rate = 1e-3, dropout rate = 0.1). All computations were performed on the Gadi HPC cluster at Australia’s National Computational Infrastructure (NCI) using four NVIDIA Tesla V100-SXM2-32 GB GPUs.

The cellular context was encoded using normalised, batch-corrected gene expression profiles from the DepMap CCLE dataset (n = 16,720 genes). For drug input features, we investigated 16 distinct molecular representation strategies (**Supplementary Table 1**), encompassing expert-defined structural fingerprints, learned embeddings, physicochemical descriptors, and functional profiles that collectively capture chemical structure, bioactivity, and systems-level context. From this comprehensive pool, four candidate feature blocks were shortlisted based on literature support, information complementarity, and diversity (**Table 1**): PubChem fingerprints ^35^, Morgan/ECFP fingerprints ^36^, RDKit physicochemical descriptors ^24^, and a pretrained ChemBERTa transformer embedding ^25^ (trained on 77 million SMILES). An ablation study was then conducted to quantify the contribution of each representation to predictive performance (**Supplementary Table 2**). Accordingly, ChemBERTa (n = 384 features) and RDKit descriptors (n = 1,024) were selected as complementary feature sets for all downstream modelling and validation.

**Table 1.** Short-listed drug representations used for ablation analysis and final model selection.

| Feature block | What it is / how it is built | Information it carries |
| --- | --- | --- |
| **PubChem FP (881 bits)** | Large substructure key used by PubChem; includes ring topologies, atom pairs, and hybridisation. | Rich mix of functional groups, ring statistics, and atom-pair contexts. |
| **Morgan/ECFP (1024 bits)** | Radius-based circular fingerprint that hashes each atom's neighbourhood. | 3-D-agnostic "chemical neighbourhoods" up to a chosen radius. |
| **RDKit (269 floats)** | Calculated physicochemical descriptors: MW, logP, TPSA, H-bond counts, etc. | Bulk properties governing ADME, solubility, and permeability. |
| **ChemBERTa (384 dims)** | Transformer model pretrained on 77 M SMILES; mean-pooled token embeddings. | Contextual, learned chemical "language" embedding captures structural semantics. |

**Figure 3a** summarises the model’s predictive performance across four synergy metrics, evaluated by Mean Absolute Error (MAE), R², and Pearson’s Correlation Coefficient (PCC). AlgoraeOS achieved strong and comparable performance for Bliss, ZIP, and HSA, while the prediction of the Loewe score was consistently less accurate. This discrepancy likely reflects the weaker correlation of Loewe with the other metrics. Within the joint optimisation framework, shared signal structure across the more correlated metrics reinforces learning, while the distinct assumptions underlying Loewe reduce its alignment with these shared representations.

**Figure 3.**
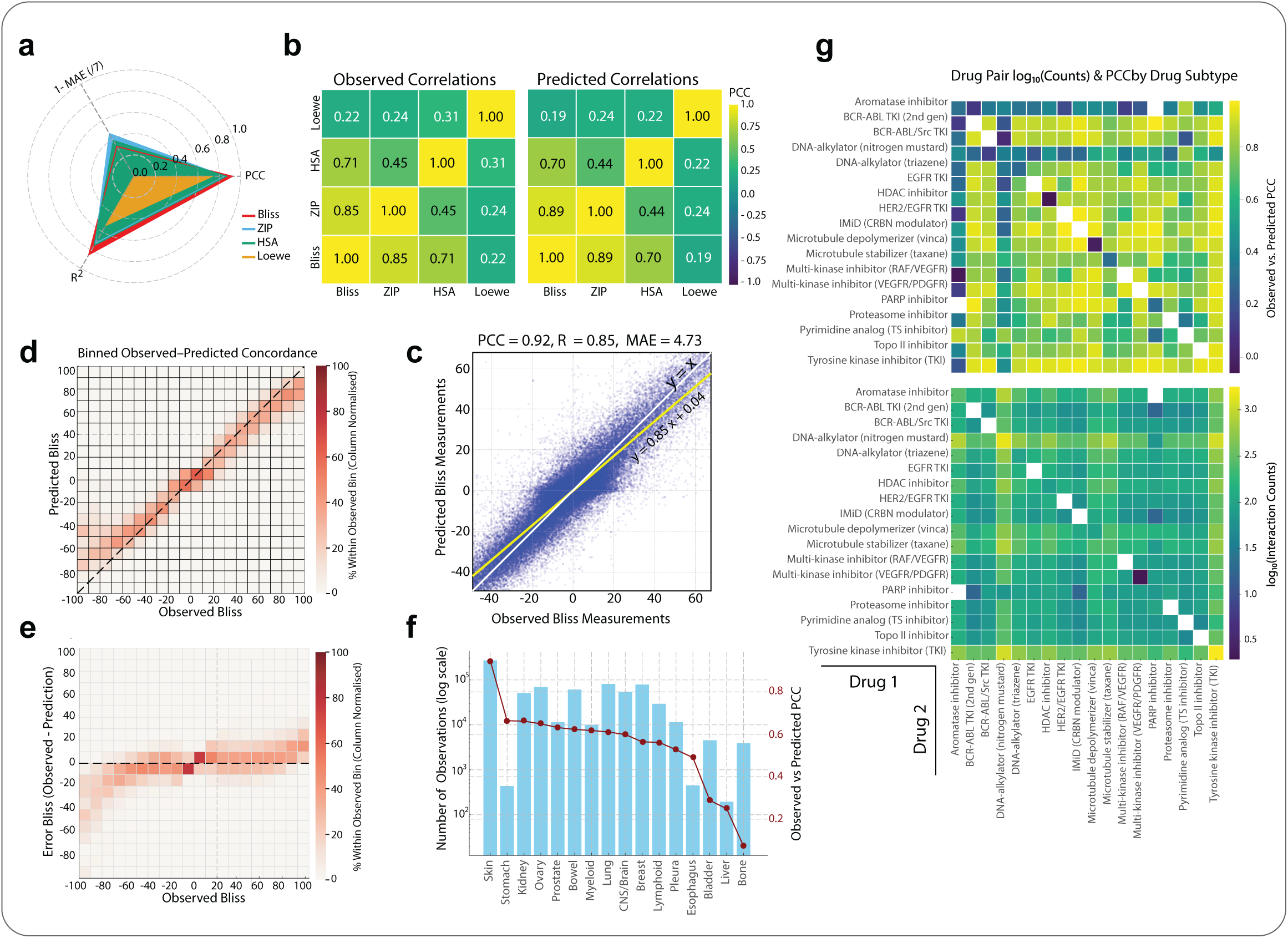
Predictive performance of AlgoraeOS across synergy metrics, tissues, and drug classes. **(a)** Radar plot showing average model performance across the four synergy metrics (ZIP, Loewe, HSA, and Bliss) evaluated using Pearson’s correlation coefficient (PCC), coefficient of determination (R²), and mean absolute error (MAE). **(b)** Heatmaps showing pairwise correlations between the observed (left) and predicted (right) synergy metrics. **(c)** Scatter plot comparing predicted versus observed ZIP synergy scores across all test folds; the yellow line represents the identity line (y = x), and the grey line shows the fitted regression. **(d)** Binned concordance between observed and predicted ZIP values (bin size = 10), with colour intensity indicating the percentage of samples within each bin. **(e)** Binned error distribution (observed – predicted ZIP values) showing deviation across the observed range. **(f)** Bar and line plot showing the number of observations per cancer tissue type (blue bars, log scale) and corresponding PCC between observed and predicted ZIP scores (red line). **(g)** Heatmaps showing the Pearson correlation coefficient (top) and the log₁₀-transformed number of training instances (bottom) for drug pairs grouped by their mechanism of action (MoA) class.

Importantly, the model preserved the observed inter-metric relationships, maintaining the same correlation patterns in its predictions as seen in the experimental data (**Figure 3b**). Given that Bliss showed the highest concordance with other metrics, subsequent detailed analyses are presented for Bliss (**Figure 3c–g**), with corresponding results for HSA, ZIP, and Loewe provided in **Supplementary Figures 1–3**.

**Figure 3c** shows a scatter plot comparing predicted versus observed Bliss synergy scores across the test folds. Each point represents an individual drug–cell–drug combination, with the yellow line indicating the identity line (y = x) and the fitted regression line shown in grey. The model demonstrates strong predictive performance (PCC = 0.92, R² = 0.85, and MAE = 4.73), indicating that predicted synergy values closely align with experimentally measured Bliss scores. Similarly, **Figure 3d** shows the binned concordance between observed and predicted-Bliss scores, where values were grouped in bins of size 10. This representation highlights that predictions align closely with observations across most ranges, including both synergistic and antagonistic extremes, confirming that model accuracy is not dominated by the more abundant additive (near-zero) values. Additionally, **Figure 3e** presents the corresponding binned error distribution (observed – predicted), illustrating that error magnitudes increase slightly toward the extreme tails, particularly in the antagonistic region, as expected due to lower data density. Importantly, the model rarely confuses synergistic and antagonistic cases, with most mispredictions occurring near the additive region (between –10 and 10). Together, these analyses demonstrate that AlgoraeOS maintains consistent predictive accuracy across the entire synergy range, including the biologically most relevant high- and low-interaction regions.

Building on the overall predictive performance observed across all combinations, we evaluated how model accuracy varies across different cancer tissue types (**Figure 3f**). While the model maintains generally high predictive performance across tissues, the results reveal that accuracy is influenced by data availability. Tissues with extensive data coverage, such as skin, kidney and ovary, achieve high concordance (PCC > 0.7), whereas tissues with limited representation, such as liver and oesophagus, exhibit reduced predictive consistency. This pattern underscores the dependence of model generalisability on dataset size, indicating that richer experimental coverage leads to more stable and reliable predictions across biological contexts.

Extending this analysis, **Figure 3g** evaluates model performance across drug mechanism-of-action (MoA) classes (see **Supplementary File 2** for detailed drug classifications). Overall, the model demonstrates high concordance across most MoA categories; however, predictive performance is inversely related to data availability, with drug classes represented by fewer training examples showing reduced correlation. This analysis provides insight into how combinations across different pharmacological classes perform and can guide the interpretation of model predictions for novel drug pairs lacking experimental ground truth.

### Out-of-Distribution Generalisability and Zero-/Few-Shot Evaluation

While the preceding analyses assessed model accuracy under standard cross-validation, these represent in-distribution evaluations, where training and test data are drawn from similar distributions. Testing out-of-distribution (OOD) generalisation is the most direct approach to identifying shortcut learning ^17^. To rigorously assess OOD generalisability, we evaluated AlgoraeOS under progressively more challenging scenarios, examining its ability to extrapolate to rare or unseen drug combinations.

*Few-Shot Evaluation*. We first assessed model robustness under a few-shot learning setting, categorising drug or drug-pair representation frequencies in the training data as rare (≤10), low (11–50), medium (51–200), or high (>200). As shown in **Figure 4a**, the model maintained consistent predictive performance across categories, with only a modest increase in median error and interquartile range for rare combinations.

**Figure 4.**
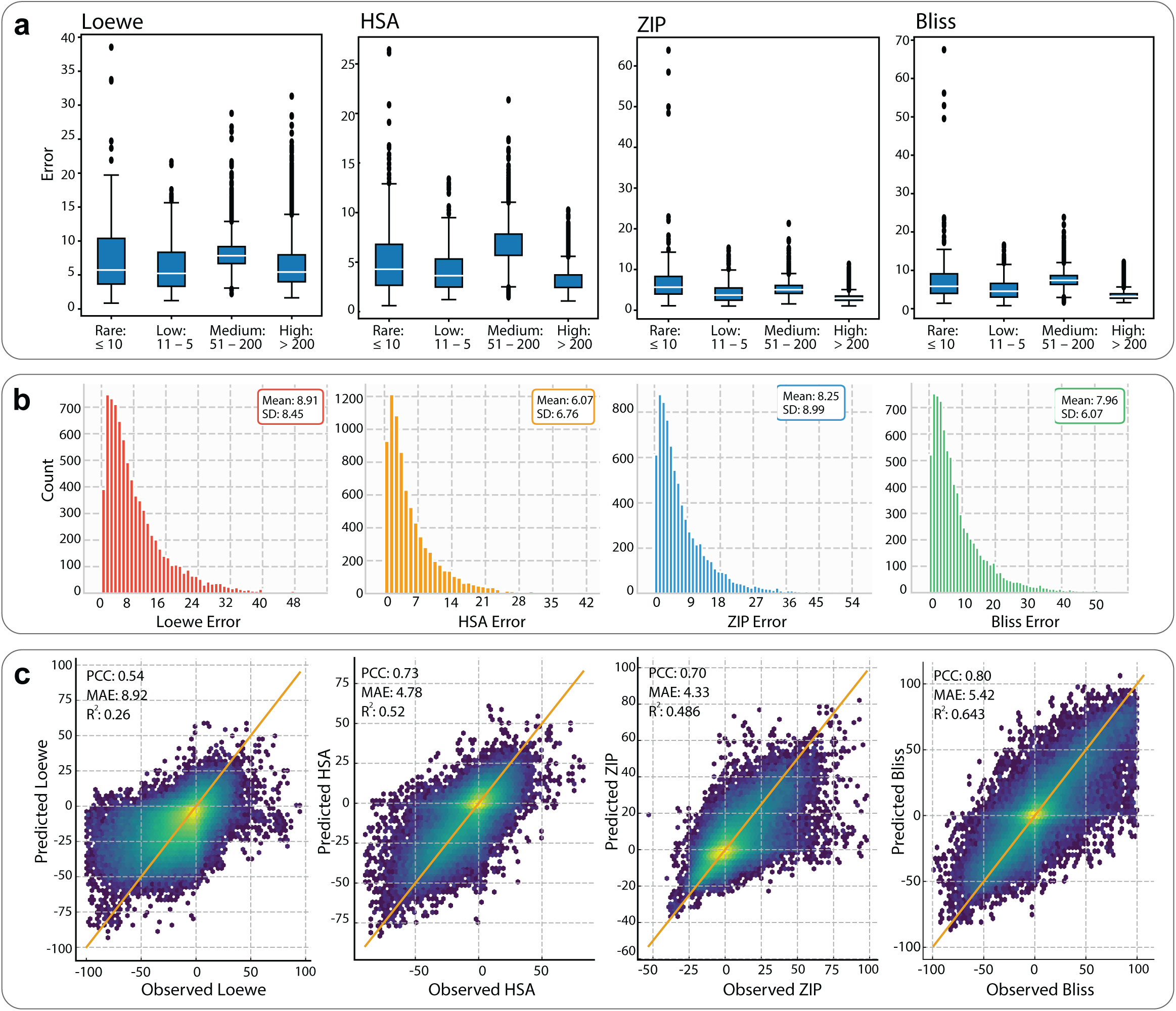
Out-of-distribution (OOD) generalisability and few-/zero-shot performance of AlgoraeOS. **(a)** Boxplots showing prediction error across drug-pair frequency categories in the training set: rare (≤10), low (11–50), medium (51–200), and high (>200) records, for each synergy metric (Bliss, HSA, Loewe, ZIP). **(b)** Distribution of absolute prediction errors (|Observed – Predicted|) for drug combinations unseen during training, illustrating zero-shot prediction capability. The mean and standard deviation of the errors are indicated for each metric. **(c)** Hexbin scatter plots comparing observed and predicted synergy scores under the leave-combination-out OOD setting, in which drug pairs with moderate or high similarity to training pairs (0.6 ≤ Tanimoto ≤ 0.7 and 0.7 ≤ cosine similarity ≤ 0.8) were excluded. Each plot reports the Pearson correlation coefficient (PCC), mean absolute error (MAE), and coefficient of determination (R²) for the corresponding synergy metric.

*Zero-Shot Evaluation*. To test zero-shot capability, we examined drug combinations completely absent from the training set and analysed the absolute prediction errors across all synergy metrics (**Figure 4b**). The error distributions for all four metrics were sharply centred near zero, indicating that most predictions closely approximate experimental values despite the absence of prior exposure to those specific combinations.

*Leave-Combination-Out Cross-Validation*. Finally, a leave-combination-out experiment was performed to emulate a stringent OOD condition. In this setting, all drug pairs with moderate-to-high similarity to training pairs (Tanimoto ≥ 0.5 on physicochemical descriptors and cosine similarity ≥ 0.6 on SMILES embeddings) were excluded from training. As shown in **Figure 4c**, the model maintained strong predictive power across all synergy metrics, particularly for ZIP (PCC = 0.80, MAE = 5.4, R² = 0.64), while continuing to perform robustly across the full spectrum of synergy values, from antagonistic to highly synergistic interactions.

Together, these analyses demonstrate that AlgoraeOS exhibits strong generalisation beyond its training distribution, with reliable few-shot and zero-shot learning capability and sustained performance under stringent OOD evaluation.

### Uncertainty Quantification and Reliability of Predictions

Reliable uncertainty quantification (UQ) is essential for building confidence in AI models applied to drug discovery, ensuring that predictions are not only accurate but also trustworthy when guiding experimental decisions ^37,38^. AlgoraeOS implements an explicit framework for estimating prediction confidence by separating two complementary sources of uncertainty: aleatoric (data-related) and epistemic (model-related).

The aleatoric uncertainty ^28^ captures variability inherent to the experimental data—such as assay noise, biological heterogeneity, and variability between replicates. This is estimated through a dedicated network head that learns the covariance among the four synergy metrics by minimising a multivariate Gaussian negative log-likelihood. This setup allows the model to recognise regions where the data itself is noisy or ambiguous without affecting the main prediction pathway.

The epistemic uncertainty reflects the model’s uncertainty due to limited or biased training data, such as unseen drug classes or rare dosage conditions. It is estimated using Monte-Carlo Dropout ^37^, where multiple stochastic forward passes through the network generate a distribution of predictions. The variability of these predictions reflects how confident the model is in a given region of the input space.

The total predictive uncertainty is computed as the sum of the aleatoric and epistemic estimates, providing a full predictive distribution for every drug–cell–dose combination. **Figure 5a** illustrates the relationship between prediction error and model uncertainty across the full range of Bliss synergy predictions. Each coloured cell represents the average predicted uncertainty for samples within a given range of Bliss values (x-axis) and absolute errors (y-axis). Within each prediction bin, higher absolute errors coincide with warmer colours (yellow– red), indicating greater mean uncertainty, whereas regions of minimal error remain consistently green, reflecting low uncertainty. **Figure 5b** quantifies this relationship across prediction bins, reporting Pearson r, Spearman ρ, and Kendall τ correlations between uncertainty and absolute error. The correlations are predominantly positive across bins, indicating that higher uncertainty accompanies higher error. This strong error–uncertainty concordance demonstrates that AlgoraeOS not only produces accurate predictions but also assigns confidence appropriately, i.e., being most certain where its predictions are correct and expressing caution where deviations increase. This alignment between uncertainty magnitude and predictive error reflects reliable uncertainty quantification consistent with best practices in AI-driven drug discovery and molecular modelling ^37,39^.

**Figure 5.**
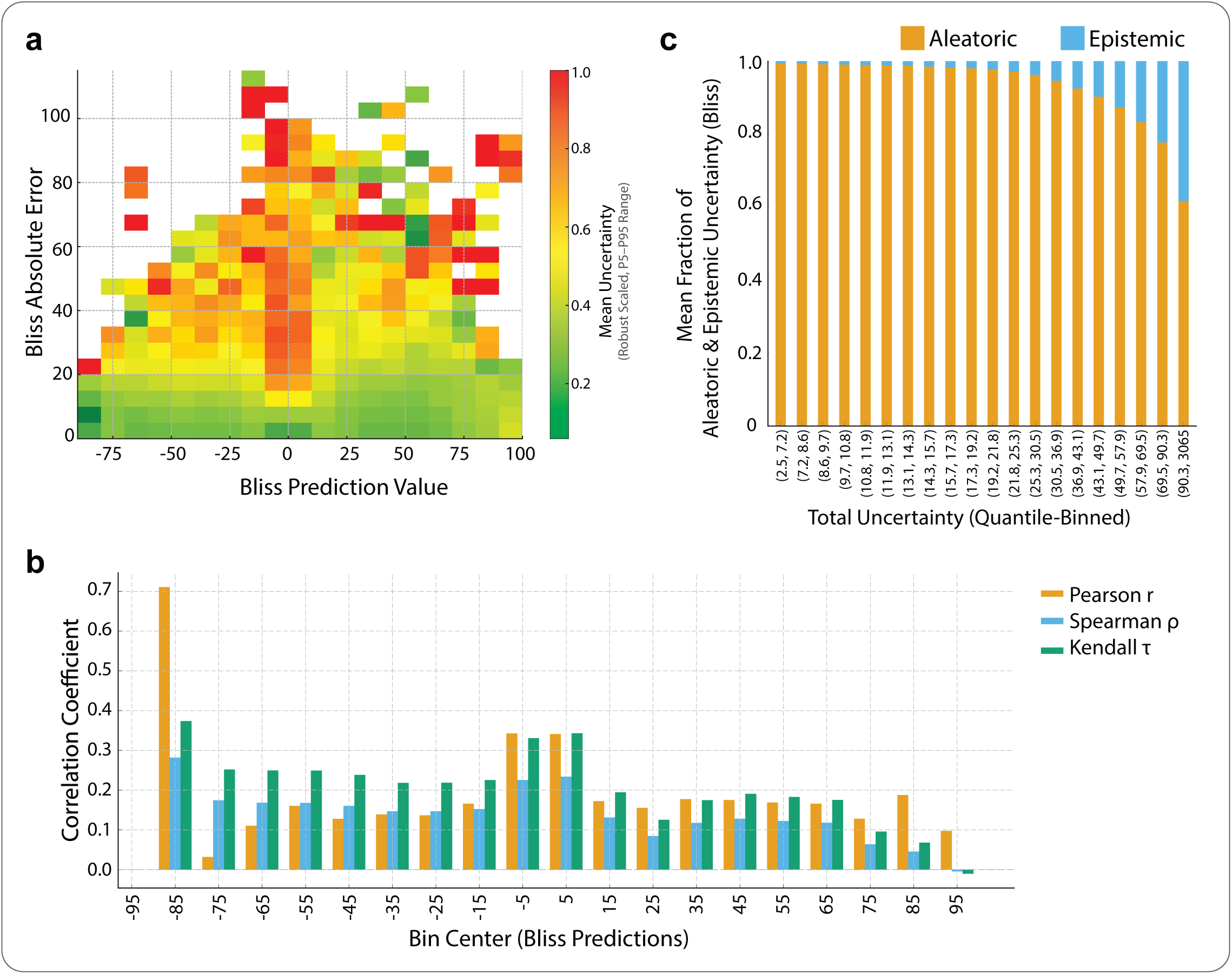
Uncertainty quantification and decomposition in AlgoraeOS. **(a**) Relationship between prediction error and model uncertainty across the full range of ZIP synergy predictions. Each coloured cell represents the average predicted uncertainty for samples within a given range of predicted ZIP values (x-axis) and absolute error (y-axis). **(b)** Correlation between model uncertainty and absolute prediction error across ZIP bins, shown for Pearson *r*, Spearman ρ, and Kendall τ coefficients. **(c)** Decomposition of total predictive uncertainty into aleatoric (data-dependent) and epistemic (model-dependent) components, expressed as mean fractional contributions across quantile-binned total uncertainty levels.

**Figure 5c** breaks down total predictive uncertainty into its aleatoric and epistemic components. The x-axis shows quantile-binned total uncertainty, while the y-axis indicates the mean fractional contribution of each component. At low total uncertainty levels, nearly all uncertainty arises from aleatoric sources, reflecting stable, well-sampled experimental regimes. As total uncertainty increases, the epistemic share (blue) expands progressively, indicating that model uncertainty dominates in regions with limited data coverage, such as rare chemotypes, extreme doses, or underrepresented tissue contexts. This gradient confirms that AlgoraeOS correctly attributes uncertainty to data noise versus model ignorance across the prediction spectrum, reinforcing its interpretability and reliability for uncertainty-aware decision-making.

### Benchmarking Against Existing and Baseline Models

To contextualise the performance of AlgoraeOS within the broader landscape of deep learning models for drug synergy prediction, we conducted a comprehensive literature review identifying 12 representative architectures developed between 2018 and 2025 (**Supplementary File 1**). The earliest of these was *DeepDDS* (2018), a pioneering multi-modal deep neural network for synergy regression, and the most recent is *TxGemma* (2025), a large language model (LLM)-based therapeutic discovery framework from Google DeepMind, which extends to drug combination prediction.

Most existing models were trained or fine-tuned on subsets of the DrugComb V1.5 dataset (typically smaller panels such as NCI-ALMANAC, O’Neil, or Merck) with only four models utilising datasets approaching the scale and diversity of ours (ranging from approximately 157K to 286K drug–cell–drug records). To enable fair comparison, AlgoraeOS was retrained on matching subsets (e.g., ALMANAC and O’Neil) and evaluated using the same synergy metrics as those reported in the respective publications. When model data splits, or training protocols were unavailable or ambiguous (e.g., *MARSY*), those models were excluded from direct benchmarking.

**Table 2** summarises this comparative analysis, listing one representative or best-performing version of each model. Across all datasets and synergy formalisms (Bliss, HSA, Loewe, ZIP), AlgoraeOS consistently outperformed previously published architectures, achieving markedly lower absolute errors (MAE) and higher correlation coefficients (PCC). The greatest performance margins were observed on datasets exhibiting higher biological heterogeneity, such as **DrugComb** and **O’Neil**, highlighting the model’s improved generalisability.

**Table 2.** Summary of major deep learning models developed for drug-synergy regression prediction, alongside corresponding performance of AlgoraeOS on equivalent datasets. Metrics include mean absolute error (MAE), mean squared error (MSE), root mean squared error (RMSE), Pearson correlation coefficient (PCC), and coefficient of determination (R²). For DeepDDS, metrics were extracted from the MultiComb paper since the original training dataset was not clearly specified. For models with multiple configurations or variants, only the best-performing version is reported here. Full details and additional models are provided in **Supplementary File 1**.

| Model (Year) | Dataset | Evaluation Strategy | Reported Performance | AlgoraeOS (Equivalent Dataset) |
| --- | --- | --- | --- | --- |
| **DeepSynergy (2018)** <sup>16</sup> | O'Neil / Merck | 5-fold CV | **Loewe:** MSE = 255.49, RMSE = 15.91, PCC = 0.73 | **Loewe:** MAE = 3.47, RMSE = 4.75, PCC = 0.953 |
| **AuDNNsynergy (2021)** <sup>43</sup> | O'Neil / Merck | 5-fold CV | **Loewe:** MSE = 241.12, RMSE = 15.46, PCC = 0.74 | **Loewe:** MAE = 3.47, MSE = 22.52, RMSE = 4.75, PCC = 0.953 |
| **TranSynergy (2021)** <sup>44</sup> | O'Neil / Merck | 5-fold CV | **Loewe:** MSE = 231, PCC = 0.75 | **Loewe:** MSE = 22.52, PCC = 0.953 |
| **PRODeepSyn (2022)** <sup>45</sup> | O'Neil / Merck | 5-fold CV | **Loewe:** MSE = 229.49, RMSE = 15.09, PCC = 0.75 | **Loewe:** MAE = 3.47, RMSE = 4.75, PCC = 0.953 |
| **Synpathy (2022)** <sup>46</sup> | O'Neil / Merck | Repeated 5-fold CV | **Loewe:** MSE = 64.7, RMSE ≈ 8.0 | **Loewe:** MSE = 22.52, RMSE = 4.75 |
| **MatchMaker (2022)** <sup>47</sup> | DrugComb | 60 / 20 / 20 split + 5-fold CV | **Loewe:** MSE = 79.49, PCC = 0.79 | **Loewe:** MSE = 30.43, PCC = 0.853 |
| **SYNPRED Ensemble (2022)** <sup>48</sup> | NCI ALMANAC | 80 % train / 20 % test | **Bliss:** MAE = 3.07, RMSE = 4.35, PCC = 0.71<br>**HSA:** MAE = 2.86, RMSE = 4.09, PCC = 0.73<br>**Loewe:** MAE = 6.49, RMSE = 10.58, PCC = 0.71<br>**ZIP:** MAE = 2.74, RMSE = 3.86, PCC = 0.70 | **Bliss:** MAE = 3.22, RMSE = 4.42, PCC = 0.675<br>**HSA:** MAE = 3.00, RMSE = 4.13, PCC = 0.697<br>**Loewe:** MAE = 6.46, RMSE = 9.74, PCC = 0.744<br>**ZIP:** MAE = 2.89, RMSE = 3.93, PCC = 0.666 |
| **DeepDDS (2024)\*** <sup>15</sup> | O'Neil / Merck | 5-fold CV | **Loewe:** MAE = 9.98, MSE = 248.20, RMSE = 15.69, $R^2$ = 0.54, PCC = 0.74 | **Loewe:** MAE = 3.47, MSE = 22.52, RMSE = 4.75, PCC = 0.953, $R^2$ = 0.908 |
| **MultiComb (2024)** <sup>49</sup> | O'Neil / Merck | 5-fold CV | **Loewe:** MAE = 9.59, RMSE = 15.19, PCC = 0.76, $R^2$ = 0.57 | **Loewe:** MAE = 3.47, RMSE = 4.75, PCC = 0.953, $R^2$ = 0.908 |
| **Gemma-2-27B (Base, 2025)** <sup>50</sup> | NCI ALMANAC | 70 / 10 / 20 split | **Bliss:** MAE = 6.46; **HSA:** MAE = 6.67; **Loewe:** MAE = 14.73; **ZIP:** MAE = 5.40 | **Bliss:** MAE = 3.22; **HSA:** MAE = 3.00; **Loewe:** MAE = 6.46; **ZIP:** MAE = 2.89 |
| **Tx-LLM M (2025)** <sup>51</sup> | NCI ALMANAC | 70 / 10 / 20 split | **Bliss:** MAE = 4.10; **HSA:** MAE = 4.12; **Loewe:** MAE = 17.38; **ZIP:** MAE = 3.78 | **Bliss:** MAE = 3.22; **HSA:** MAE = 3.00; **Loewe:** MAE = 6.46; **ZIP:** MAE = 2.89 |
| **TxGemma-27B-Predict (2025)** <sup>50</sup> | NCI ALMANAC | 70 / 10 / 20 split | **Bliss:** MAE = 4.16; **HSA:** MAE = 4.21; **Loewe:** MAE = 17.34; **ZIP:** MAE = 3.81 | **Bliss:** MAE = 3.22; **HSA:** MAE = 3.00; **Loewe:** MAE = 6.46; **ZIP:** MAE = 2.89 |

\* Metrics from MultiComb paper

Recognising that deep learning models do not always outperform traditional regression methods in biomedical prediction tasks ^40^, we next benchmarked AlgoraeOS against a suite of widely used baseline models designed to capture both linear and non-linear structure. These included a Multi-Layer Perceptron (MLP), Random Forest, Linear Regression, and Ridge Regression, collectively spanning a representative spectrum from non-parametric ensemble learners to linear parametric estimators.

All baseline models were trained and evaluated under identical conditions on the DrugComb dataset to ensure comparability. Across all synergy scores and performance metrics, AlgoraeOS consistently achieved the strongest results. Among baselines, the MLP performed best (reflecting the inherently non-linear nature of drug–response relationships), achieving, for example, on Bliss regression, MAE = 5.05, RMSE = 7.61, and PCC = 0.913. Relative to this strongest baseline, AlgoraeOS achieved 6.25% lower MAE and 6.08% lower RMSE.

However, aggregate metrics can obscure subtle but clinically relevant error structures. To probe these effects, we visualized column-normalized Observed versus Predicted and Residual (Observed – Predicted error) heatmaps (**Supplementary Figures 4–7**). Despite competitive overall performance, the MLP displayed *(i)* heteroscedasticity, with widening error variance at higher/lower Bliss values; *(ii)* systematic underestimation in strongly synergistic (positive) regimes; and *(iii)* asymmetric residual tails, suggesting calibration drift not captured by MAE or RMSE. In contrast, AlgoraeOS exhibited tight alignment along the identity line and a narrow, symmetric residual band centred around zero, signifying uniform calibration and homoscedastic error across the full synergy spectrum.

Overall, the combined benchmarking against both state-of-the-art deep learning architectures and classical regression baselines establishes AlgoraeOS as a robust, well-calibrated, and generalizable framework that not only surpasses prior models in predictive accuracy but also delivers greater reliability across biologically diverse and clinically relevant synergy regions.

### Dose Sensitivity and Inhibition Rate Prediction

To capture the dynamic, dose-dependent nature of drug interactions, we extended the AlgoraeOS architecture to operate on non-aggregated, dose-specific data. This involved training a new model on a vastly expanded dataset of 5.5 million unique inhibition records from DrugComb (11 million records after reverse augmentation). This concentration-aware model introduces two key architectural innovations: **(i)** the addition of inhibition as a fifth prediction head, explicitly linking potency to synergy, and **(ii)** the integration of a Concentration Encoder and Feature-wise Linear Modulation (FiLM) ^41^ layers to make the model dose-aware.

The Concentration Encoder transforms raw scalar drug dosages into rich, functional embeddings. Instead of feeding log-transformed concentrations directly to the network, the encoder projects them into a high-dimensional feature space using a hybrid basis of Fourier, radial-basis (RBF), and polynomial functions. This learned projection effectively linearises complex, non-linear dose-response relationships, enabling the network to model pharmacodynamic surfaces without being constrained by predefined curve-fitting models like the Hill equation. These rich dose embeddings are then used by FiLM layers to modulate the drug feature representations before they enter the cross-attention module. This mechanism allows the model to understand not just the identity of a drug, but its relative potency and position on its pharmacodynamic trajectory in a given experiment.

**Figure 6a** presents the error distribution across inhibition bins, with experimental inhibition values divided into ten deciles. A stringent admissible error range was defined as ± (0.1 × |observed inhibition| + 5), represented by the dashed red lines, corresponding to 10 % of the measured inhibition plus a 5 % tolerance. Across all bins, the violin plots remain tightly centred around zero, with the majority of predictions falling well within this admissible range and a median absolute error of approximately 3–4 %. The uniform error variance across bins confirms that explicit concentration encoding mitigates response-surface aliasing and yields consistent calibration across the entire pharmacodynamic spectrum.

**Figure 6.**
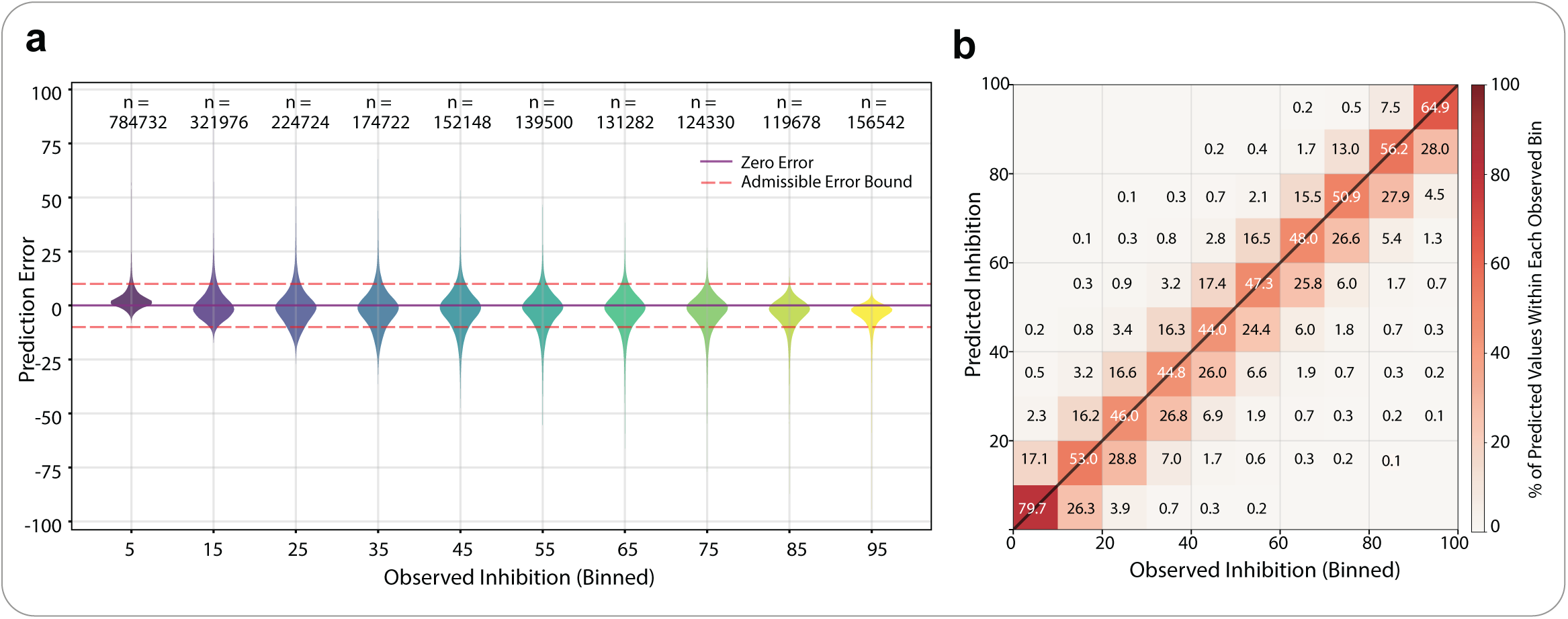
Evaluation of dose-specific inhibition prediction performance. **(a)** Error distribution across inhibition bins. Inhibition values were divided into ten deciles to assess calibration as a function of effect strength. Violin plots show prediction errors distribution. The dashed red lines indicate an admissible error range defined as ±(0.1 × |observed inhibition| + 5). **(b)** Heatmap displaying the joint density of observed versus predicted inhibition values, binned into ten intervals based on observed inhibition. Density concentration along the y = x line indicates overall prediction concordance.

**Figure 6b** compares ground-truth versus predicted inhibition values across all test folds. The dense diagonal ridge along y = x confirms near-perfect correspondence between measured and predicted inhibition, evidencing excellent calibration across the full dynamic range. Statistical metrics (PCC = 0.922, Concordance Correlation Coefficient = 0.915) confirm strong rank-order fidelity and absolute-scale calibration. Even at the extreme ends of the spectrum (≤5% and ≥95% inhibition), prediction contours remain sharp, showing that the model correctly captures both weak and strong inhibitory effects. A modest broadening in the midrange (40– 60%) corresponds to biological heterogeneity, where replicate variance is highest in the DrugComb dataset.

Overall, the concentration-aware framework converts simple dosage inputs into rich, biologically meaningful representations, allowing the model to accurately capture complex dose–response behaviour. This leads to sharper calibration, reduced bias across inhibition ranges, and improved biological relevance, establishing a robust foundation for dose-aware drug-synergy prediction.

### Prospective Experimental Validation Using *In Vitro* Combination Screening

To assess model performance in a prospective setting, we conducted an independent *in vitro* validation using Cannabidiol (DB09061) as an anchor compound, for which no prior synergy measurements were present in the training corpus (∼5.8 million dose–combination records), enabling an out-of-distribution evaluation. Screening was performed across three molecularly distinct cancer cell lines representing glioblastoma multiforme, castration-resistant prostate cancer, and triple-negative breast cancer (T98G, 22Rv1, and BT-20). Cannabidiol was tested in combination with 24 partner compounds spanning diverse mechanistic classes (**Supplementary File 3**). The degree of prior exposure to these partner compounds in the training data differed across cell lines: no matching partner drugs were present for 22Rv1 (0/24; 0 records), while limited overlap was observed for BT-20 (2/24; 398 records) and T98G (14/24; 37 records), reflecting a spectrum from complete novelty to sparse prior exposure. Partner compounds were selected to reflect pharmacological diversity and clinical relevance; compound identities are withheld for commercial sensitivity but are available upon reasonable request. Combinations were evaluated using a full 6 × 6 matrix of non-zero concentrations (36 dose pairs per combination), yielding 2,592 dose–combination measurements (24 drugs × 3 cell lines × 36 dose pairs). All experiments were conducted independently of model training without retrospective calibration, and detailed results at the compound–cell line–dose level, including inhibition measurements, are provided in **Supplementary File 3.**

Across the three cell lines, AlgoraeOS demonstrated consistent rank-order fidelity between predicted and observed inhibition values despite the anchor compound being entirely absent from training. Spearman’s rank correlation was ρ = 0.59 for the prostate cancer line (22Rv1), ρ = 0.51 for the breast cancer line (BT-20), and ρ = 0.52 for the glioblastoma line (T98G) (all p-values < 0.001), indicating that the model reliably preserves the relative ordering of combination responses within each biological context (**Figure 7a**). To assess decision-level utility, we applied a concordance–discordance framework using a 10% inhibition threshold with a ±5% boundary zone (**Figure 7b**), classifying predictions as concordant, discordant, or borderline. The boundary zone captures values near the decision threshold where small fluctuations can change classification without reflecting meaningful biological differences. Of 2,592 dose combinations, 1,857 (71.6%) were concordant, 137 (5.3%) discordant, and 598 (23.1%) fell within the boundary zone. Directional agreement was quantified using Kendall’s τₐ = (C - D)/(C + D), where C and D denote the number of concordant and discordant pairs, respectively; this reached 0.863 overall, demonstrating strong alignment between predicted and observed effects when predictions fall outside the uncertainty region. This pattern was consistent across cell lines, with τₐ = 0.915 (glioblastoma; C = 565, D = 25), 0.785 (prostate; C = 631, D = 76), and 0.897 (breast; C = 661, D = 36). These results show that, even under out-of-distribution conditions, AlgoraeOS yields reliable directional predictions for prioritising candidate combinations.

**Figure 7.**
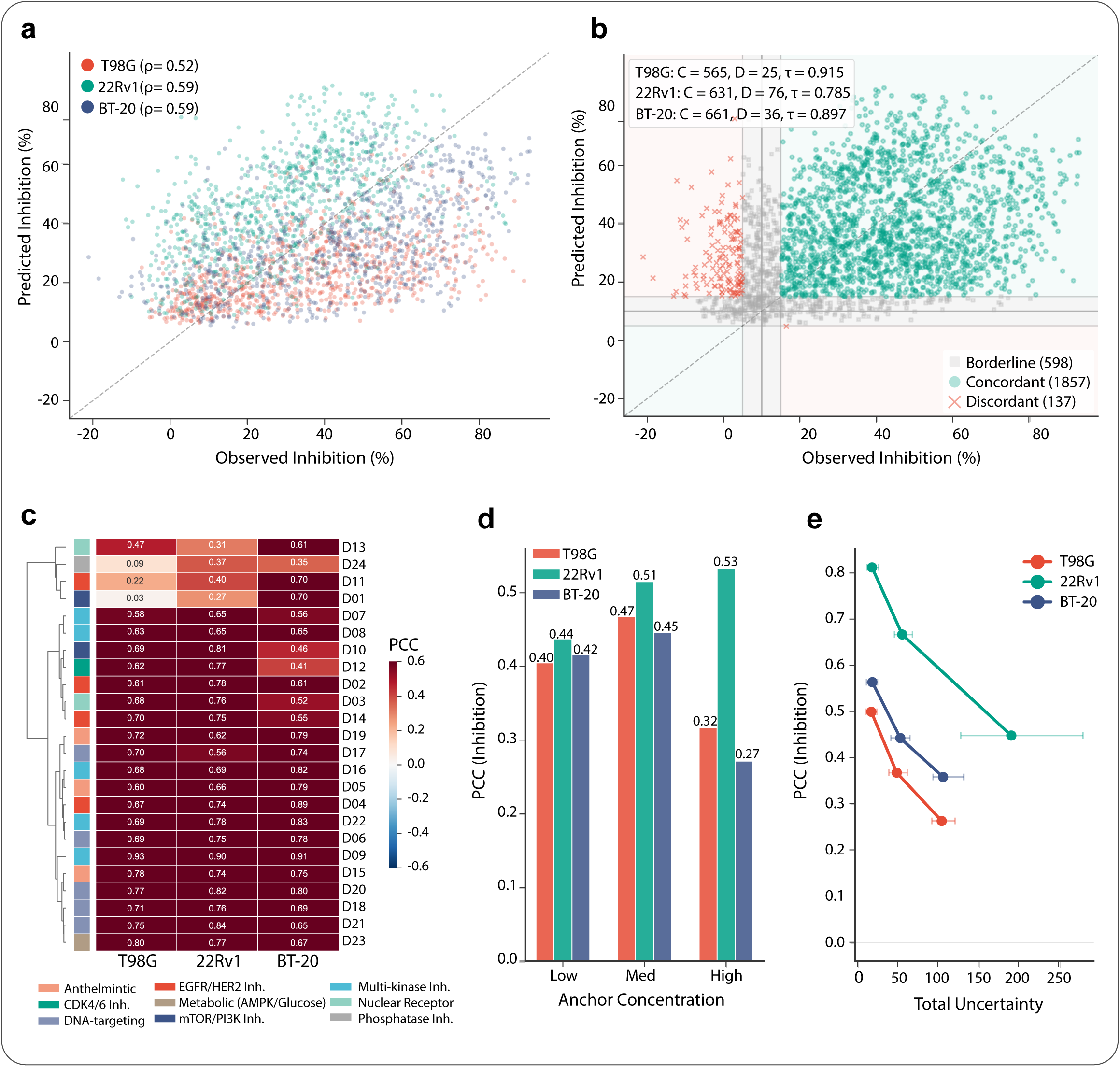
Prospective *in vitro* validation of AlgoraeOS using Cannabidiol as an anchor compound across three cancer cell lines (T98G, 22Rv1, BT-20). **(a)** Predicted versus observed inhibition (%) for all dose–combination measurements, coloured by cell line, demonstrating rank-order agreement across biological contexts. Spearman correlation values are shown for each cell line and overall. **(b)** Concordance– discordance analysis using a 10% inhibition threshold with a ±5% boundary zone. Shaded regions denote the boundary zone (light grey), concordant regions (green), and discordant regions (red). Counts of concordant (C), discordant (D), and borderline predictions are reported, with directional agreement quantified using Kendall’s τₐ = (C - D)/(C + D). **(c)** Dose-aggregated Pearson correlation coefficient (PCC) for each drug–cell line combination, with partner drugs ordered by hierarchical clustering. Side colour bars denote pharmacological classes, illustrating performance consistency across mechanistically diverse drug categories. **(d)** PCC between predicted and observed inhibition stratified by anchor (Cannabidiol) concentration level (low: 1.0–3.0 µM; medium: 5.0–7.0 µM; high: 9.0–15.0 µM), showing stable performance across concentrations with modest reductions at higher doses. **(e)** Relationship between model-estimated total uncertainty and prediction accuracy, with observations pooled and stratified into three equal-frequency uncertainty bins (terciles). Points represent PCC values per bin and cell line, with horizontal bars indicating interquartile ranges of uncertainty values, showing reduced predictive accuracy with increasing uncertainty.

Dose-aggregated Pearson correlation at the individual combination level (**Figure 7c**) further indicates that predictive performance is not driven by prior exposure of specific partner drugs within a given cell line. In T98G, combinations involving partner drugs present in the training set did not outperform those entirely unseen, with no significant difference in performance (Mann–Whitney test p = 0.66; BT-20 p = 0.59). Instead, predictive performance was more strongly associated with the pharmacological class of the partner compound, with mechanisms well represented in the global training corpus (e.g., multi-kinase inhibitors and metabolic modulators) achieving consistently higher concordance across cell lines. This suggests that AlgoraeOS learns transferable representations of underlying drug mechanisms rather than relying on cell line–specific exposure to individual compounds.

Across concentration levels, predictive performance was evaluated by stratifying each experiment into low (1.0, 3.0 µM), medium (5.0, 7.0 µM), and high (9.0, 10.0, 12.0, 15.0 µM) anchor (Cannabidiol) concentration groups. Spearman correlation between predicted and observed inhibition values (**Figure 7d**) remained relatively stable across concentrations for 22Rv1, indicating consistent performance despite complete absence of training exposure. T98G and BT-20 showed reduced concordance at higher concentrations. This pattern likely reflects increased biological and experimental variability at high dose levels, including saturation effects, non-linear dose–response behaviour, and reduced dynamic range, which collectively make accurate prediction more challenging. Similar trends were observed when stratifying by partner drug concentration (**Supplementary Figure 8**).

The relationship between prediction accuracy and model-estimated total uncertainty is shown in **Figure 7e**. All 2,592 observations were pooled and stratified into three equal-frequency uncertainty bins (low, medium, high), yielding balanced sample sizes given the right-skewed distribution of uncertainty values. For each cell line and bin, Spearman correlation between predicted and observed inhibition was computed, with the median and interquartile range indicating the spread of uncertainty values contributing to each estimate. Across all cell lines, lower-uncertainty bins consistently showed higher correlation than higher-uncertainty bins. This trend was most pronounced in 22Rv1, where correlation decreased from approximately 0.81 in the lowest-uncertainty bin to ∼0.45 in the highest, and was also observed in T98G and BT-20. These results indicate that the model’s uncertainty estimates provide a meaningful signal for stratifying predictions by expected reliability.

## Conclusion

In this study, we developed AlgoraeOS, a biologically-informed, attention-aware deep learning platform designed to address the critical limitations of contemporary drug synergy prediction models. By moving beyond single, dose-averaged metrics, our framework provides a holistic, uncertainty-aware, and dose-resolved assessment of drug combination efficacy. We have demonstrated that AlgoraeOS not only achieves state-of-the-art predictive accuracy but also establishes a new, more rigorous standard for model development and validation in the field.

Our findings yield several key insights for the future of computational synergy prediction. First, we have shown that relying on single, aggregated error metrics is insufficient and can be misleading. While AlgoraeOS outperforms baseline models on metrics like MAE and PCC, our deeper analysis of residual distributions reveals that it also provides superior calibration across the full synergy spectrum, a critical feature that aggregated scores obscure. Second, our multi-task learning architecture, which simultaneously predicts four distinct synergy scores, demonstrates the importance of preserving the empirical correlations between metrics. This ensures the model learns a more robust and nuanced representation of drug interactions rather than overfitting to a single mathematical definition of synergy. Third, and perhaps most crucially, we underscore the absolute necessity of rigorous out-of-distribution validation. Our zero-shot and few-shot learning assessments confirm that AlgoraeOS can successfully generalise to unseen drugs and cell lines, providing a level of translational confidence that is often missing from models validated only with standard cross-validation.

Critically, prospective in vitro validation using an anchor compound entirely absent from the training corpus confirmed that these computational gains translate to experimentally meaningful predictions, with consistent rank-order fidelity observed across three molecularly distinct cancer cell lines and 2,592 dose–combination measurements. The further finding that model-estimated uncertainty stratified predictions by their experimental concordance underscores the practical utility of uncertainty-aware frameworks in guiding and prioritising resource-intensive wet-lab validation.

Finally, while AlgoraeOS was trained on cancer cell line data, its fundamental architecture is designed to model any high-throughput screen (HTS) with a cellular inhibition or proliferation readout. This suggests a strong potential for its application to other complex diseases where combination therapies are paramount, such as infectious diseases, immunology, and fibrosis; however, validating this broader applicability will require dedicated future assessments on disease-specific datasets. It is also important to acknowledge that the performance of any data-driven model, including AlgoraeOS, is ultimately constrained by the scope of the training data. Our own exploratory analysis highlighted that the current HTS landscape is highly uneven, with a small subset of drugs being extensively profiled while the vast majority of the combinatorial space remains unexplored. To overcome this, future progress should focus on more systematic experimental designs. Integrating predictive models like AlgoraeOS into an *active learning* ^42^framework represents a particularly promising path forward. In such a closed-loop system, the model’s predictions and its own uncertainty estimates could guide the selection of the most informative new experiments to perform, enabling a more efficient and targeted exploration of the vast combinatorial space.

## Materials and Methods

### Data Preprocessing

We curated data from DrugComb v1.5—one of the field’s most comprehensive resources, which compiles many combination-screen datasets, harmonises assay metadata and dose– response formats, and standardises synergy scores via SynergyFinder—where each experiment is a (drug A, drug B, cell line) block over a concentration grid. We retained only dual therapies (excluding drug₁ = drug₂) on cell lines with CCLE DepMap transcriptomes; removed entries with SMILES inconsistencies, missing synergy, or scores outside [−100, 100]; and resolved duplicate triplets by keeping the within-block median. To enforce order-agnostic modelling, we applied on-the-fly drug-swap augmentation (drug₁↔drug₂), yielding 736,246 augmented records. CCLE DepMap RNA-seq TPMs were aligned to a unified gene universe and aggregated into KEGG pathway activations with length normalisation. Drug features combined RDKit physicochemical descriptors^24^ with ChemBERTa SMILES embeddings^25^; available concentrations were log-transformed with a small positive floor. For inhibition analyses, values were constrained to [0, 100] and processed with the same alignment and quality-control steps, yielding ∼5.5 million assay-level inhibition records.

### Data Splits and Evaluation Design

We designed three complementary evaluation regimes to assess in-distribution generalisation, cross-study transfer, and similarity-aware out-of-distribution (OOD) robustness.

1. *Main Dataset:* The core dataset was derived from DrugComb and comprises approximately 736,000 unique drug–drug–cell line triplets.
2. *Cross-Study External Evaluation:* To enable fair comparison with prior work, we partitioned the full DrugComb dataset by source study (e.g., O’Neil/Merck, NCI-ALMANAC) and evaluated each subset independently without cross-study leakage. This design yields study-specific performance estimates suitable for benchmarking against existing study-level baselines.
3. *Stratified Subset for Efficient Experimentation:* The 136K subset was stratified to preserve the balance and difficulty of the prediction tasks. ZIP and Bliss scores were discretised into empirical quintiles, and samples were drawn to proportionally represent each combination of these bins. A 120,000/16,000 training/validation split was then created using the same stratification strategy to maintain consistent label distributions.
4. *Similarity-Aware Leave-Pair-Out (OOD) Evaluation:* To rigorously assess model robustness, we implemented a similarity-aware leave-pair-out scheme that withholds entire drug-pair neighbourhoods from training. Pairwise drug–drug similarity was computed using RDKit with Jaccard (Tanimoto) similarity and ChemBERTa embeddings with cosine similarity, with each score scaled to [0, 1]. Two drugs were deemed similar if 0.6 ≤ Tanimoto ≤ 0.7 and 0.7 ≤ cosine similarity ≤ 0.8. we evaluated both bijective matchings (*d*_1_, *d*_2_) and (*d*_3_, *d*_4_); a matching was accepted only if each mapped drug pair met the similarity criterion and each drug in one pair mapped to a distinct drug in the other (i.e. we did not allow both *d*_1_ and *d*_2_ to be similar to the same counterpart). Two pairs were labelled “similar” if at least one such matching existed. For each anchor drug pair, its neighbourhood was defined as all pairs similar to it, and all dose–response triplets whose drug pair fell within any anchor neighbourhood were withheld and assigned to the OOD test set, ensuring that neither anchor pairs nor their close analogues appeared in training.

This approach avoids both trivially easy (near-duplicate) and unrealistically difficult (completely dissimilar) cases, targeting a moderate OOD zone that best reflects real-world generalisation performance.

### Problem Formulation and Learning Objective

We modelled combination prediction as supervised, probabilistic, multi-task regression over drug–cell contexts. Let each training instance *i* ∈ {1, …, *N*} be an assay triplet with inputs:

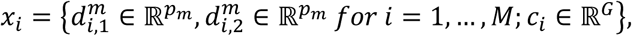

where (*m*) indexes drug feature modalities (e.g., RDKit, ChemBERTa) and (*c*_*i*_) is the cell-line gene-expression vector. The target is the 4-dimensional synergy vector:

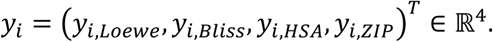

The predictive model parameterised by *w* defines a conditional multivariate normal distribution, *p*_*w*_(*y*|*x*) = *p*(*y*; *μ*_*w*_(*x*), Σ_*w*_(*x*)), where *μ*_*w*_(*x*): *χ* → ℝ^4^ is produced by the four synergy heads and the covariance Σ_*w*_(*x*) ∈ ℝ^4×4^ is produced by an uncertainty head. We decompose covariance into aleatoric and epistemic components:

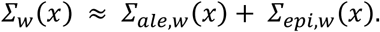

Aleatoric uncertainty is predicted by a dedicated head operating on detached fused features and parameterised as low-rank-plus-diagonal:

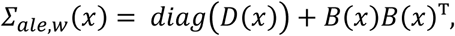

with 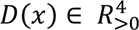 (enforced via soft-plus) *B*(*x*) ∈ ℝ^4×*r*^, *r* ≪ 4, (*here r* = 2). We use Cholesky-based computations with a small diagonal jitter to keep the covariance symmetric positive definite (SPD; 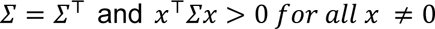), which is required for a valid Gaussian likelihood, invertibility, and stable gradients.

Epistemic uncertainty is estimated at inference via Monte-Carlo Dropout. Drawing *T* stochastic forward passes 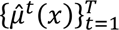 over the predictive heads, 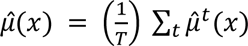and 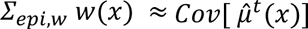.

Hence, the model yields a calibrated 4-variate predictive PDF whose mean encodes synergy estimates and whose covariance captures both data noise (aleatoric) and model uncertainty (epistemic).

### Loss, Masking and Curriculum

For sample *i*, let *T*_T_ ⊆ (*Loewe*, *Bliss*, *HSA*, *ZIP*) denote the set of available synergy metrics. We minimise a phase-scheduled empirical risk that first sharpens means and then learns covariance:

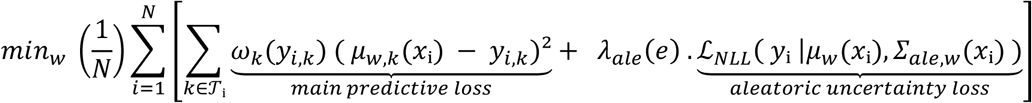

where *e* is the epoch. The per-task weights 𝜔_𝑘_(·) upweights rare/extreme synergies via inverse-density weighting computed from a smoothed empirical density

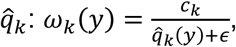

with *c*_𝑘_ a normalisation constant; task-specific head scales are registered as non-trainable buffers. The curriculum 𝜆_𝑎𝑙*e*_(*e*) is piecewise constant: 𝜆_𝑎𝑙*e*_(*e*) = 0 during an initial MSE-only phase, then 𝜆_𝑎𝑙*e*_(*e*) = _0_ > 0 after a threshold epoch to activate covariance learning on residuals. Optimisation uses separate parameter groups for the predictor vs. aleatoric head and gradient clipping for stability.

The multivariate Gaussian negative log-likelihood minimised by the aleatoric head is:

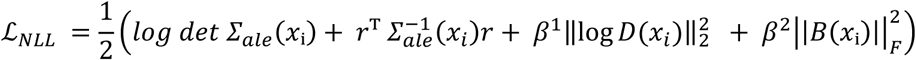

with residual *r*_*i*_ = (*y*_T_ - *μ*_*w*_ (*x*_*i*_)) and mild regularisation (𝛽_1_, 𝛽_2_) for variance and low-rank factors.

### Test-Time Predictive Distribution

At test time, we report the full predictive distribution: 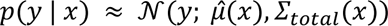, 𝛴_𝑡𝑜𝑡𝑎𝑙_(*x*) = 𝛴_𝑎𝑙*e*_(*x*) + 𝛴_*epi*_(*x*), from which calibrated marginal standard deviations 𝜎_𝑡𝑜𝑡,𝑘_(*x*), off-diagonal covariances, and joint prediction intervals are obtained for the four synergy measures. Putting the above together, learning solves a regularised conditional NLL with masking and curriculum:

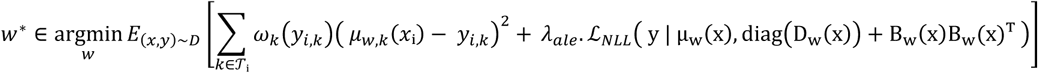

### Architecture

To predict drug combination synergy, we developed AlgoraeOS, a hybrid early-to-mid-fusion architecture that integrates multimodal drug features and cell-line transcriptomics (Fig. 1). The model leverages cross-modal attention to learn context-dependent interactions and projects the integrated representations to multi-task regression heads that jointly predict four synergy metrics: Loewe, Bliss, HSA, and ZIP. Crucially, the framework explicitly quantifies aleatoric (data-dependent) and epistemic (model-dependent) uncertainty.

The model architecture comprises five key modules:

#### 1. Modality-Specific Drug Embedders

For each of the two drugs in a combination, heterogeneous feature modalities are first processed by separate modality-specific embedders. Each embedder, implemented as a two-layer MLP with Layer Normalisation, ReLU activation, and dropout, projects its input modality into a shared latent space 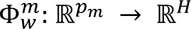, where *p*_m_is the dimension of modality *m* and 𝐻 = 256 is the hidden dimension. This initial embedding normalises feature scales and distributions, creating stable unimodal representations. The resulting embeddings for each drug are stacked into tensors *D*_1_,*D*_2_ ∈ {ℝ^*M_d_*×*H*^}, where 𝑀_*d*_ is the number of drug modalities.

#### 2. Cell-Line Pathway Encoder

The cell-line gene expression profile 𝑋 ∈ ℝ^𝐺^ (where G=16,720 genes) is transformed into a compact, biologically structured embedding, 𝜓_*w*_ℝ^𝐺^ → ℝ^𝐻^. This is achieved via a two-stage process. First, a fixed, non-trainable linear transformation aggregates gene-level signals into pathway activations. To prevent bias from pathway size, each pathway activation is normalised by the number of genes it contains. The resulting pathway profile is passed through a Soft-plus activation to ensure non-negativity and then projected by a trainable MLP into a final cell-context vector 𝐶 ∈ ℝ^𝐻^. This approach introduces a strong biological prior, enhances generalizability, and produces a dense representation of the cell’s functional state.

#### 3. Cross-Modal Attention Hub

The core of AlgoraeOS is a multi-head attention module (8 heads) that facilitates interaction between the two drug representations (𝐷_1_, 𝐷_2_) and the cellular context (𝐶). Three distinct attention operations are applied:

- **Drug→Cell:** Each drug embedding queries the cell context to learn cell-specific drug response features: Attn(𝑄 = 𝐷_*i*_, 𝐾 = 𝐶, 𝑉 = 𝐶) for *i* ∈ {1,2}
- **Cell→Drug:** The cell embedding queries each drug’s modalities to incorporate drug-conditioned cellular effects: Attn(𝑄 = 𝐶, 𝐾 = 𝐷_*i*_, 𝑉 = 𝐷_*i*_) for *i* ∈ {1,2}
- **Drug↔Drug:** Bidirectional attention between 𝐷_1_ and 𝐷_2_ models synergistic or antagonistic chemical interactions.

Each attention block is followed by residual connections and Layer Normalisation. This hub generates context-aware representations that capture non-additive interactions essential for synergy prediction.

#### 4. Feature Integration and Multi-Task Prediction

Attended representations are integrated to form a unified feature vector for prediction. First, mean pooling is applied across the modality axis of the attention outputs, yielding cell-attended drug features 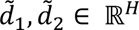 and drug-drug interaction features 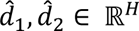. The drug-attended cell representation is computed as the projected mean of the two cell→drug outputs, 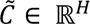.

These are concatenated into a composite vector:

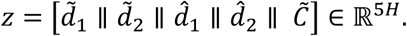

This vector 𝑧 is passed through a feature-integration MLP, which acts as a regularising bottleneck, producing a shared trunk representation ℎ ∈ 𝑅^256^. From this trunk, four independent two-layer MLP heads generate the final synergy predictions:

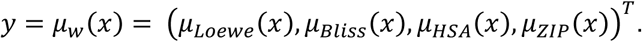

The main predictive model is trained by minimising a weighted Mean Squared Error (MSE) loss on these outputs.

#### 5. Uncertainty Quantification

AlgoraeOS decomposes predictive uncertainty into its aleatoric and epistemic components. Data-dependent (aleatoric) noise is modelled by a separate head that predicts a structured covariance matrix for the four synergy tasks from the detached feature vector 𝑧 via the mentioned equations. This head is trained independently by minimising the regularised multivariate Gaussian negative log-likelihood (NLL). This decoupled optimisation ensures the uncertainty head learns the empirical residual structure without affecting the mean predictor. Model uncertainty (Epistemic) is estimated at inference using Monte-Carlo Dropout (MC-Dropout) over *T* stochastic forward passes through the predictive trunk and heads. The epistemic covariance is the empirical covariance of these predictions. The total predictive uncertainty is the sum of these components, yielding a full predictive distribution. Normal-approximate 95% confidence intervals (±1.96 𝜎) are derived from the marginal variances. This decomposition improves predictive calibration by attributing heteroscedastic measurement noise to the aleatoric term and distributional or OOD uncertainty (e.g., unseen drugs, rare dose regimes, or tissue-specific shifts) to the epistemic component, enabling uncertainty-aware and risk-sensitive decision-making in downstream analyses.

### Model Rationale and Design Principles

The design of AlgoraeOS integrates strong domain priors to learn biologically plausible interactions. By employing pathway-aware cell embeddings and modality-specific drug encoders, the model grounds its representations in established biology and chemistry. This structured approach is balanced with a flexible cross-modal attention hub that captures context-dependent, non-additive relationships.

This hybrid design confers significant advantages in sample efficiency and predictive calibration over standard architectures, such as simple early- or late-fusion MLPs and vanilla Transformers. These benefits stem directly from two key architectural choices: 1) the explicit conditioning of each drug’s representation on both the cellular context and the partner drug, and 2) the structured, low-rank modelling of cross-task residual covariance, which captures dependencies between synergy metrics. The result is a computationally tractable model that learns robust, generalizable representations from limited data.

We acknowledge several limitations. The model’s performance is sensitive to the quality and completeness of the underlying pathway annotations. Furthermore, the use of Monte-Carlo Dropout provides only an approximation of epistemic uncertainty, which may be underestimated in highly over-parameterised regimes. Finally, significant imbalances across synergy label distributions may require careful loss scaling to ensure stable multi-task learning.

### Ablation Studies and Design Justification

Ablation studies were conducted to validate the contributions of key architectural and training choices in AlgoraeOS.

#### Cross-Modal Attention

Removing the attention hub reverts the model to a concatenation-based fusion baseline. This significantly degrades performance on context-dependent interactions and increases miscalibration under cell-line shifts, confirming the importance of dynamic feature conditioning.

#### Pathway Normalisation

Disabling length normalisation in the pathway encoder introduces a strong bias toward large gene sets, diminishing the biological specificity and interpretability of the learned cell-line representations.

#### Loss Weighting

Omitting the inverse-Gaussian loss weighting improves the aggregate Root Mean Squared Error (RMSE) but at the cost of higher prediction errors for strong synergistic combinations. This trade-off is unacceptable for clinical applications, where accurately identifying high-synergy pairs is paramount.

#### Uncertainty Quantification

The dual uncertainty components are critical. Excluding the aleatoric head results in overconfident and mis-calibrated predictions in heteroscedastic data regimes. Similarly, removing Monte-Carlo Dropout eliminates epistemic uncertainty estimates, impairing the model’s ability to detect out-of-distribution samples. The aleatoric head’s design, which uses detached features, intentionally decouples the optimisation of the mean and covariance. This stabilises training by preventing the uncertainty objective from interfering with the primary predictive task.

### Training Protocol

The model was trained using a three-phase schedule designed to promote stable convergence and accurate uncertainty estimation.

- **Phase A: Initial Stabilisation.** The model was first trained by minimising a standard Mean Squared Error (MSE) loss, with uniform weighting across all four synergy tasks. This phase established a stable baseline for the predictive representations.
- **Phase B: Tail-Aware Fine-Tuning.** To counteract regression-to-the-mean and improve performance on clinically relevant high-synergy examples, the MSE loss was augmented with an inverse-Gaussian weighting scheme. This approach selectively up-weights samples in the tails of the synergy distributions, using task-specific scales of for Loewe, Bliss, HSA, and ZIP, respectively.
- **Phase C: Aleatoric Uncertainty Learning** Finally, while retaining the weighted MSE loss, the aleatoric uncertainty head was activated. Its contribution to the total loss was governed by a coefficient, 𝜆_ale_, which was linearly ramped from 0 to its final value of 0.05 over 50 epochs to ensure smooth convergence.

### Concentration encoding

To model dose-dependent effects, we developed a concentration encoder that transforms a given concentration, *c* ≥ 0, into a fixed-length embedding, *e*(*c*) ∈ 𝑅^64^. This encoder operates in log-concentration space, 𝑙 = 𝑙𝑜𝑔(*m*𝑎*x*(*c*, 10^−12^), aligning with pharmacological conventions where dose-response curves are typically sigmoidal.

The encoding process begins by constructing a feature vector, 𝛷(𝓁), from a concatenated basis of functions:

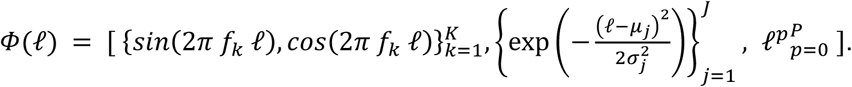

This basis includes Fourier features for k=1 to 8 to capture smooth, multi-scale variations, a bank of Radial Basis Functions (RBFs) with learnable centres and widths for j=1 to 16, to provide localised sensitivity around key concentration ranges (e.g., EC50-like regions), and low-order polynomials for p=0 to 4 to model global trends. This composite basis offers a universal and data-adaptive function family for dose effects, avoiding commitment to a single parametric form while retaining a strong inductive bias for pharmacologically plausible curves. For instance, with zero or unknown concentrations (*c* ≤ 10^−12^), a multiplicative mask ensures 𝛷(𝓁) = 0, preventing spurious signals. The final embedding, *e*(*c*), is obtained by projecting 𝛷(𝓁) through a small multi-layer perceptron (MLP).

Dosage information is integrated into the model through two complementary mechanisms to ensure its influence on predictions:

#### Feature-wise Linear Modulation (FiLM)

The contextualised drug representation, derived from cross-modal attention, is conditioned on its corresponding concentration. The concentration embedding, *e*(*c*), is passed through an MLP to produce modulation parameters (𝛾, 𝛽). These parameters then scale and shift the drug’s feature vector 𝑧′ = 𝑧 + (𝛾 - 1) ⊙ 𝑧 + 𝛽. The FiLM layer is initialised as an identity function to ensure stable training. This mechanism allows the model to modulate drug features in a context-aware manner, considering both the cell line and any interacting drugs.

#### Direct Embedding Concatenation

The concentration embeddings for each drug, *e*(*c*₁) and *e*(*c*₂), are explicitly concatenated with the pooled drug and cell features before being fed into the final feature-integration MLP. This provides downstream predictors, including the multivariate aleatoric uncertainty head, with direct access to the raw dose information, supporting additive effects and cross-task learning.

This dual-pathway architecture—combining multiplicative FiLM modulation with additive concatenation—provides a robust and principled method for integrating concentration data. It enables the model to capture context-sensitive and asymmetric drug responses, crucial for synergy prediction. Furthermore, by allowing the dose embeddings to inform the uncertainty prediction head, the model can learn to express higher uncertainty in dose regimes with sparse data, thereby improving calibration.

### Multi-Task Synergy Prediction Model

We quantify drug combination activity relative to a null reference model at each dose pair (*d*_𝐴_, *d*_*B*_). The synergy score, 𝑆 is defined in the inhibition space 𝐸 where 𝐸 = 1 - 𝑣*i*𝑎𝑏*i*𝑙*i*𝑡*y* as:

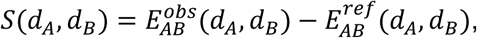

here 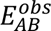 is the experimentally observed inhibition, while 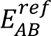 is the expected inhibition under a specific null hypothesis of non-interaction. We consider four standard reference models:

1. **Bliss Independence**: Assumes probabilistic independence between the effects of two agents. The reference surface is defined as 𝐸_*B*𝑙*i*𝑠𝑠_ = 𝐸_𝐴_ + 𝐸_*B*_ - 𝐸_𝐴_𝐸_*B*_. This local model is suitable for drugs with distinct mechanisms of action.
2. **Highest Single Agent (HSA)**: A conservative model positing that a non-interacting combination cannot exceed the effect of its most active component. The reference is 𝐸_𝐻𝑆𝐴_ = max (𝐸_𝐴_, 𝐸_*B*_). HSA provides a high bar for efficacy, highlighting combinations with clear performance benefits.
3. **Loewe Additivity**: Formalises dose-additivity for mechanistically similar agents. A combination is deemed additive if its Combination Index (CI) is one, where

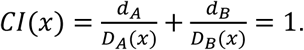

Here 𝐷_𝐴_(*x*) and 𝐷_*B*_(*x*) are the doses of drugs A and B required to achieve the effect level *x* alone. The reference effect, 𝐸_𝐿𝑜*ewe*_(*d*_𝐴_, *d*_*B*_), is the effect *x* for which 𝐶𝐼(*x*) = 1, found by inverting the single-agent dose-response curves.
4. **Zero Interaction Potency (ZIP)**: Assumes that in a non-interacting pair, the dose-response curve of one drug is not affected by the presence of the other. The reference surface, 𝐸_𝑍𝐼𝑃_, is synthesised from the monotherapy Hill curves such that the potency and slope parameters remain unchanged in the combination.

These four models probe different aspects of non-interaction—probabilistic independence (Bliss), pragmatic dominance (HSA), dose equivalence (Loewe), and potency preservation (ZIP), providing a multi-faceted view of drug interaction. We employ a multi-task learning (MTL) framework to jointly predict the vector of four synergy scores, *y* = (𝑆_𝐿𝑜*ewe*_, 𝑆_*B*𝑙*i*𝑠𝑠_, 𝑆_𝐻𝑆𝐴_, 𝑆_𝑍𝐼𝑃_)^*T*^∈ 𝑅^4^. Given input features x representing drug and cell-line properties, our model predicts the conditional probability distribution of the synergy scores as a multivariate Gaussian, 𝒩(𝛍(𝐱), 𝚺(𝐱)). The mean vector 𝛍(𝐱) provides the point estimates for each synergy score, while the covariance matrix 𝚺(𝐱) captures both the per-score uncertainty and the correlations between them.

This probabilistic MTL approach offers significant advantages over training independent models. A shared feature encoder learns common representations of underlying biological and chemical mechanisms, enhancing data efficiency and reducing model variance. By explicitly modelling the covariance 𝚺(𝐱), the framework learns the statistical dependencies among the synergy metrics. This learned structure acts as a powerful inductive bias, enforcing coherence across predictions (e.g., avoiding strong HSA synergy predictions when monotherapy effects are weak) and enabling the model to denoise less reliable scores (e.g., unstable Loewe scores) by borrowing information from more robust metrics. In contrast, separate models discard this crucial cross-metric signal, leading to inconsistent predictions, underestimated uncertainty, and no principled way to aggregate evidence for decision-making.

For practical implementation, monotherapy dose-response curves, which provide the ground truth for Loewe and ZIP scores, must be fitted with care. The covariance matrix 𝚺(𝐱) is parameterised using a stable transformation, such as a Cholesky decomposition, to ensure it remains positive definite. Model performance is assessed not only by marginal metrics like RMSE per score but also by proper scoring rules such as the mean negative log-likelihood (NLL) and calibration analysis. For hit prioritisation, we advocate for uncertainty-aware criteria that leverage the full predictive distribution, rather than relying on simple thresholds on point predictions. This MTL framework, by unifying the prediction of complementary synergy metrics within a single probabilistic model, yields more accurate, coherent, and calibrated results, providing a principled foundation for risk-sensitive decision-making in drug combination discovery.

### Model Evaluation Metrics

We evaluated the performance of our predictive models using a standard suite of metrics to provide a comprehensive assessment of prediction quality. Given a set of 𝑛 ground truth values *y*_*i*_ and corresponding predictions 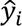, we define the following:

#### 1. Mean Absolute Error (MAE)

MAE quantifies the typical absolute deviation between predicted and actual values. Due to its linear penalty on the error, it is less sensitive to large outliers compared to the RMSE.

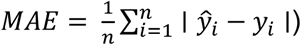

#### 2. Root Mean Squared Error (RMSE)

RMSE measures the magnitude of the error, with its quadratic term placing greater weight on larger deviations. It is thus more sensitive to outliers and provides insight into tail risk.

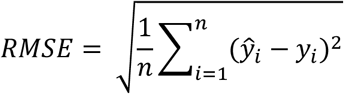

#### 3. Coefficient of Determination (𝑅^2^)

𝑅^2^ metric or “variance explained,” indicates the proportion of the variance in the dependent variable that is predictable from the independent variables. It provides a scale-invariant measure of how well the model’s predictions approximate the true values compared to a simple mean-based model 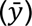.

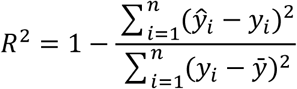

#### 4. Pearson Correlation Coefficient (PCC)

PCC measures the linear relationship between predicted and actual values. It is invariant to scaling and shifting, making it an effective measure of ranking fidelity. A high PCC indicates that the model correctly captures the direction and relative magnitude of changes, though not necessarily the absolute values.

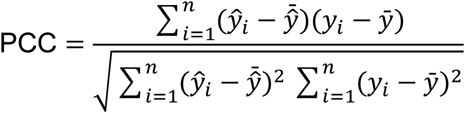

Together, this suite of metrics provides a holistic view of model performance. MAE and RMSE assess absolute prediction accuracy under different loss assumptions,𝑅^2^ contextualises the error relative to the inherent variability of the data, and PCC evaluates the model’s ability to preserve the rank-ordering of the outputs.

### In Vitro Validation Assays

In vitro validation was undertaken at the Victorian Centre for Functional Genomics (VCFG) with the cell lines (T98G, BT-20, 22Rv1) sourced from Cell Bank Australia and STR profiled by the Peter Mac genotyping core to confirm authenticity. Briefly, cells were thawed one week prior to the day of seeding and grown in T75 flasks to approximately 90% confluency. On the day of seeding, all the cell lines were washed once with PBS^−/−^ (without MgCl_2_ and CaCl_2_) then incubated with trypsin-EDTA for 6-10mins in a 37°C, 5% CO_2_ incubator for detachment. The trypsin was neutralised and cells were pelleted for 5mins at 1200rpm using an Eppendorf 5810R centrifuge. The media was aspirated and cell were resuspended in 3-5ml of culture media. Each cell line was counted twice using a Countess 3 FL automated cell counter (Thermo Fisher Scientific). The cell suspensions were diluted to a density of 1.5×10^4^/mL for T98G, 3×10^4^/mL for BT-20, and 7×10^4^/ml for 22Rv1. Cells were seeded using an automated dispenser (EL406, BioTek Agilent) into 384 well black walled plates (Corning cat #3764) at 50uL/well at high speed with a dispense height of 336. Plates were pulse spun at 500rpm then allowed to rest on a Sciclone G3 robot deck (specifically flat) for 10mins at room temperature before being placed into the LiCONiC incubator (37°C, 5% CO_2_) until drug treatment.

All drugs were sourced from MedChem Express and solubilized in dimethyl sulfoxide (DMSO), with the exception of one drug which was solubilized in water. Cannabidiol (CBD) was sourced from Sapphire Biosciences (Cat# 06-1953). Staurosporine and Mitomycin (in house VCFG at 50mM stock concentration) were used as positive controls for cell death and growth inhibition (final concentration 10, 1, 0.1uM). Addition of DMSO solubilized drugs was performed using the D300e automated dispenser (Tecan). Prior to combination screening, the drug concentration range for each drug was determined for cell line to produce an approximate inhibition range of 0-50% where possible. Combination screening using a 6×6 dose range arrays was performed with randomized layout. The water solubilized drug was added manually. Cells were sedimented by pulse spin at 500rpm (Eppendorf 5810R) and incubated for 48 h in a LiCONiC incubator (37°C, 5% CO_2_). Two technical replicates were performed for each treatment, with requisite DMSO controls.

After 48h growth the cells were fixed and stained using the EL406 automated dispenser/washer with a Z-height of 36 for aspiration and 336 for dispensing, at high speed. The culture media was removed and 25ul of fixative (4% PFA in PBS−/−) was dispensed and incubated for 10mins at room temperature. The fixative was removed and cells washed with 50ul of PBS−/−, then 25ul of the staining solution (0.2% Triton X-100, 1mg/ml DAPI, 50mM Tris-HCl pH 7.6 in water) was dispensed into the wells. The cells were allowed to stain for 20mins at room temperature. The stain was removed, and the cells were washed 2 times with PBS−/−. The wells were dispensed with a final 50ul of PBS−/− then sealed (PlateLoc; automated aluminum heat seal) to prevent evaporation.

Cells were imaged on the CellInsight CX7 Pro (Thermo Fisher Scientific) at 10X objective with 9 fields per well, auto focusing every field. The cells were imaged using the brightfield and DAPI channels. Pipelines for nuclei segmentation using the DAPI channel were created using customized pipelines established via the open-source image analysis software CellProfiler (version 4.2.8). Batch scripts exported from CellProfiler were submitted to the Peter Mac High Performance Cluster (HPC) which runs on Red Hat 9 Linux nodes. Cell counts were exported and analysed using custom pipelines R (version 4.4.2).

## Supporting information

Supplementary File 1:comparison of machine learning models, Supplementary File 2: MoA classes, Supplementary File 3: Prospective in vitro validation

## Data and Code Accessibility

The raw training datasets are available from the DrugComb data portal: https://drugcomb.fimm.fi. All curated and pre-processed datasets used in this study have been deposited in the AlgoraeOS GitHub repository https://github.com/VafaeeLab/AlgoraeOS to support reproducibility and future benchmarking against our method. The full model implementation is proprietary. To enable reproducibility and broader use, we provide access to model predictions through a public web portal: https://vafaeelab.github.io/AlgoraeOS_UI/.

## Acknowledgement

The Victorian Centre for Functional Genomics (K.J.S: RRID:SCR_0255820) is funded by the Australian Cancer Research Foundation (ACRF), Phenomics Australia (https://ror.org/0201hm243), through funding from the Australian Government’s National Collaborative Research Infrastructure Strategy (NCRIS) program, the Peter MacCallum Cancer Centre Foundation and the University of Melbourne Collaborative Research Infrastructure Program. This research was supported by computational resources provided by the Australian National Computational Infrastructure (NCI), enabled by the Australian Government through the National Collaborative Research Infrastructure Strategy (NCRIS). This project was made possible by CSIRO’s Next Generation Artificial Intelligence Graduates Program (GA221787) funded by the Australian Government. The authors wish to thank J. Luu for support in program administrative logistics.

## Supplementary Files

**Supplementary File 1**: A comprehensive comparison of machine learning models for drug synergy prediction. An interactive version of the table is also available online: https://vafaeelab.github.io/drug-synergy-models

**Supplementary File 2**: A Comprehensive catalogue of drug mechanism-of-action (MoA) classes, enabling mechanism-aware analysis and stratification of model performance.

**Supplementary File 3**: Prospective in vitro validation dataset for dose-specific inhibition measurements. Detailed results for all 2,592 dose–combination measurements generated using Cannabidiol (DB09061) as an anchor compound tested in combination with 24 partner drugs across three cancer cell lines (T98G, 22Rv1, BT-20). The file includes compound identifiers (deidentified), cell line labels, concentration pairs, and corresponding observed inhibition values (%) for each compound–cell line–dose triplet.

