## Supplementary File 1:comparison of machine learning models, Supplementary File 2: MoA classes, Supplementary File 3: Prospective in vitro validation for "Uncertainty-Aware Deep Learning for Multi-Metric and Dose-Specific Prediction of Drug Synergy": Supplementary_Figures.docx

Professor Fatemeh Vafaee

School of Biotechnology and Biomolecular Sciences

UNSW SYDNEY NSW 2052 AUSTRALIA

T: +61 (2) 9065 2699

E:

**Supplementary Figure 1.** Experimental Data for HSA Synergy Measurement (Page 3)

**Supplementary Figure 2**. Experimental Data for ZIP Synergy Measurement (Page 5)

**Supplementary Figure 3.** Experimental Data for Loewe Synergy Measurement (Page 7)

**Supplementary Figure 4.** Experimental Results for Ridge Regression (Page 8)

**Supplementary Figure 5.** Experimental Results for Linear Regression (Page 9)

**Supplementary Figure 6.** Experimental Results for MLP (Page 10)

**Supplementary Figure 7.** Experimental Results for Random Forest (Page 11)

**Supplementary Figure 8.** PCC between predicted and observed inhibition stratified by partner drug concentration level (Page 12)

| **a**  **Observed HSA**  **Predicted HSA** 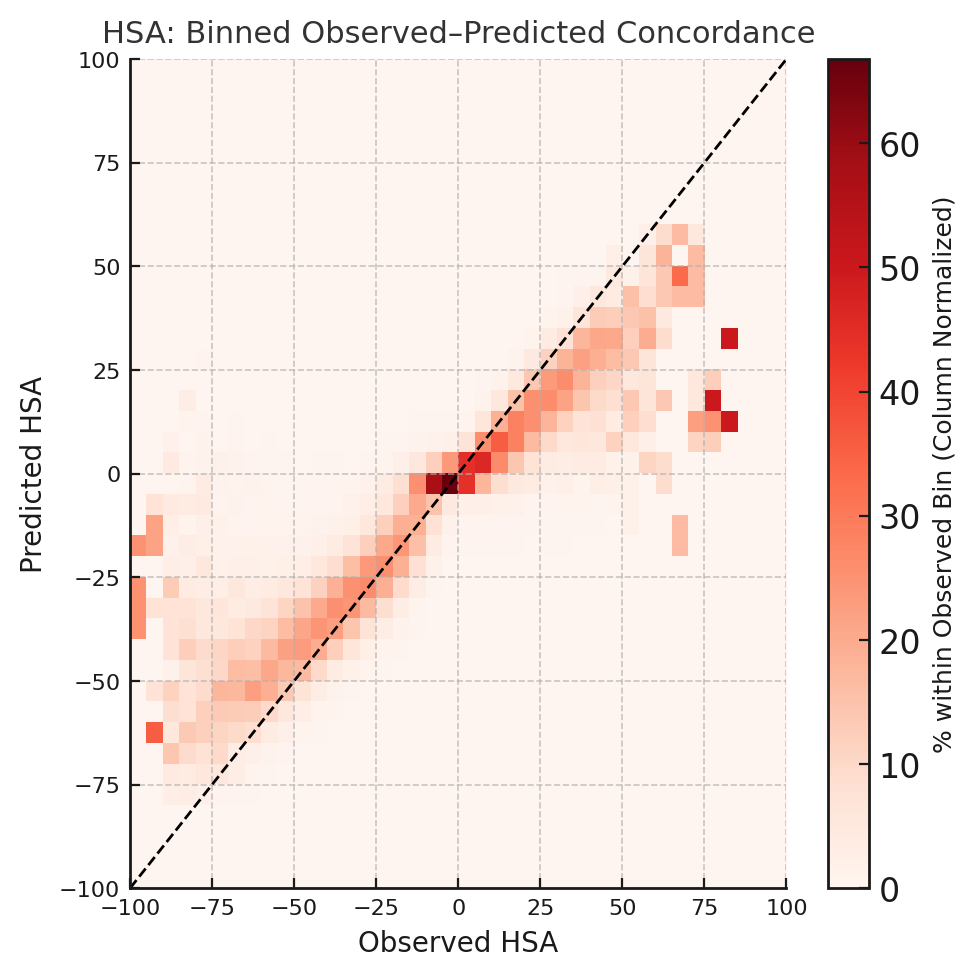 | 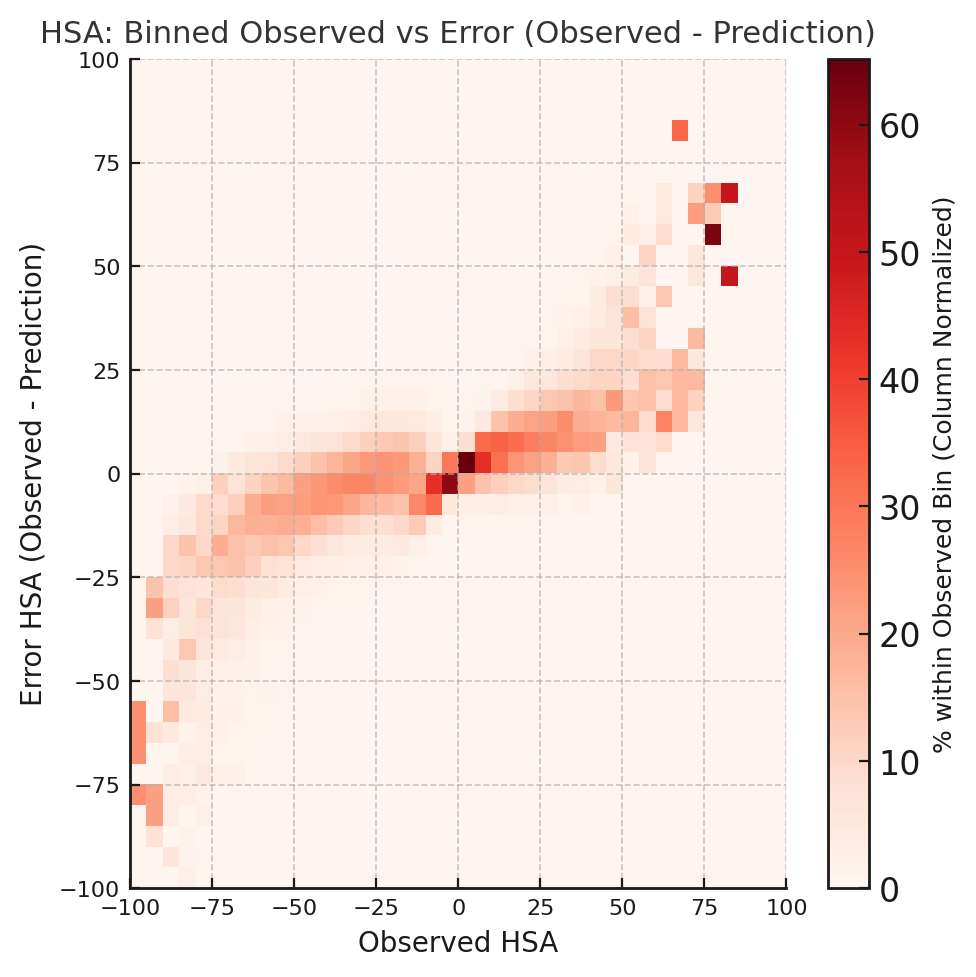 **b**  **Observed HSA**  **Error HSA (Observed -Prediction)** |
| --- | --- |
| **c** 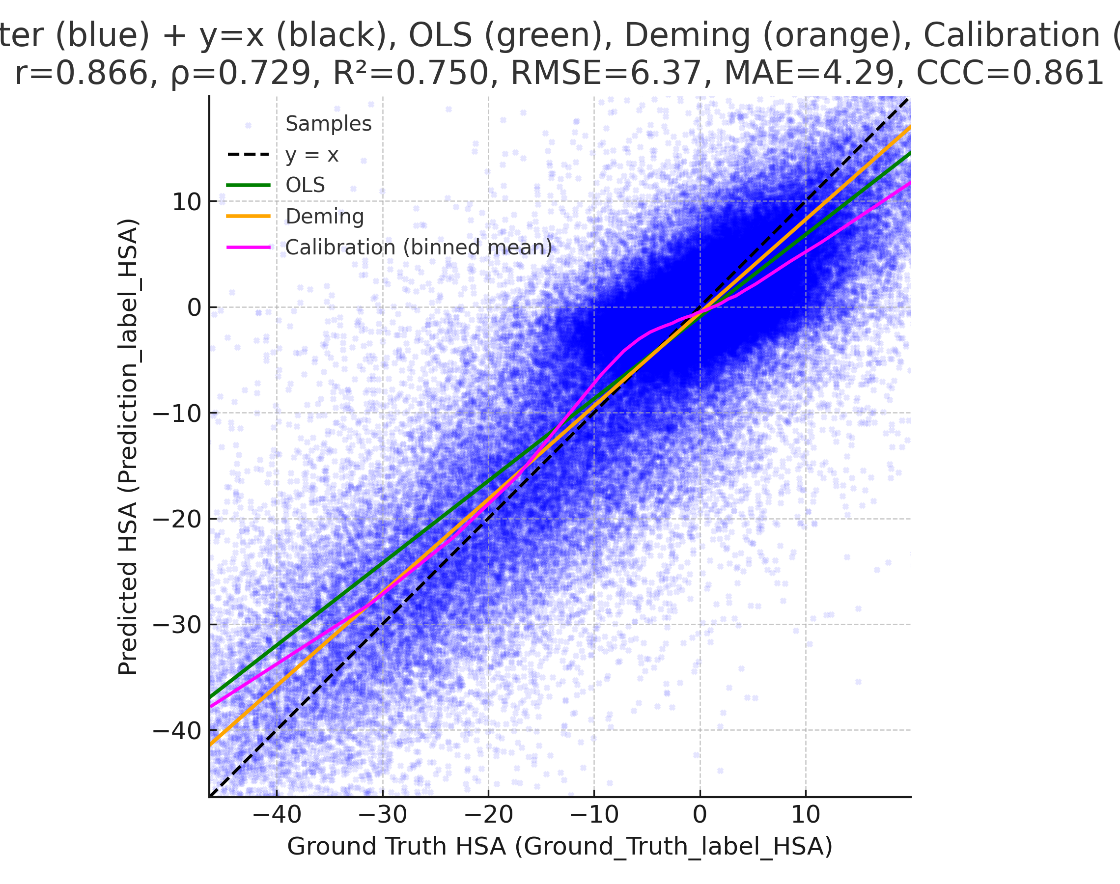 Scatter (blue), y=x (black), OLS (green), Deming (orange)  Calibration$: r=0.866, \rho= 0.729, R^{2}= 0.750, RMSE = 6.37, MAE = 4.29$  **Observed HSA (Ground Truth)**  **Predicted HSA** | |
| **d** 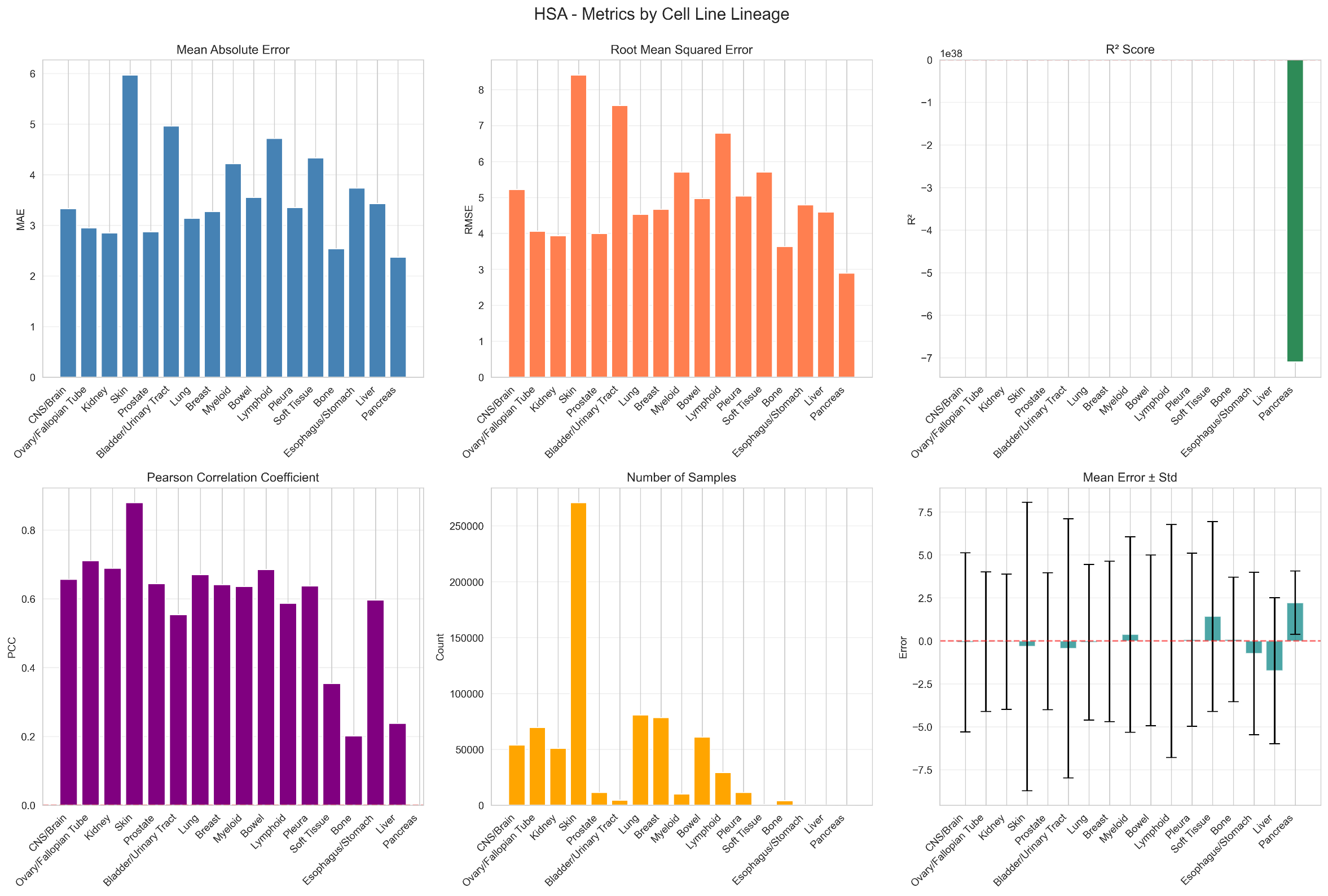 | 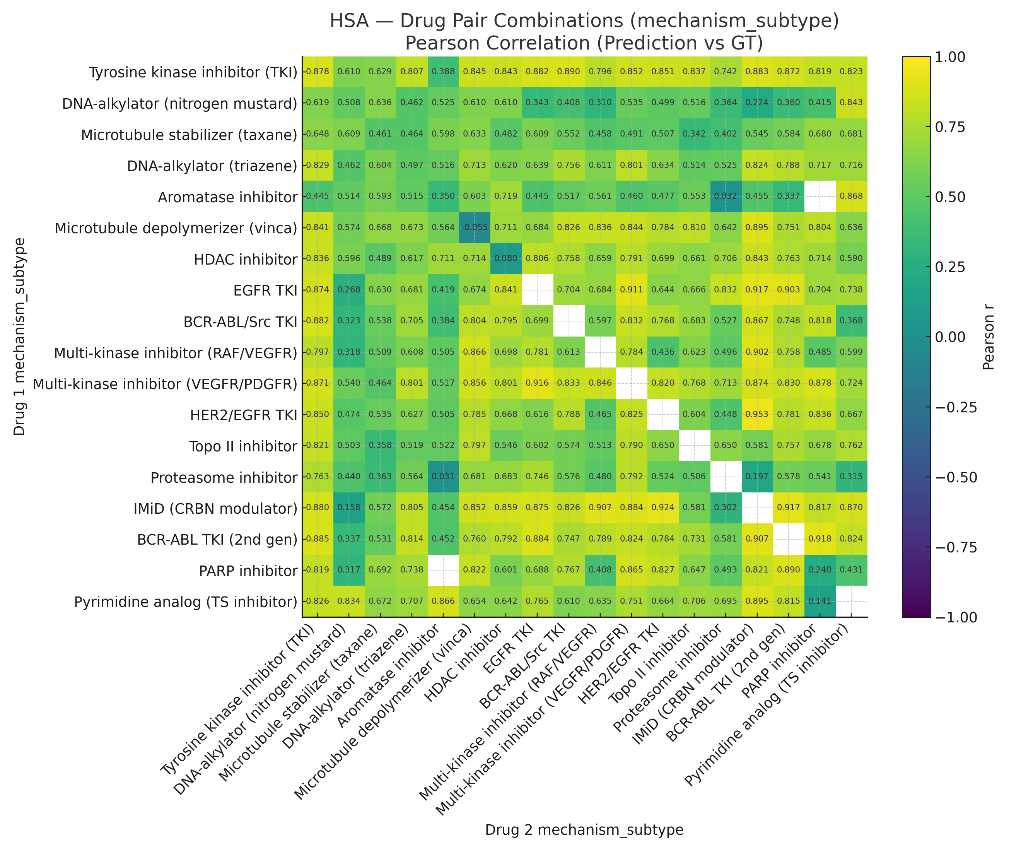 **e** |

**Supplementary Figure 1.**

(a)	Binned concordance between observed and predicted HSA values (bin size = 5), with color intensity indicating the percentage of samples within each bin.

(b)	Binned error distribution (observed – predicted HSA values) showing deviation across the observed range.

(c)	Scatter plot of predicted versus observed HSA synergy scores across all test folds; the yellow line: the identity line (y = x), and the grey line: the fitted regression.

(d)	Bar plot showing the number of observations per cancer tissue type (blue bars, log scale) and corresponding PCC between observed and predicted HSA.

(e)	Heatmap showing the Pearson correlation coefficient between observed and predicted HSA for drug pairs grouped by their mechanism of action (MoA) class.

| **a** 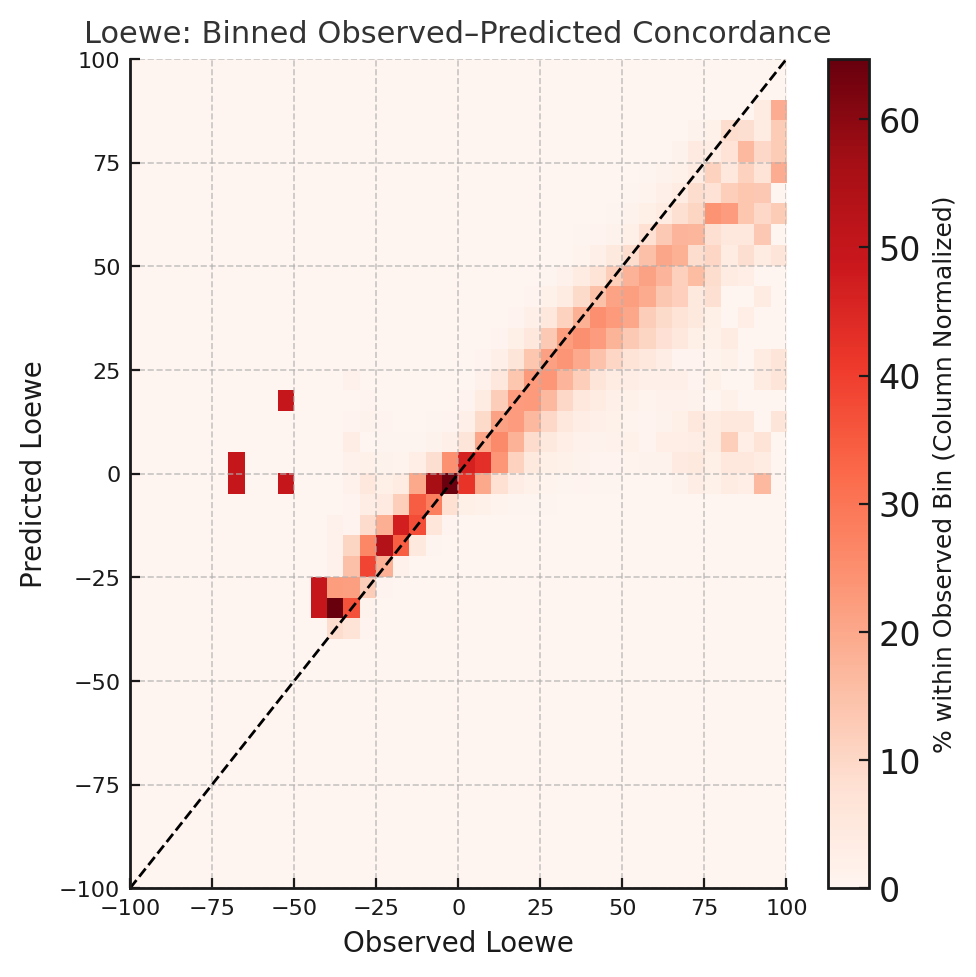 **Observed ZIP**  **Predicted ZIP** | **Observed ZIP**  **Error ZIP (Observed -Prediction)**  **b** 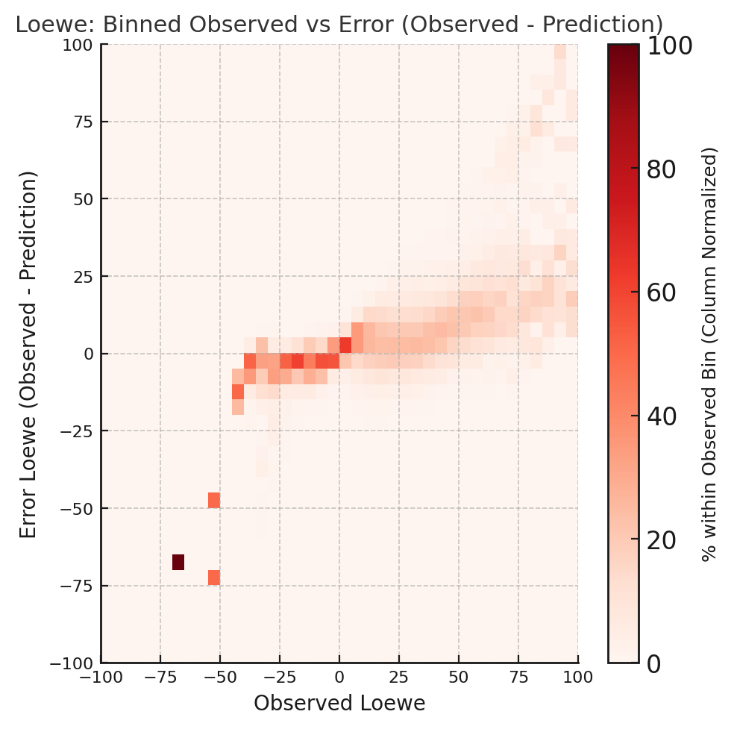 |
| --- | --- |
| **c**  **Observed ZIP**  **Predicted ZIP** 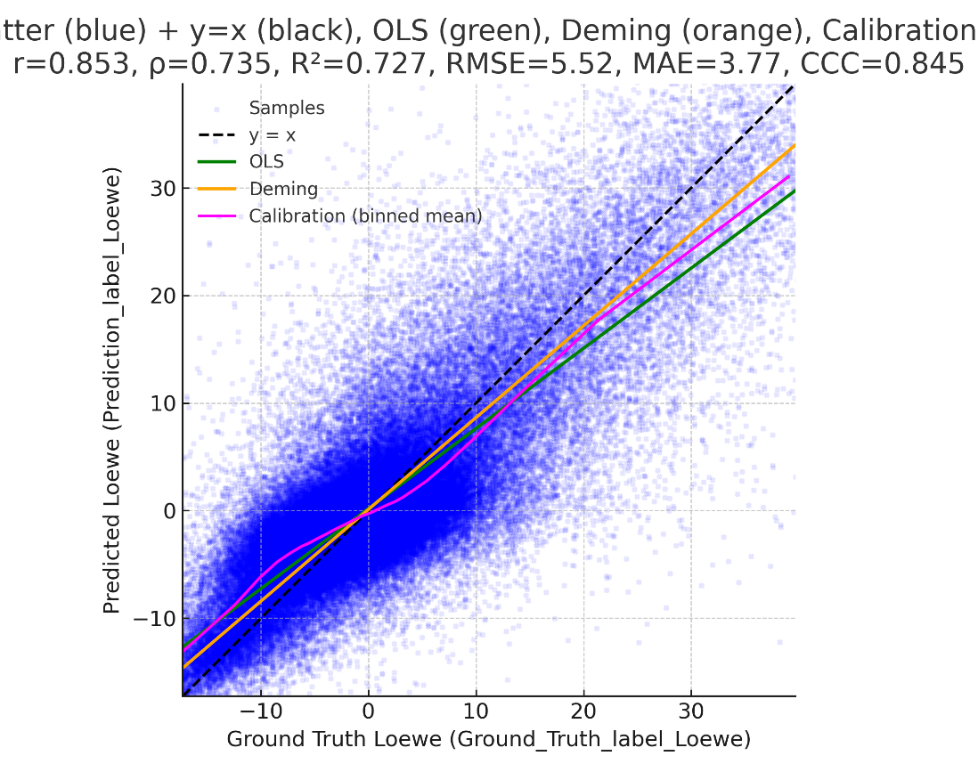 Scatter (blue), y=x (black), OLS (green), Deming (orange)  Calibration: $r=0.853, \rho= 0.735, R^{2} = 0.727, RMSE =5.52, MAE = 3.77$ | |
| **d** 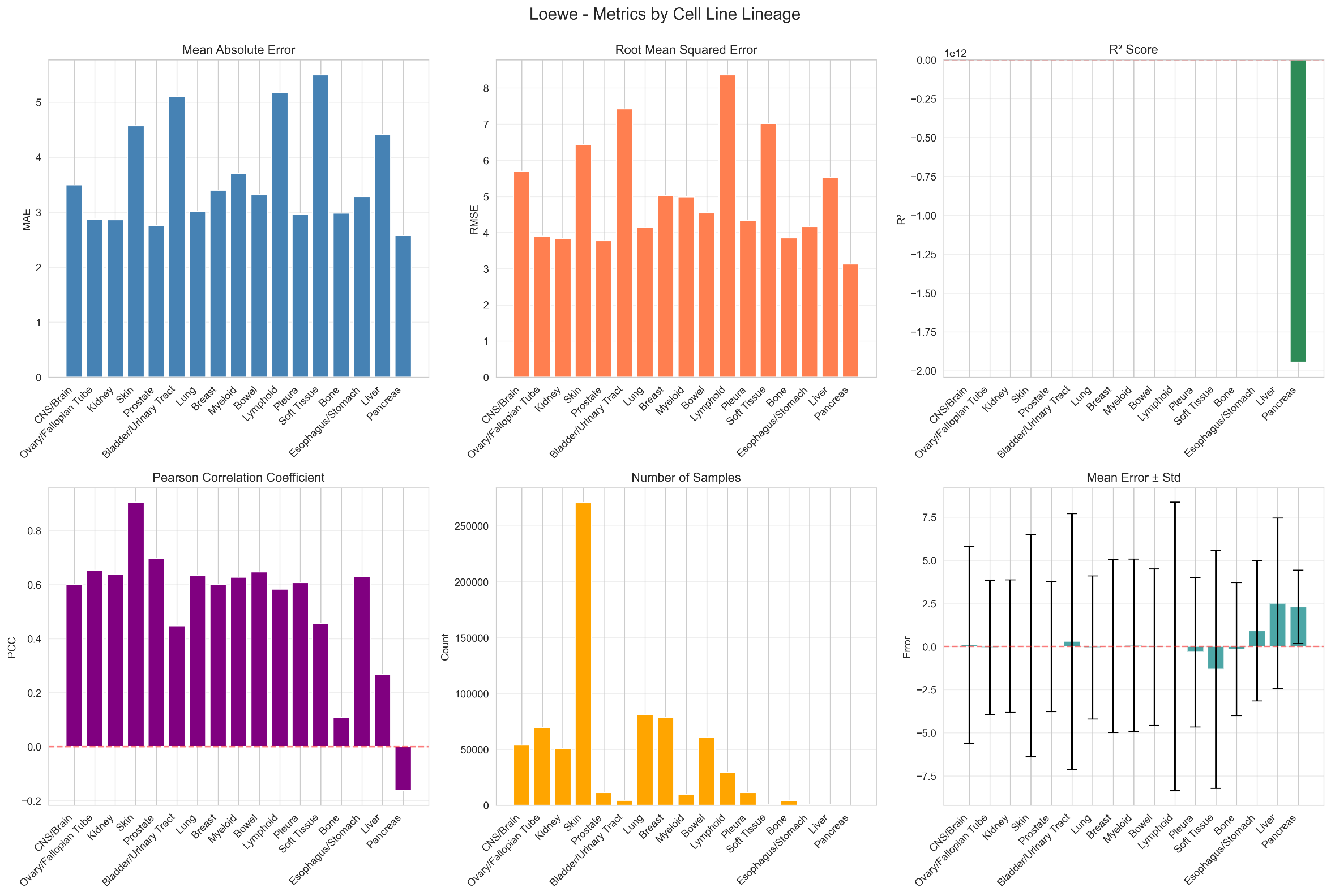 | **e** 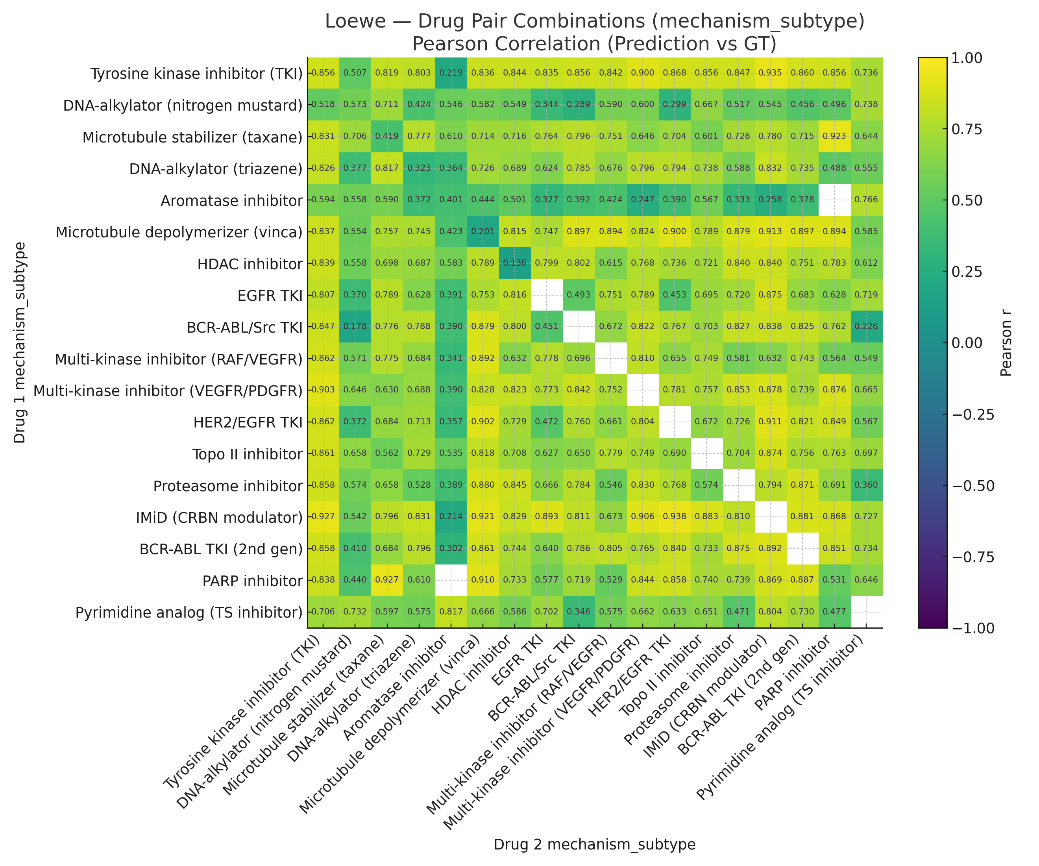 |

**Supplementary Figure 2.**

(a)	Binned concordance between observed and predicted ZIP values (bin size = 5), with color intensity indicating the percentage of samples within each bin.

(b)	Binned error distribution (observed – predicted ZIP values) showing deviation across the observed range.

(c)	Scatter plot of predicted versus observed ZIP synergy scores across all test folds; the yellow line: the identity line (y = x), and the grey line: the fitted regression.

(d)	Bar plot showing the number of observations per cancer tissue type (blue bars, log scale) and corresponding PCC between observed and predicted ZIP.

(e)	Heatmap showing the Pearson correlation coefficient between observed and predicted ZIP for drug pairs grouped by their mechanism of action (MoA) class.

| **a** 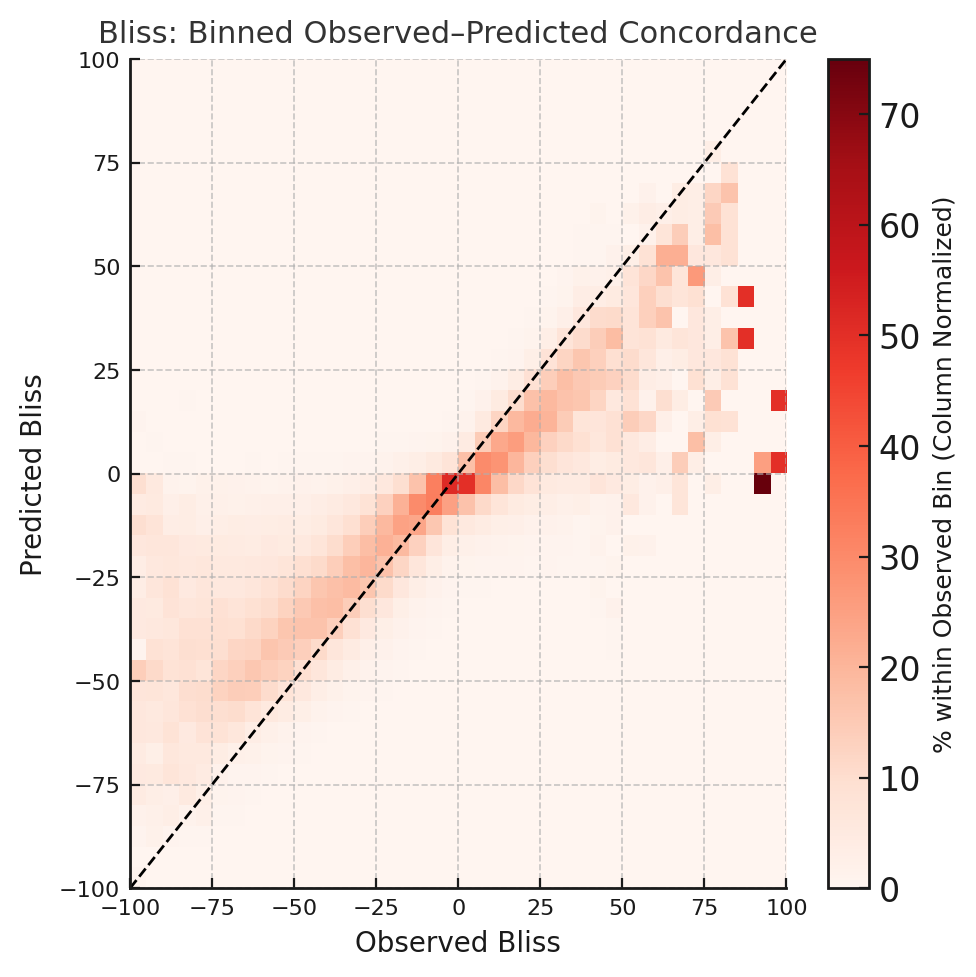 **Observed Loewe**  **Predicted Loewe** | **b** 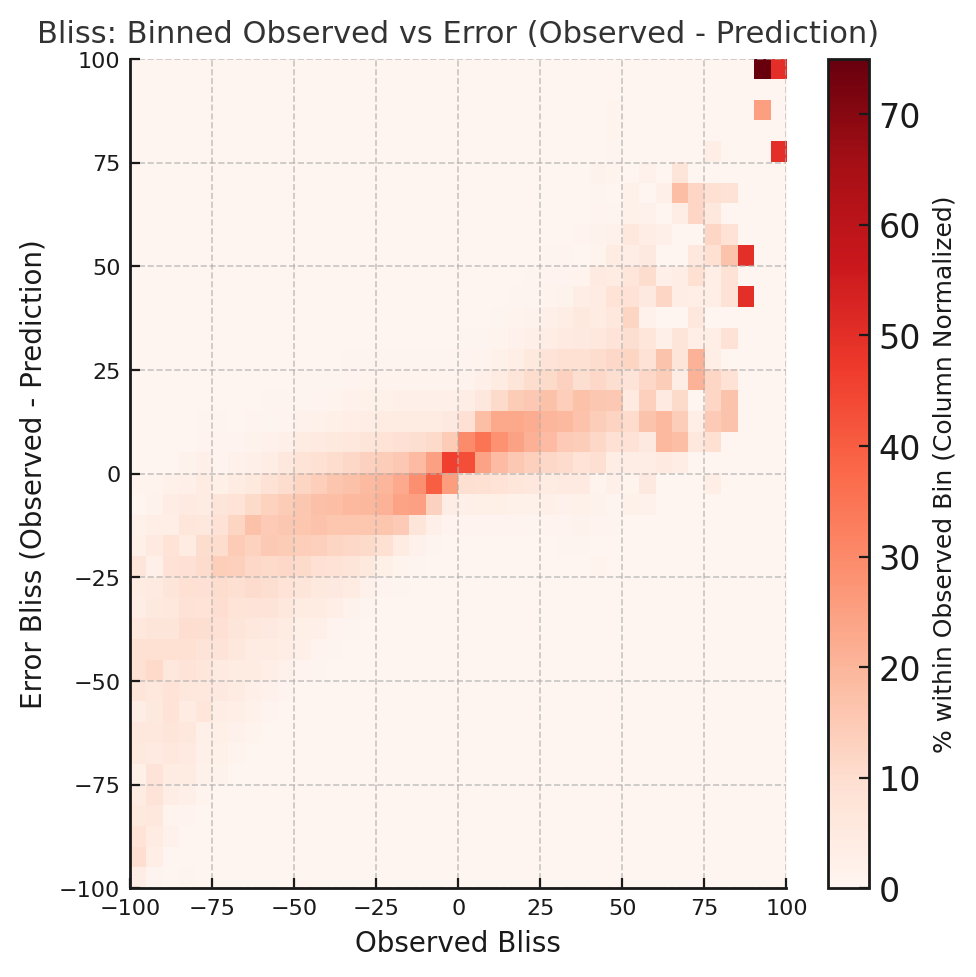 **Error Loewe (Observed -Prediction)**  **Observed Loewe** |
| --- | --- |
| 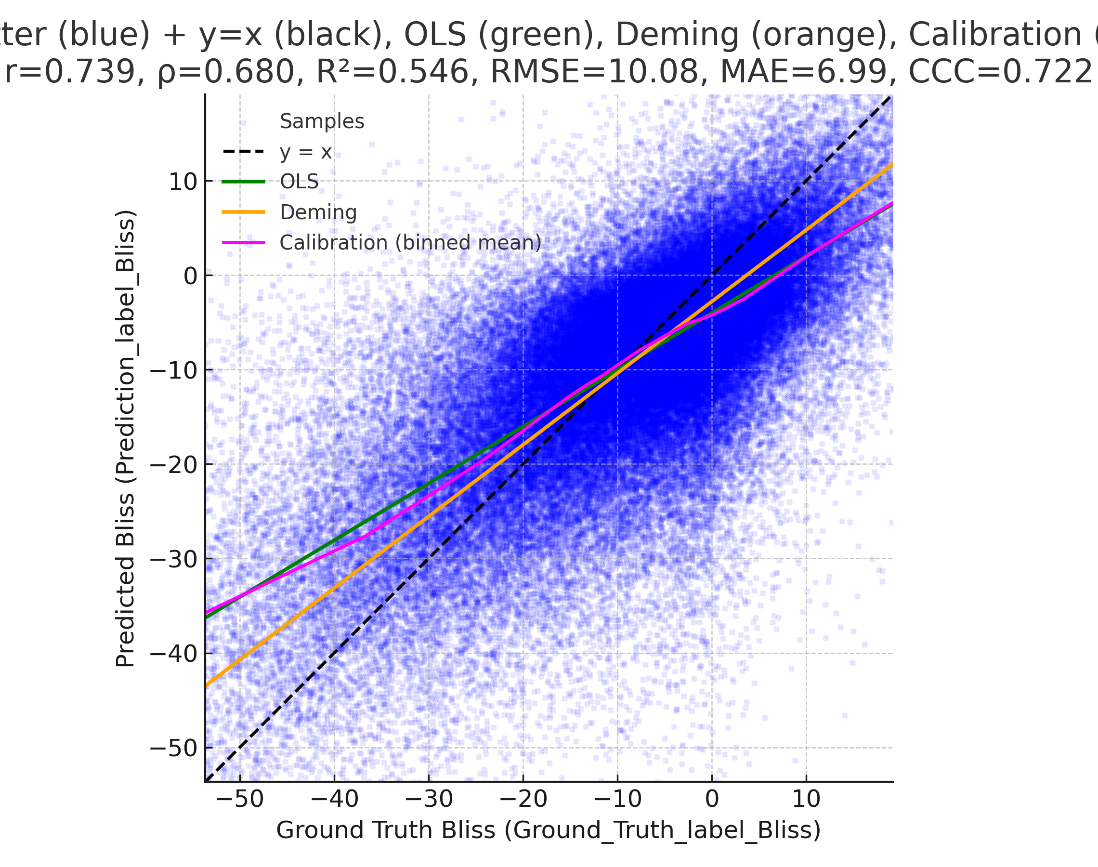 Scatter (blue), y=x (black), OLS (green), Deming (orange)  Calibration: $r=0.739, \rho= 0.680, R^{2} = 0.546, RMSE =6.99, MAE = 3.77$  **c**  **Predicted Loewe**  **Observed Loewe** | |
| **d** 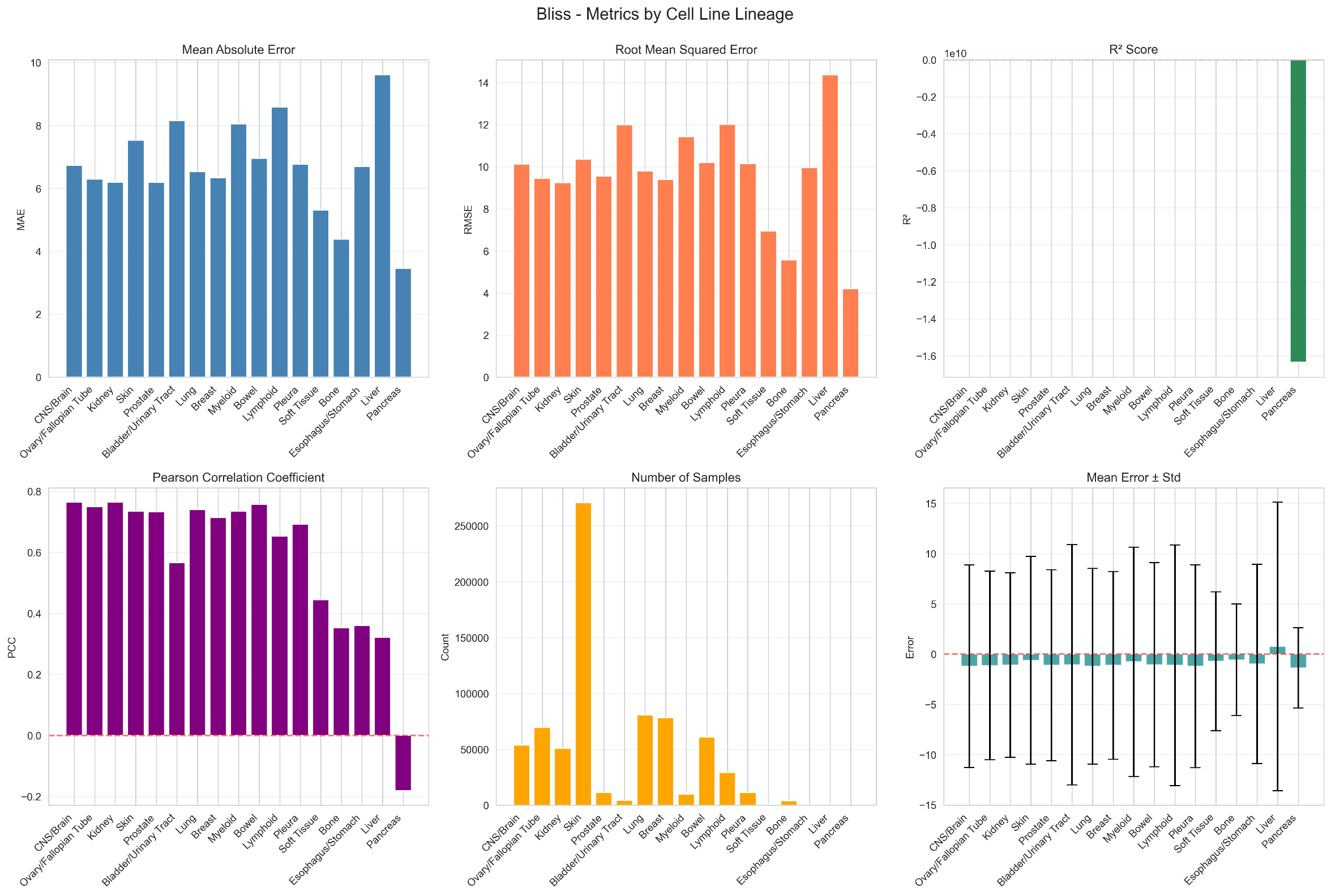 | **e** 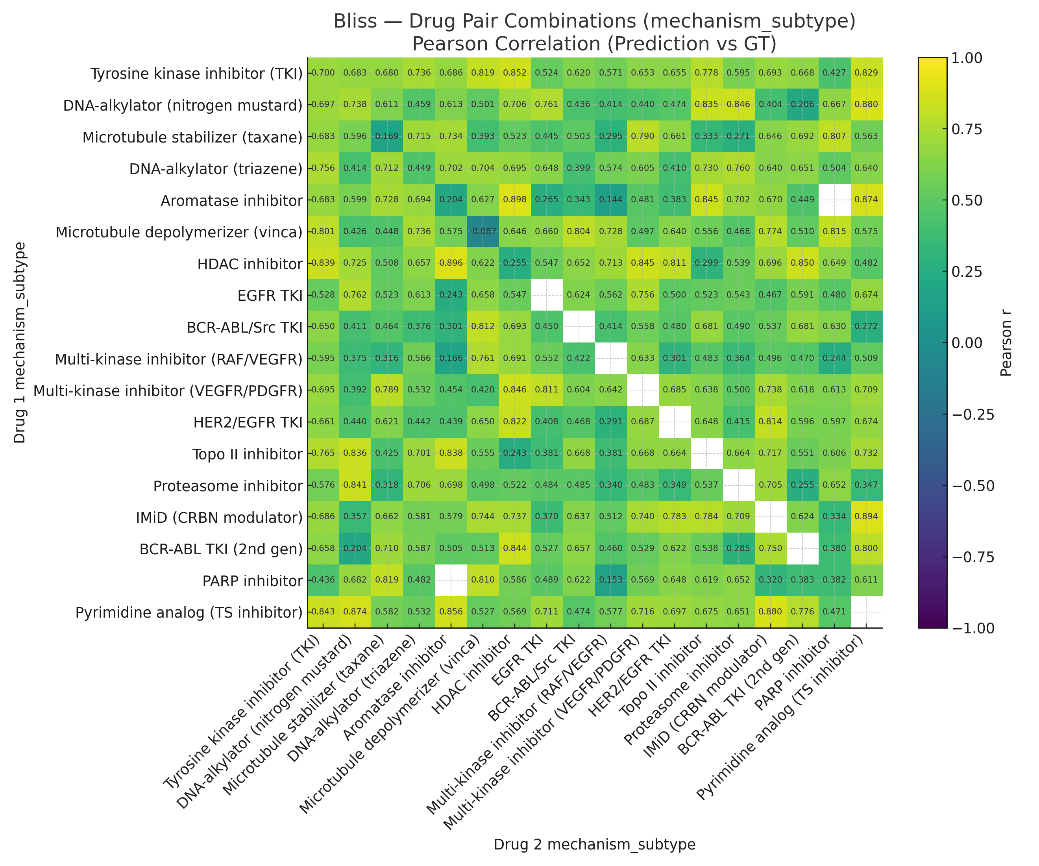 |

**Supplementary Figure 3.**

(a)	Binned concordance between observed and predicted Loewe values (bin size = 5), with color intensity indicating the percentage of samples within each bin.

(b)	Binned error distribution (observed – predicted Loewe values) showing deviation across the observed range.

(c)	Scatter plot of predicted versus observed Loewe synergy scores across all test folds; the yellow line: the identity line (y = x), and the grey line: the fitted regression.

(d)	Bar plot showing the number of observations per cancer tissue type (blue bars, log scale) and corresponding PCC between observed and predicted Loewe.

(e)	Heatmap showing the Pearson correlation coefficient between observed and predicted Loewe for drug pairs grouped by their mechanism of action (MoA) class.

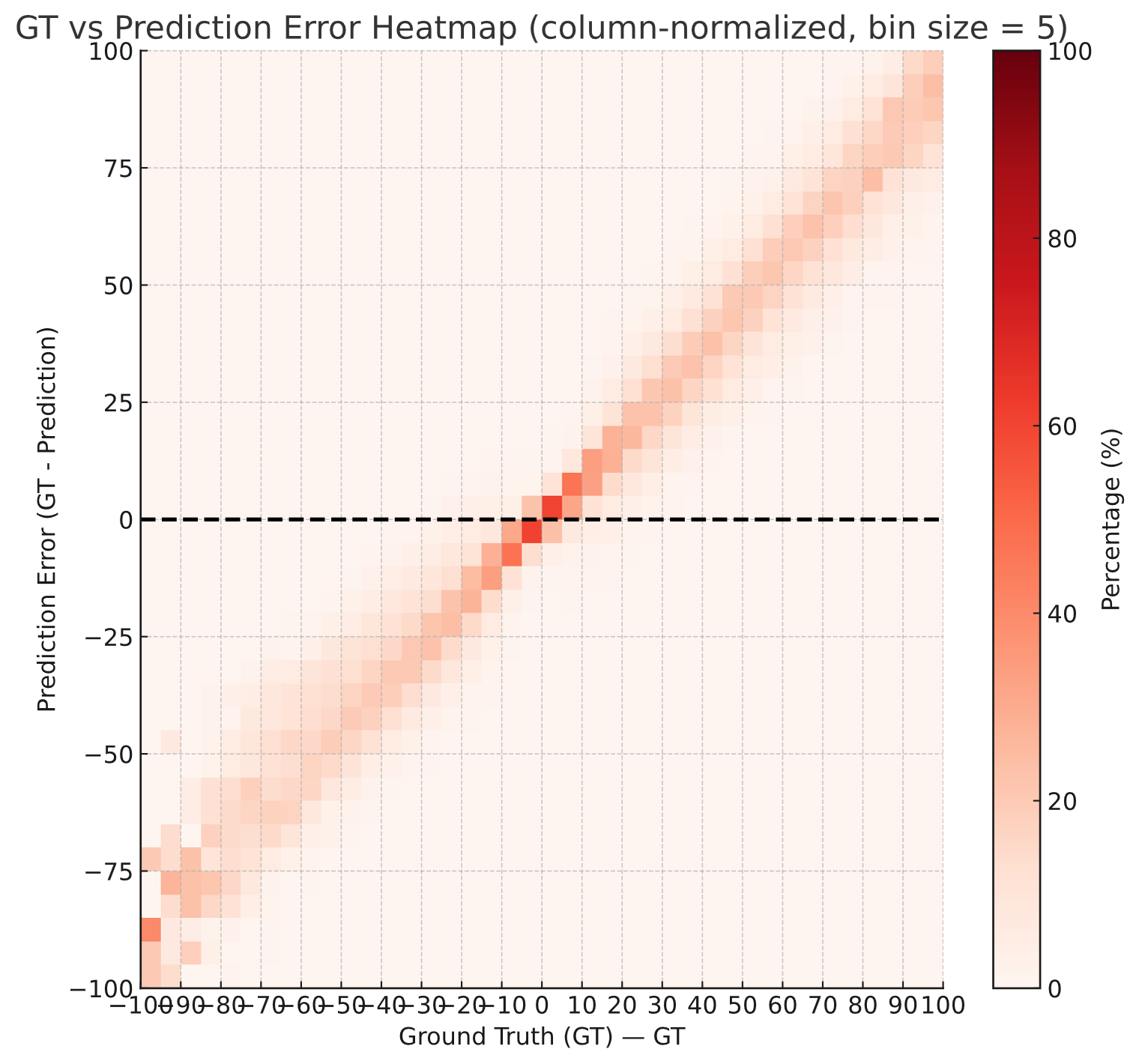

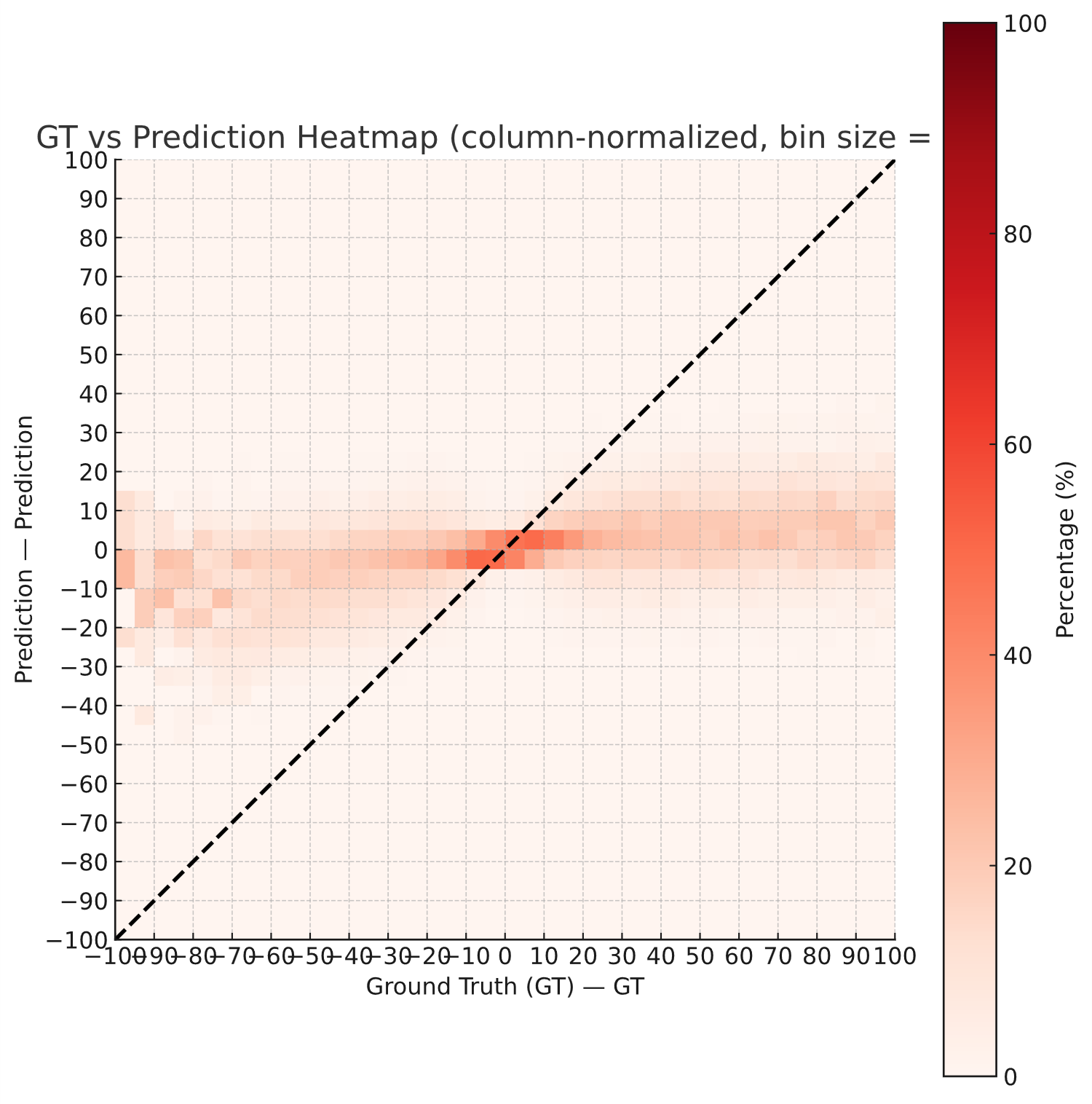

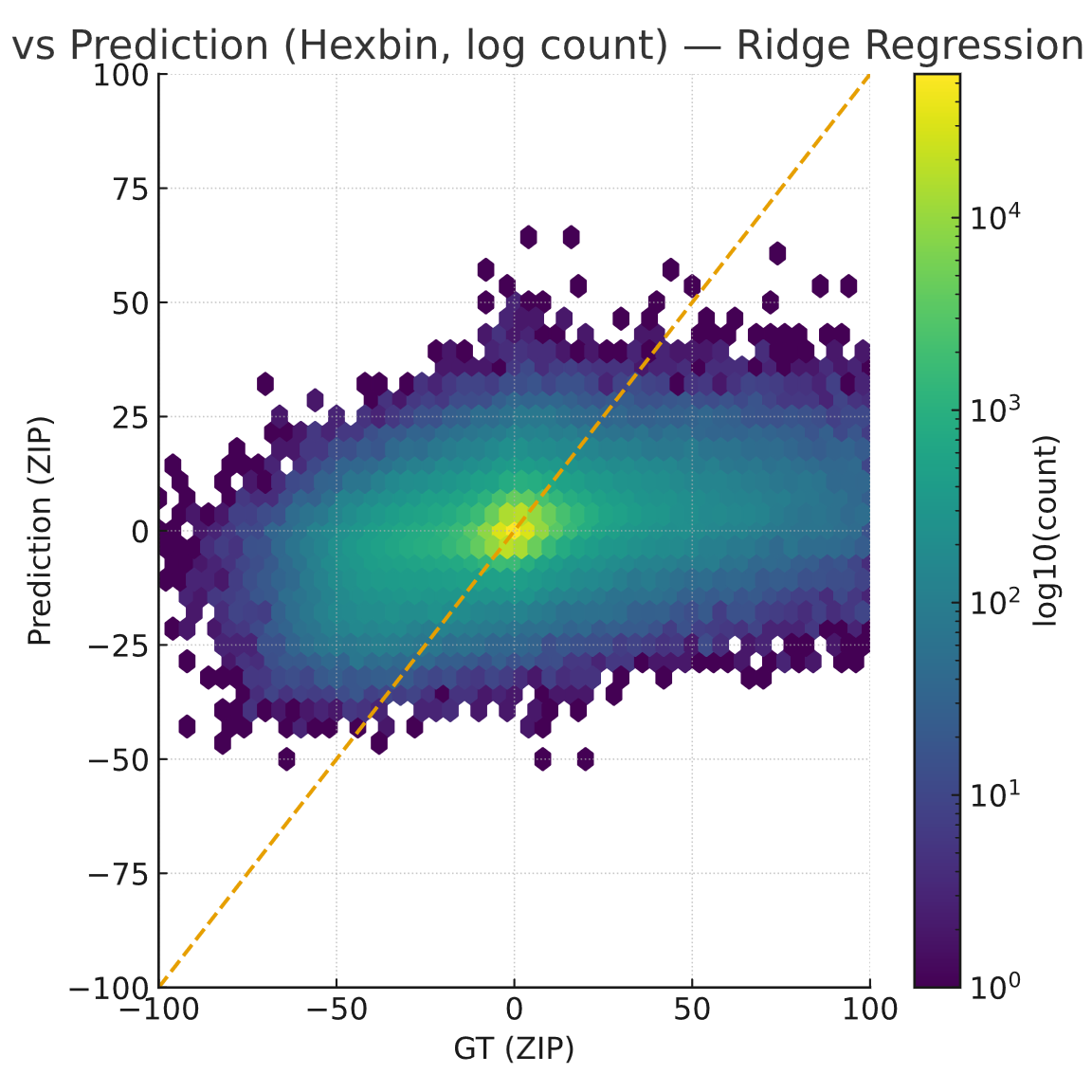

**a**

**b**

**c**

**Supplementary Figure 4.**

(a) Binned concordance between observed and predicted Bliss values by Ridge regression, (bin size = 5), color intensity indicating the percentage of samples within each bin.

(b) Binned error distribution (observed – predicted Bliss values by Ridge regression) showing deviation across the observed range.

(c) Hex-bin plot comparing observed versus predicted Bliss synergy scores by Ridge regression (log count).

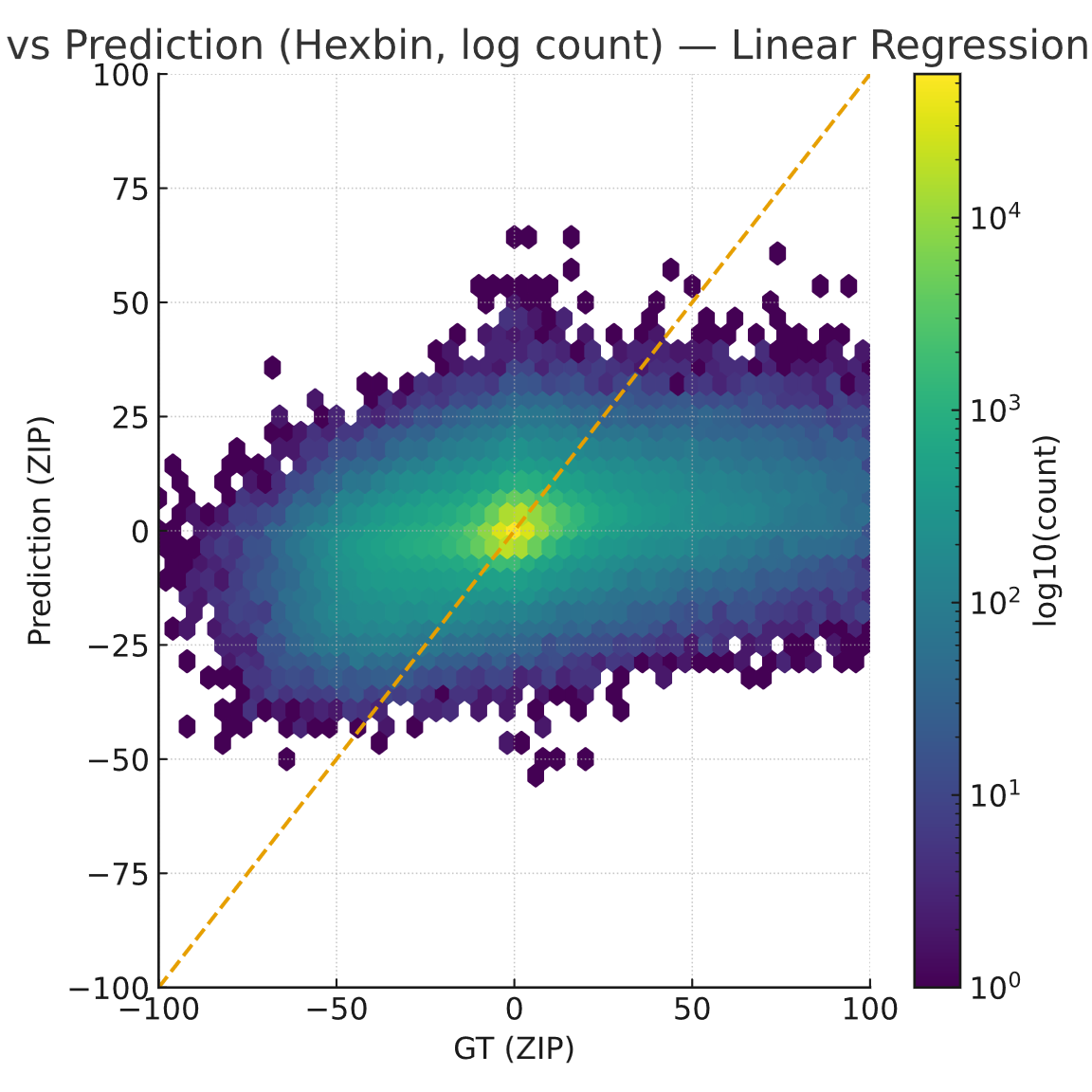

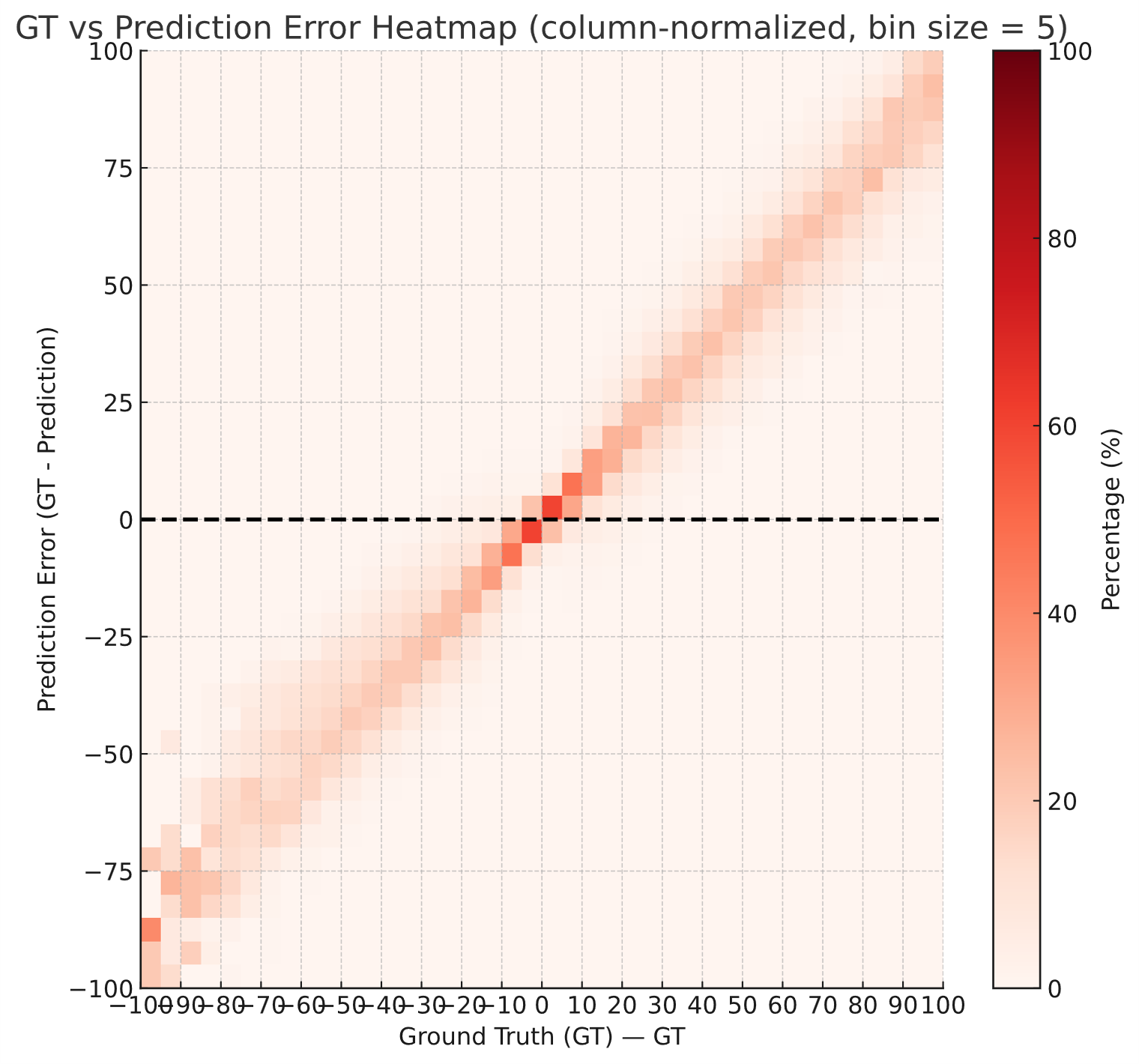

**b**

**c**

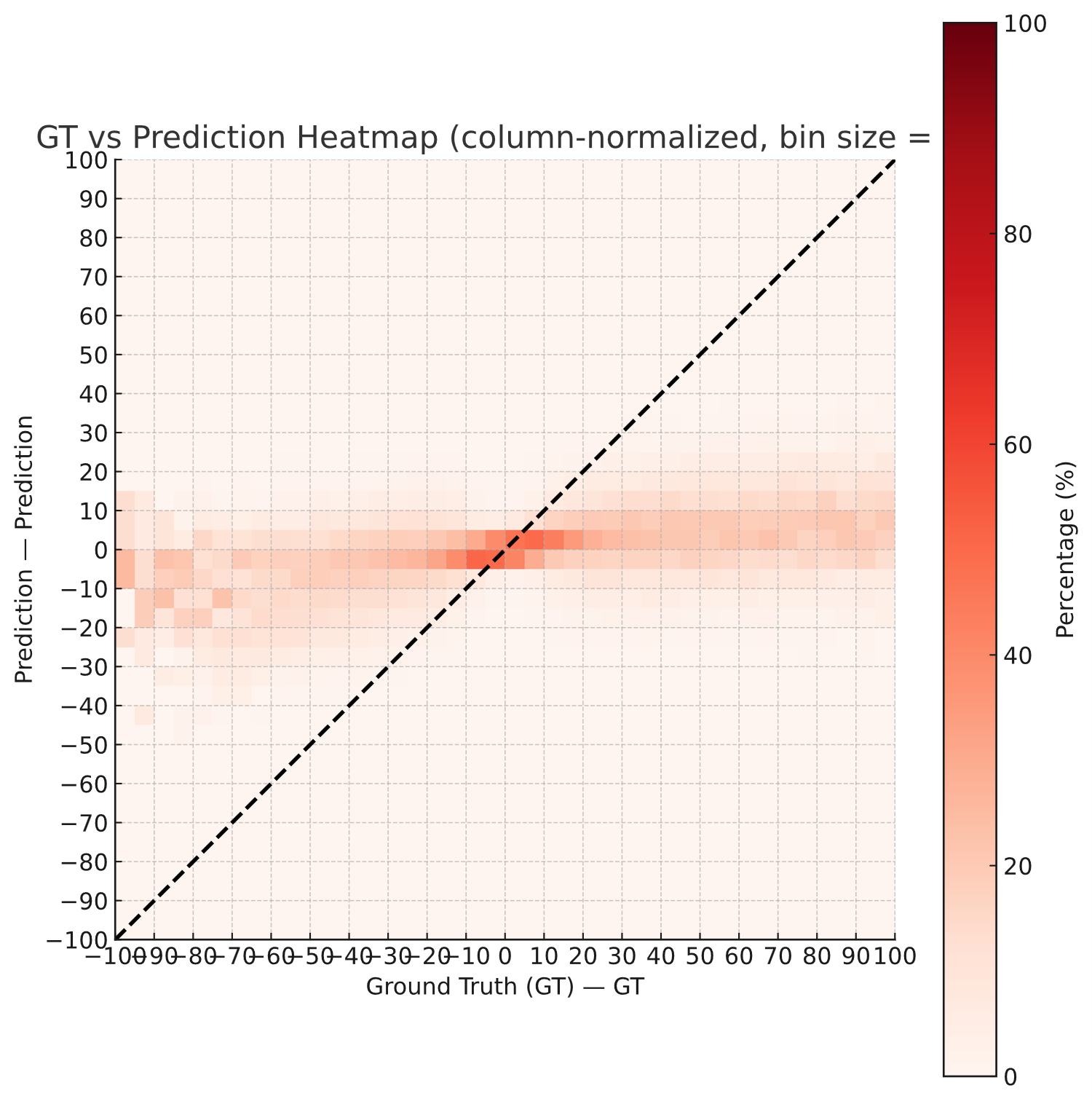

**a**

**Supplementary Figure 5.**

(a) Binned concordance between observed and predicted Bliss values by Linear regression, (bin size = 5), color intensity indicating the percentage of samples within each bin.

(b) Binned error distribution (observed – predicted Bliss values by Linear regression) showing deviation across the observed range.

(c) Hex-bin plot comparing observed versus predicted Bliss synergy scores by Linear regression (log count).

**c**

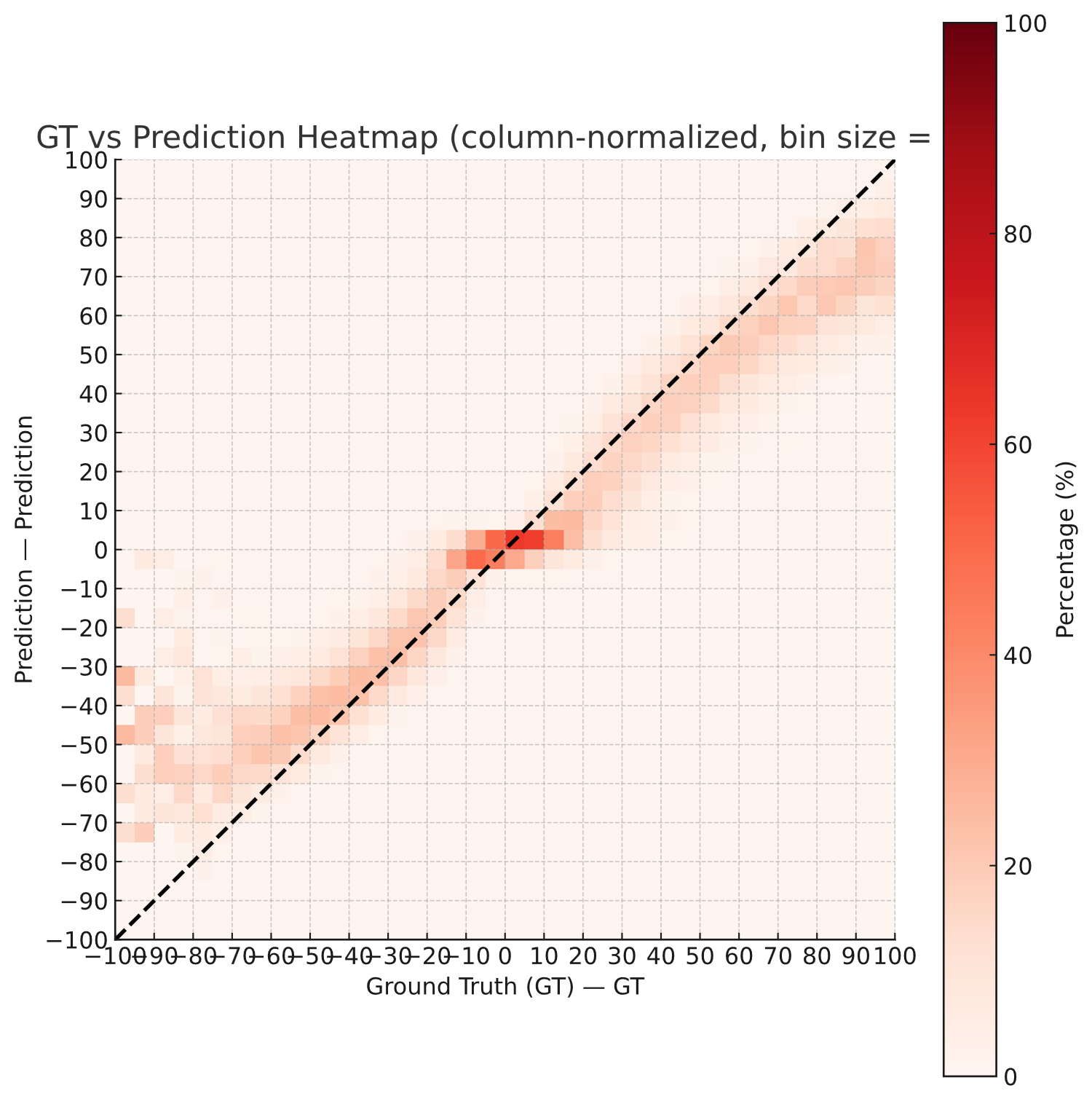

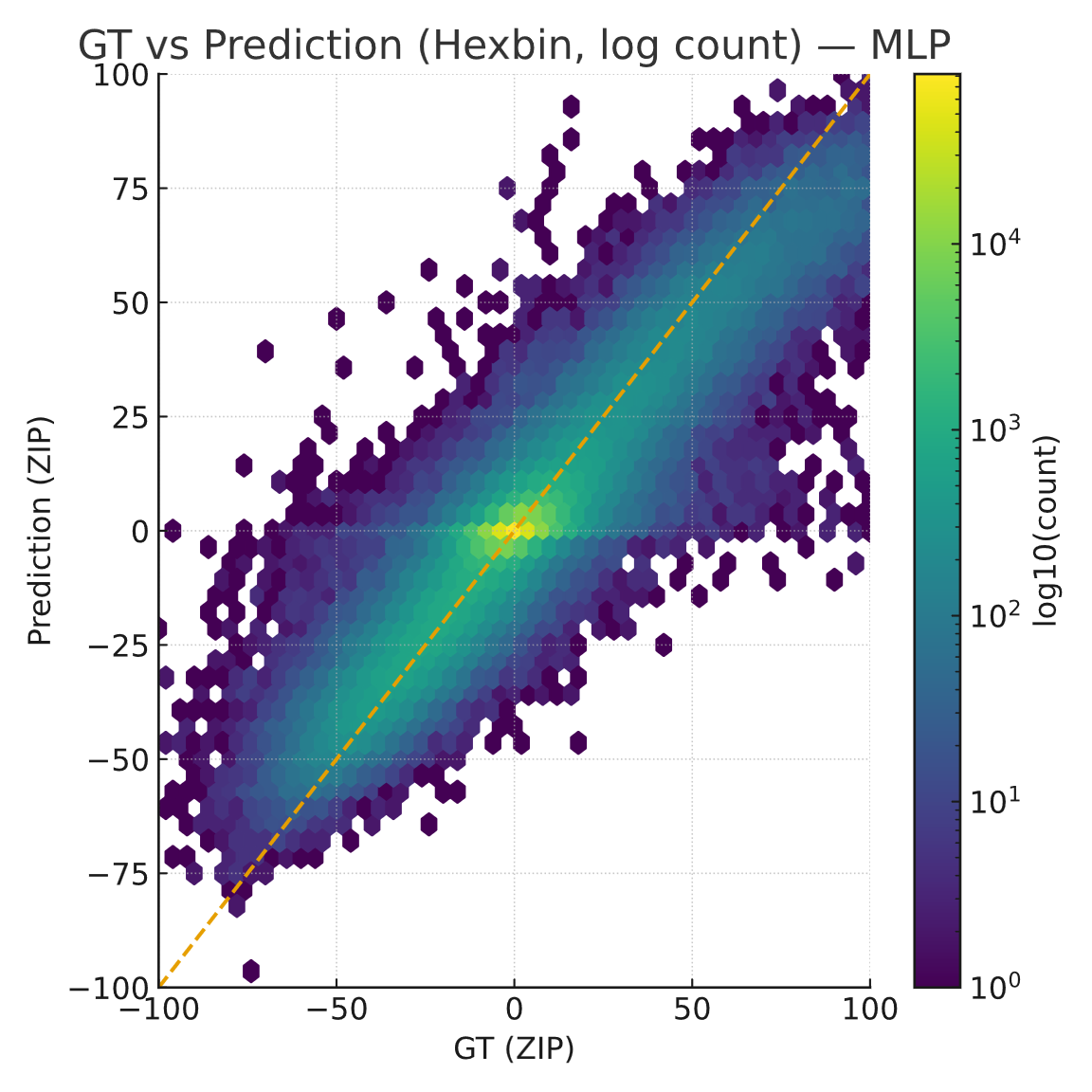

**a**

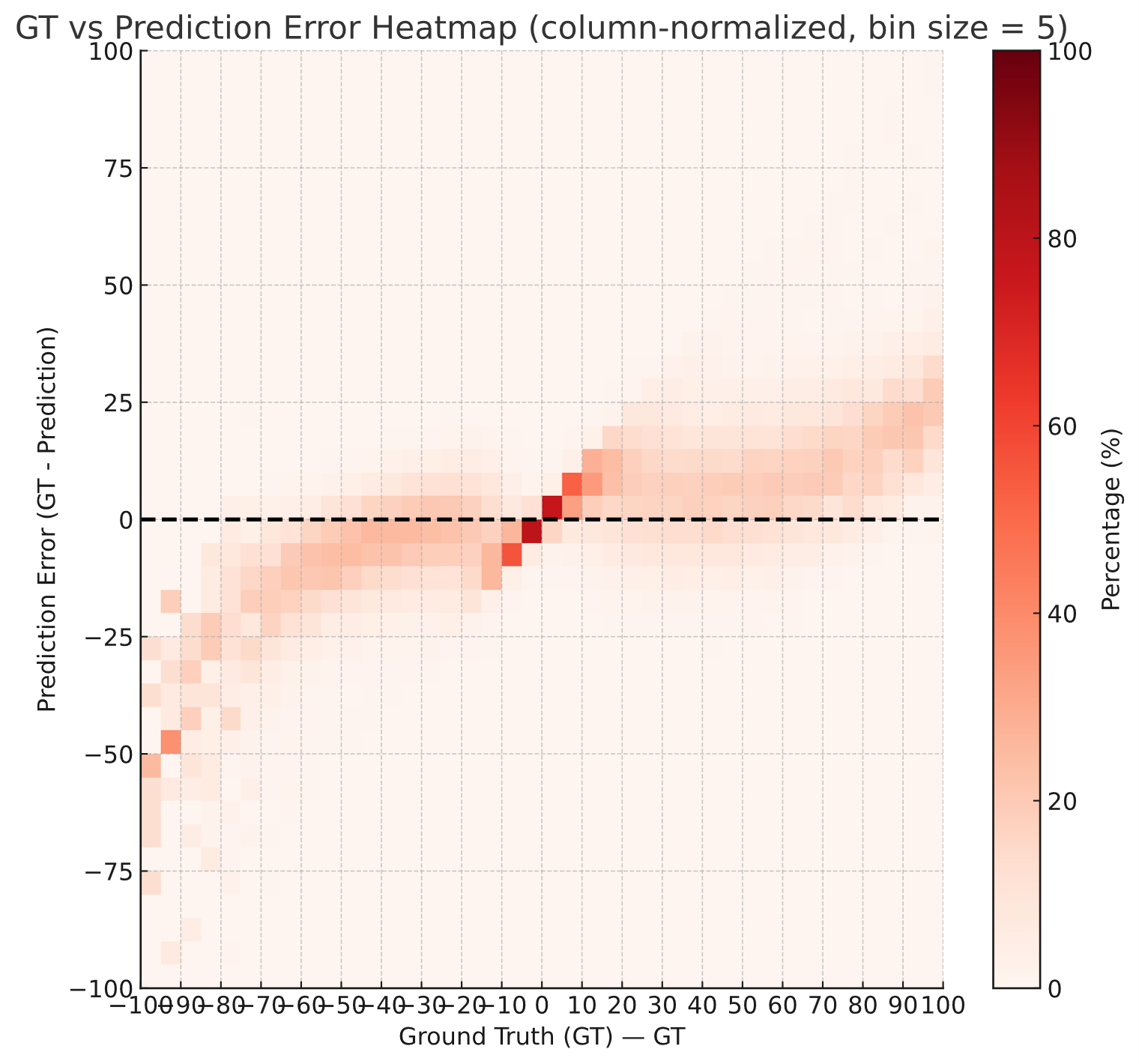

**b**

**Supplementary Figure 6.**

(a) Binned concordance between observed and predicted Bliss values by MLP, (bin size = 5), color intensity indicating the percentage of samples within each bin.

(b) Binned error distribution (observed – predicted Bliss values by MLP) showing deviation across the observed range.

(c) Hex-bin plot comparing observed versus predicted Bliss synergy scores by MLP (log count).

**c**

**b**

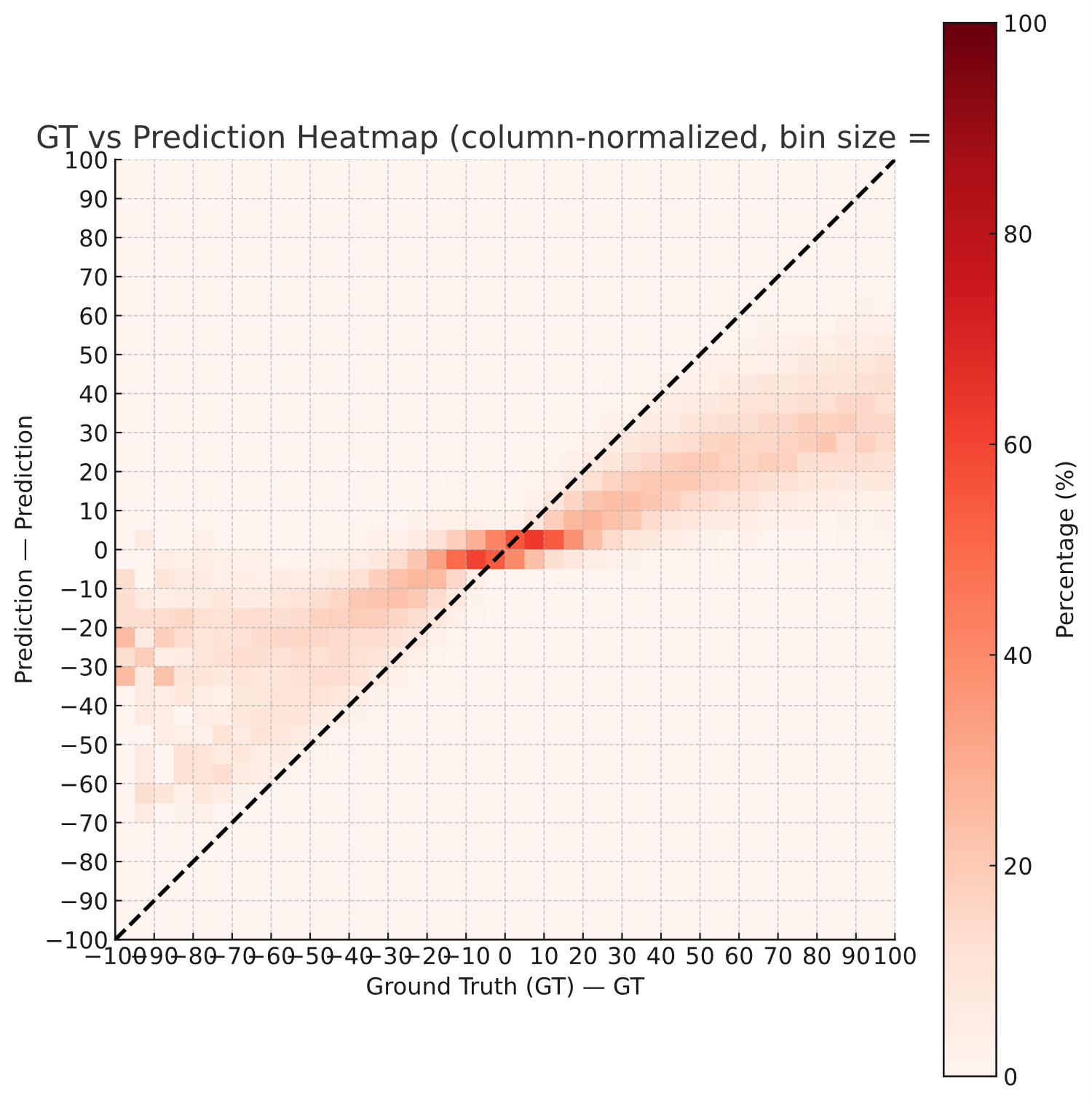

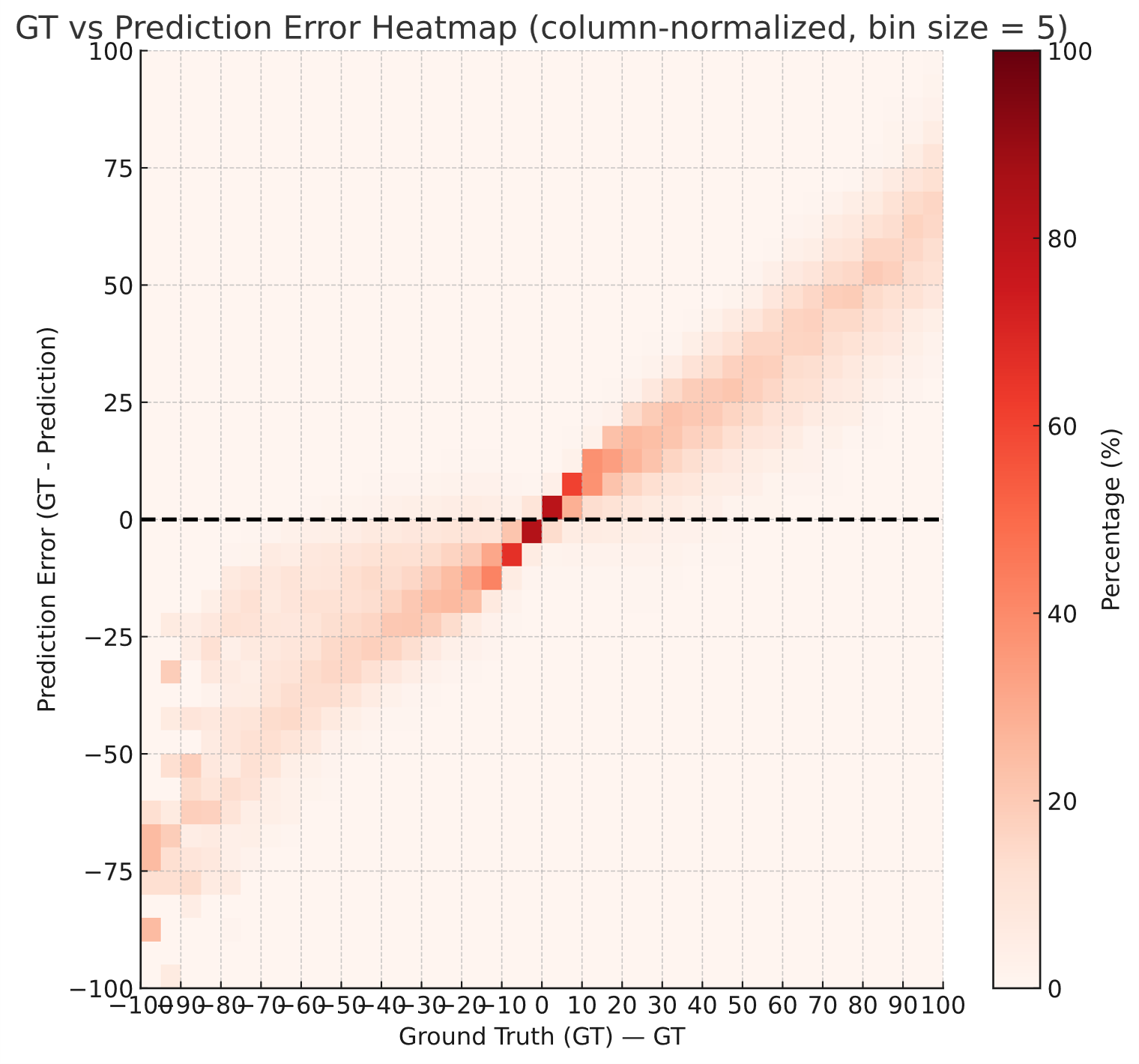

**a**

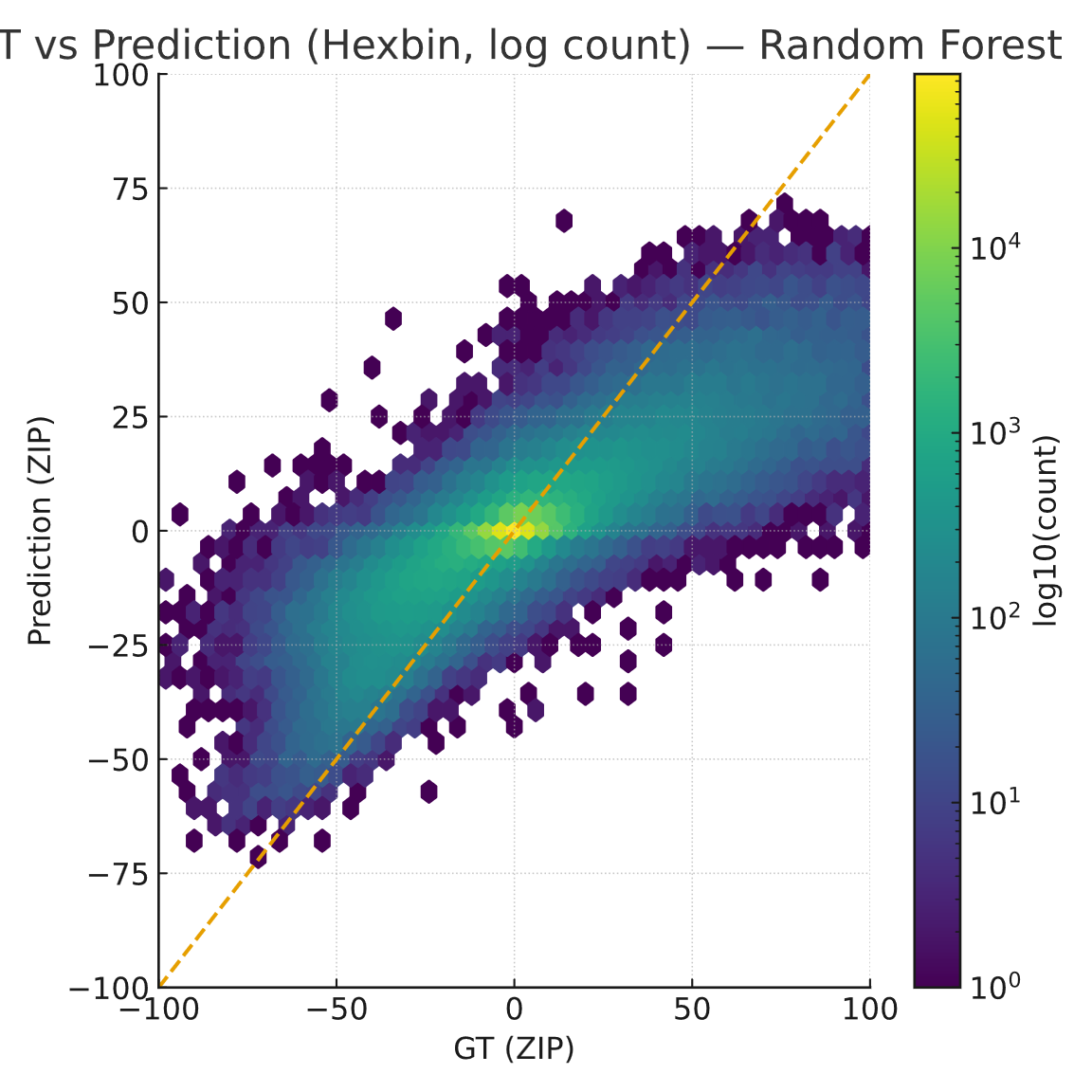

**Supplementary Figure 7.**

(a) Binned concordance between observed and predicted Bliss values by Random Forest, (bin size = 5), color intensity indicating the percentage of samples within each bin.

(b) Binned error distribution (observed – predicted Bliss values by Random Forest) showing deviation across the observed range.

(c) Hex-bin plot comparing observed versus predicted Bliss synergy scores by Random Forest (log count).

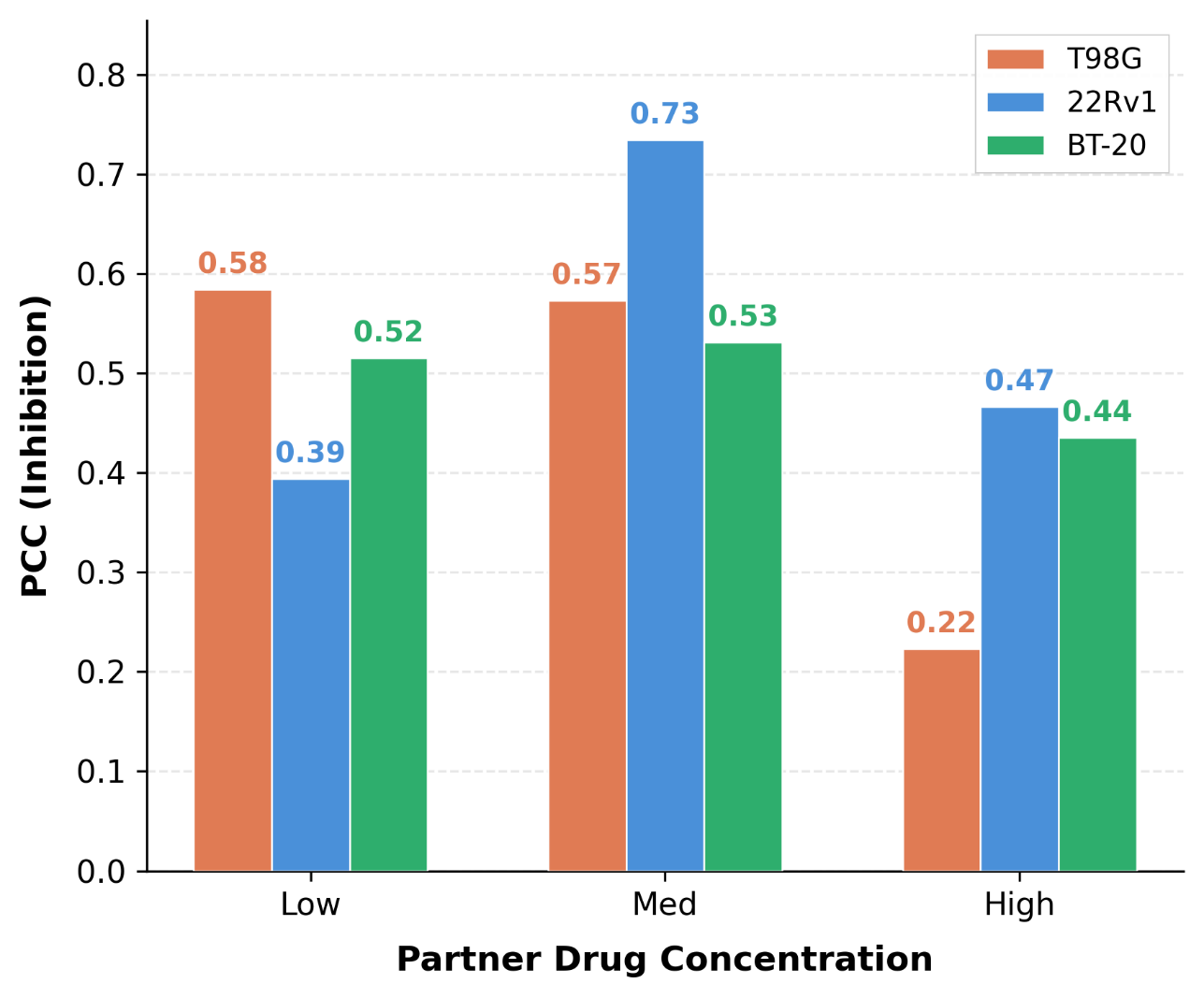

**Supplementary Figure 8 – PCC between predicted and observed inhibition stratified by partner drug concentration level (low: ≤0.5 µM; medium: 0.5–5.0 µM; high: >5.0 µM), computed separately for each cell line (T98G, 22Rv1, BT-20).**
