## Supplementary File 1:comparison of machine learning models, Supplementary File 2: MoA classes, Supplementary File 3: Prospective in vitro validation for "Uncertainty-Aware Deep Learning for Multi-Metric and Dose-Specific Prediction of Drug Synergy": Supplementary_Figures.pdf

**Supplementary Figure 1.** Experimental Data for HSA Synergy Measurement (Page 3)

**Supplementary Figure 2**. Experimental Data for ZIP Synergy Measurement (Page 5)

**Supplementary Figure 3.** Experimental Data for Loewe Synergy Measurement (Page 7)

**Supplementary Figure 4.** Experimental Results for Ridge Regression (Page 8)

**Supplementary Figure 5.** Experimental Results for Linear Regression (Page 9)

**Supplementary Figure 6.** Experimental Results for MLP (Page 10)

**Supplementary Figure 7.** Experimental Results for Random Forest (Page 11)

**Supplementary Figure 8.** PCC between predicted and observed inhibition stratified by partner drug concentration level (Page 12)

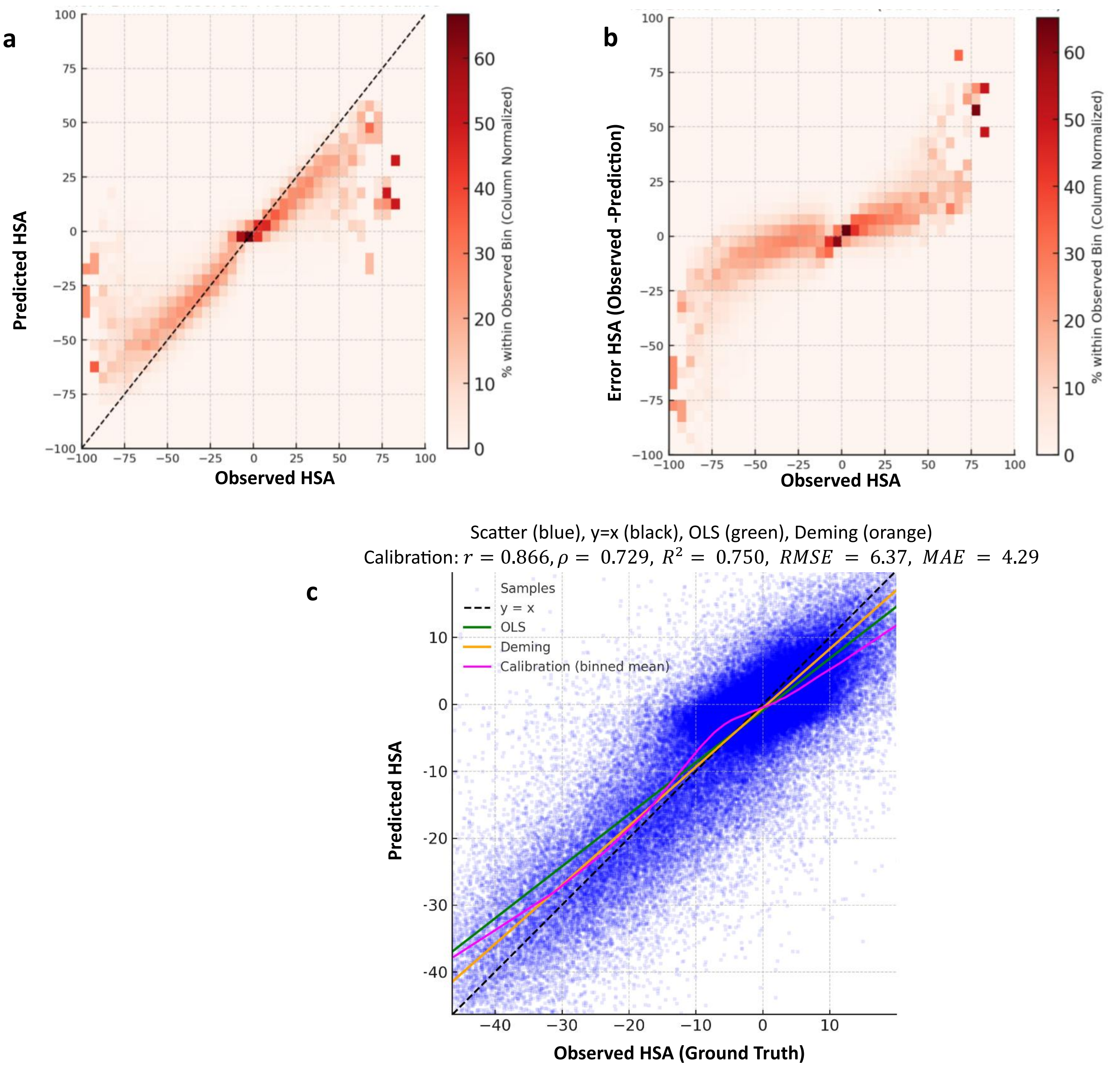
a
Predicted HSA
Observed HSA
% within Observed Bin (Column Normalized)
b
Error HSA (Observed -Prediction)
Observed HSA
% within Observed Bin (Column Normalized)
Scatter (blue), y=x (black), OLS (green), Deming (orange)
Calibration: $r = 0.866, \rho = 0.729, R^2 = 0.750, RMSE = 6.37, MAE = 4.29$
c
Samples
y = x
OLS
Deming
Calibration (binned mean)
Predicted HSA
Observed HSA (Ground Truth)

d

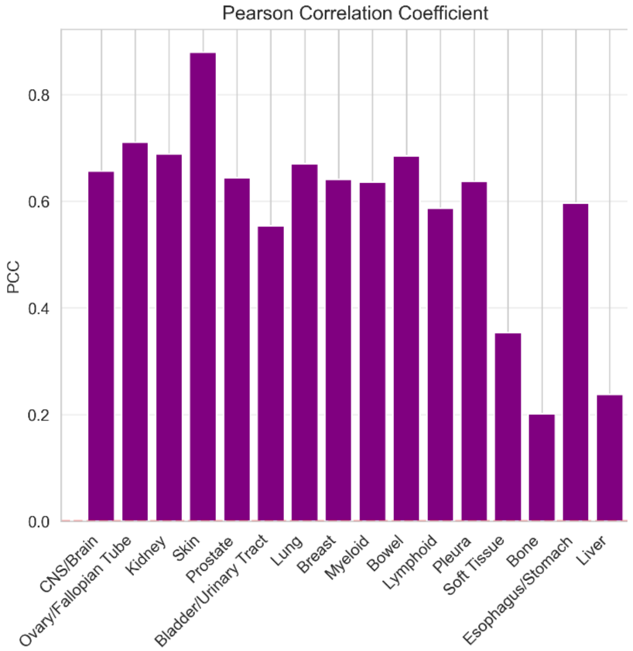


e

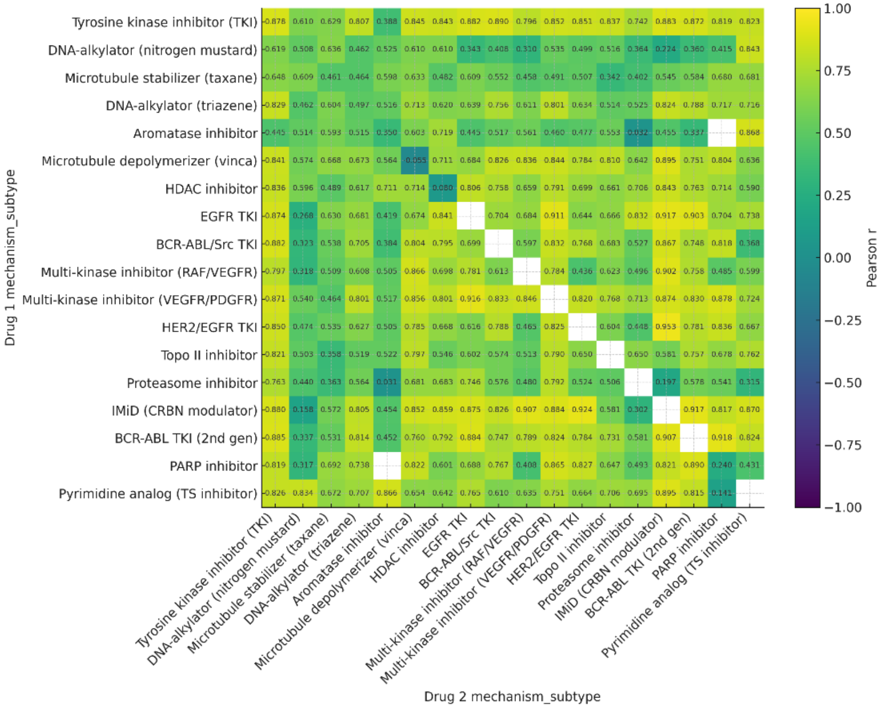


***Supplementary Figure 1.***

*(a) Binned concordance between observed and predicted HSA values (bin size = 5), with color intensity indicating the percentage of samples within each bin.*
*(b) Binned error distribution (observed – predicted HSA values) showing deviation across the observed range.*
*(c) Scatter plot of predicted versus observed HSA synergy scores across all test folds; the yellow line: the identity line (y = x), and the grey line: the fitted regression.*
*(d) Bar plot showing the number of observations per cancer tissue type (blue bars, log scale) and corresponding PCC between observed and predicted HSA.*
*(e) Heatmap showing the Pearson correlation coefficient between observed and predicted HSA for drug pairs grouped by their mechanism of action (MoA) class.*

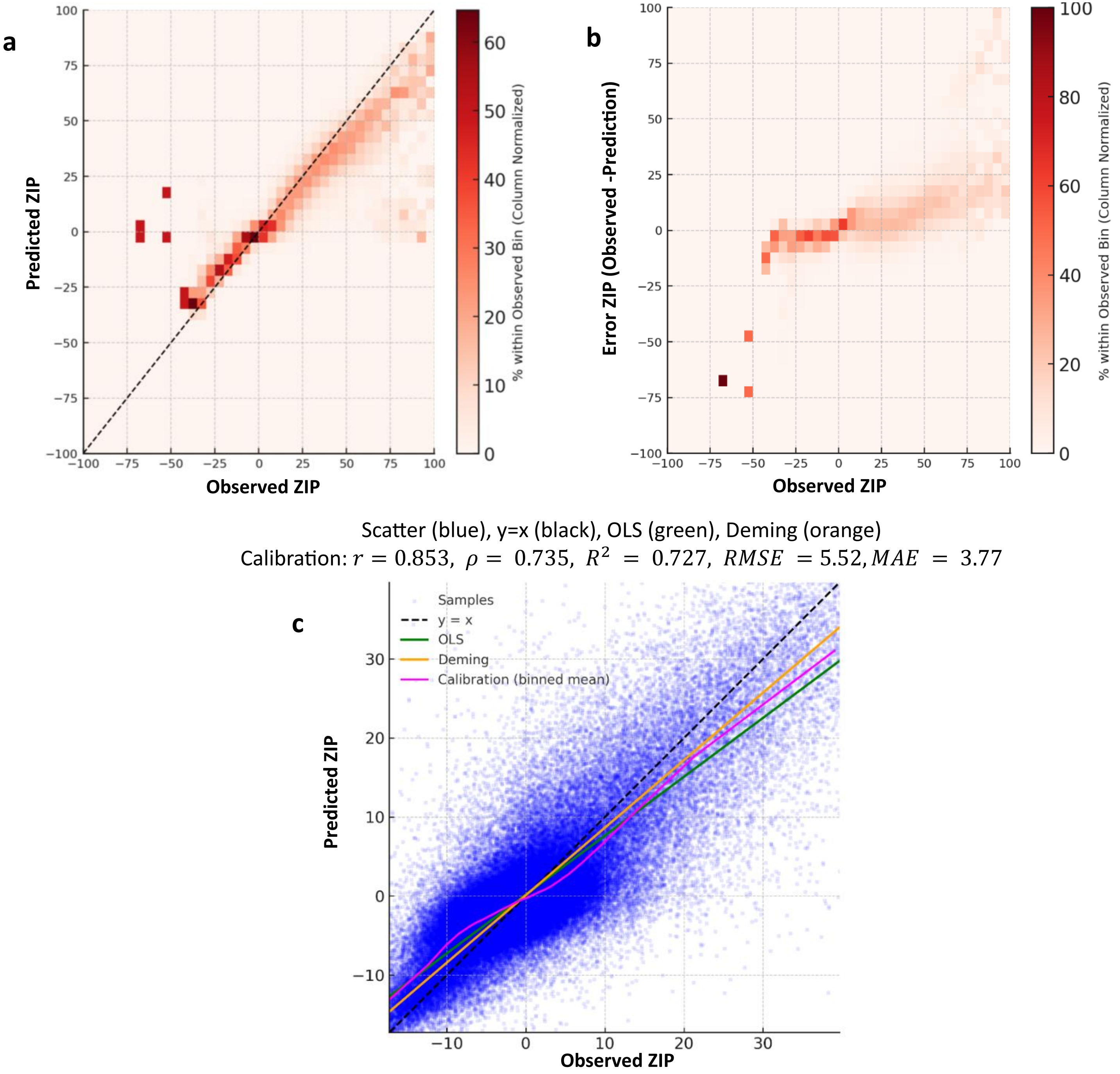
a
Predicted ZIP
Observed ZIP
% within Observed Bin (Column Normalized)
b
Error ZIP (Observed -Prediction)
Observed ZIP
% within Observed Bin (Column Normalized)
Scatter (blue), y=x (black), OLS (green), Deming (orange)
Calibration: $r = 0.853$, $\rho = 0.735$, $R^2 = 0.727$, $RMSE = 5.52$, $MAE = 3.77$
c
Samples
y = x
OLS
Deming
Calibration (binned mean)
Predicted ZIP
Observed ZIP

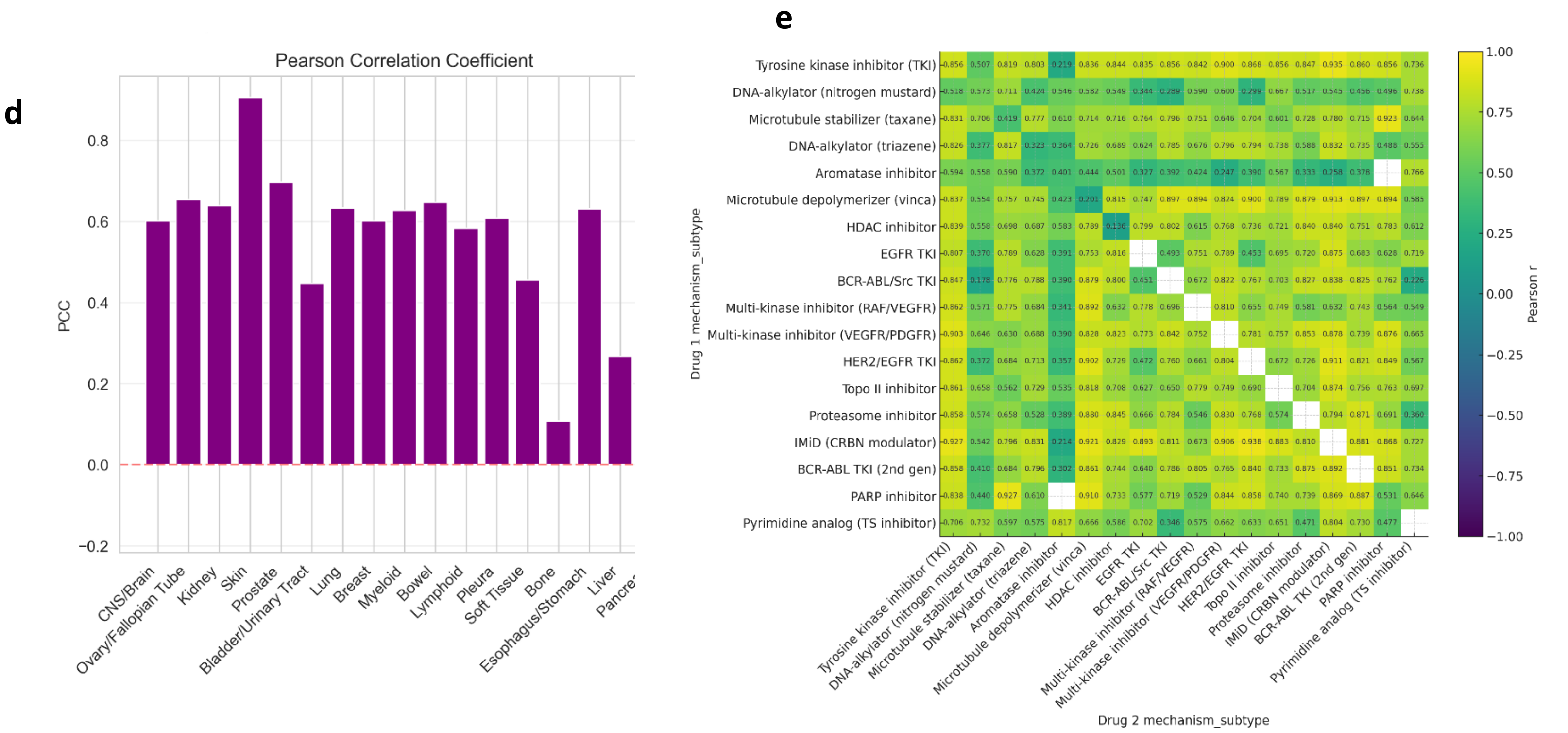


***Supplementary Figure 2.***

*(a)* *Binned concordance between observed and predicted ZIP values (bin size = 5), with color intensity indicating the percentage of samples within each bin.*
*(b)* *Binned error distribution (observed – predicted ZIP values) showing deviation across the observed range.*
*(c)* *Scatter plot of predicted versus observed ZIP synergy scores across all test folds; the yellow line: the identity line (y = x), and the grey line: the fitted regression.*
*(d)* *Bar plot showing the number of observations per cancer tissue type (blue bars, log scale) and corresponding PCC between observed and predicted ZIP.*
*(e)* *Heatmap showing the Pearson correlation coefficient between observed and predicted ZIP for drug pairs grouped by their mechanism of action (MoA) class.*

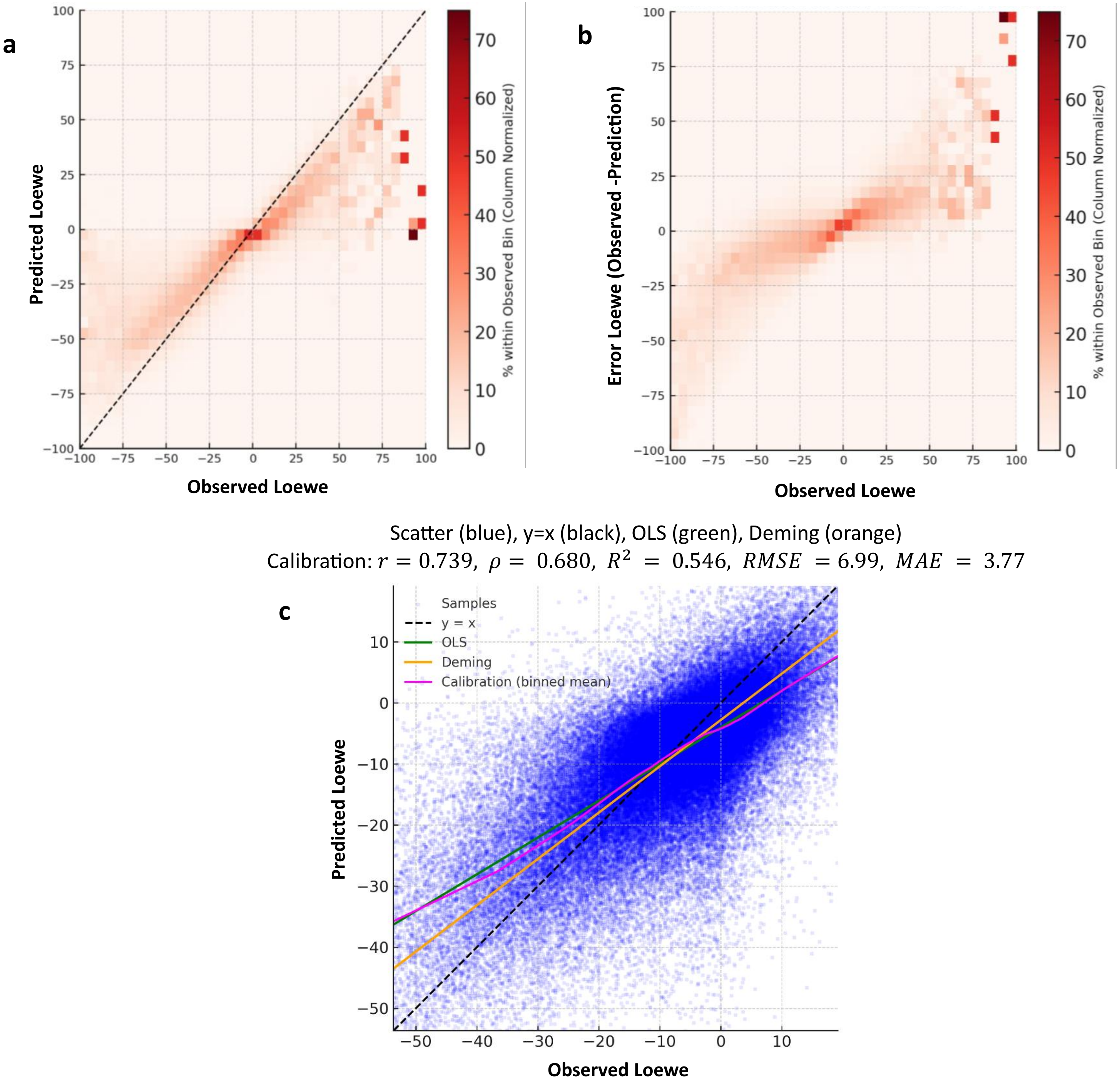
a
Predicted Loewe
Observed Loewe
% within Observed Bin (Column Normalized)
b
Error Loewe (Observed -Prediction)
Observed Loewe
% within Observed Bin (Column Normalized)
Scatter (blue), y=x (black), OLS (green), Deming (orange)
Calibration: $r = 0.739,\ \rho =\ 0.680,\ R^2\ =\ 0.546,\ RMSE\ = 6.99,\ MAE\ =\ 3.77$
c
Samples
y = x
OLS
Deming
Calibration (binned mean)
Predicted Loewe
Observed Loewe

d

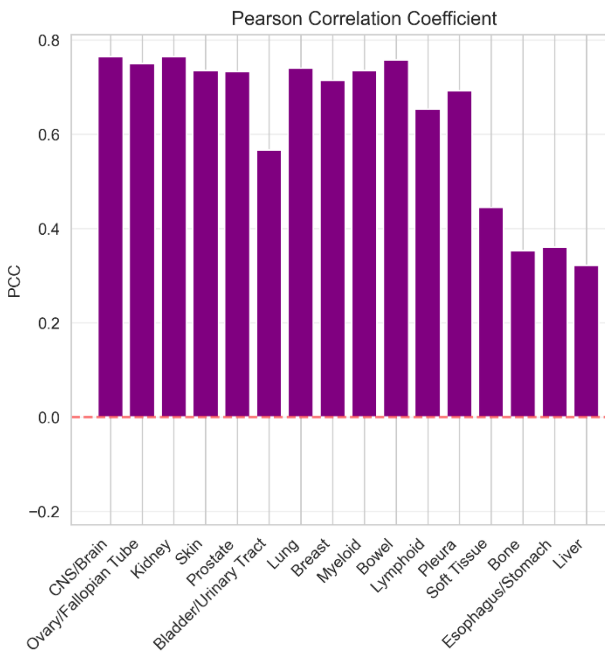


e

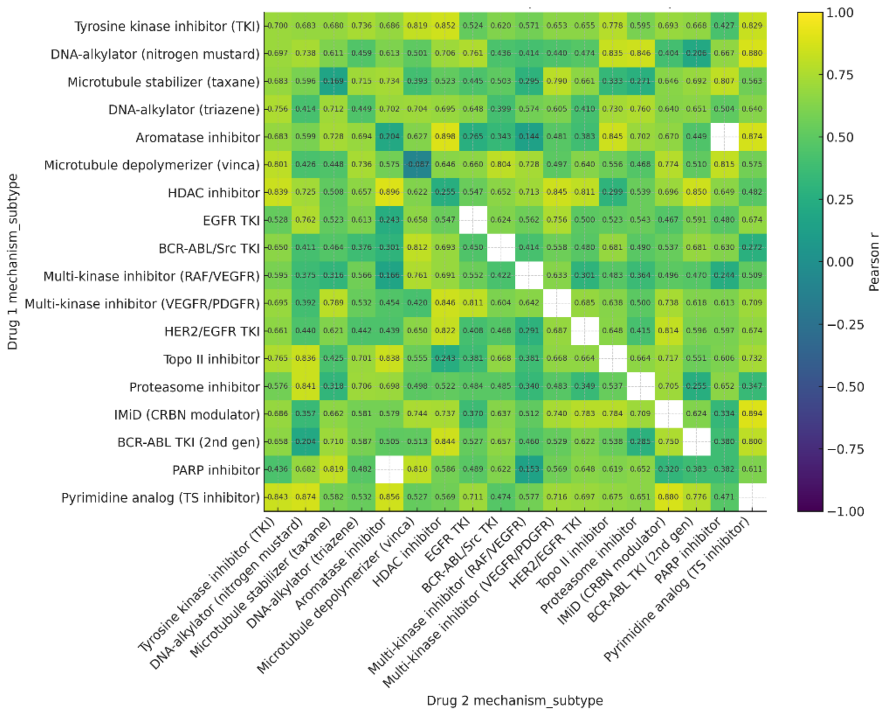


***Supplementary Figure 3.***

*(a) Binned concordance between observed and predicted Loewe values (bin size = 5), with color intensity indicating the percentage of samples within each bin.*
*(b) Binned error distribution (observed – predicted Loewe values) showing deviation across the observed range.*
*(c) Scatter plot of predicted versus observed Loewe synergy scores across all test folds; the yellow line: the identity line (y = x), and the grey line: the fitted regression.*
*(d) Bar plot showing the number of observations per cancer tissue type (blue bars, log scale) and corresponding PCC between observed and predicted Loewe.*
*(e) Heatmap showing the Pearson correlation coefficient between observed and predicted Loewe for drug pairs grouped by their mechanism of action (MoA) class.*

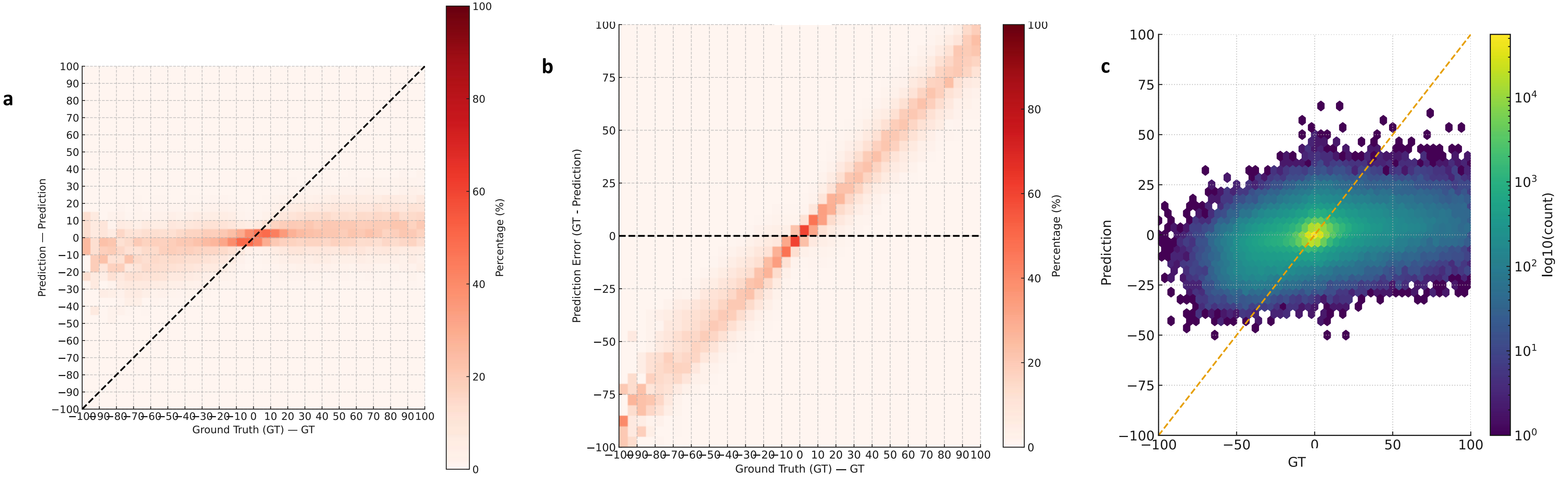


***Supplementary Figure 4.***

*(a) Binned concordance between observed and predicted Bliss values by Ridge regression, (bin size = 5), color intensity indicating the percentage of samples within each bin.*
*(b) Binned error distribution (observed – predicted Bliss values by Ridge regression) showing deviation across the observed range.*
*(c) Hex-bin plot comparing observed versus predicted Bliss synergy scores by Ridge regression (log count).*

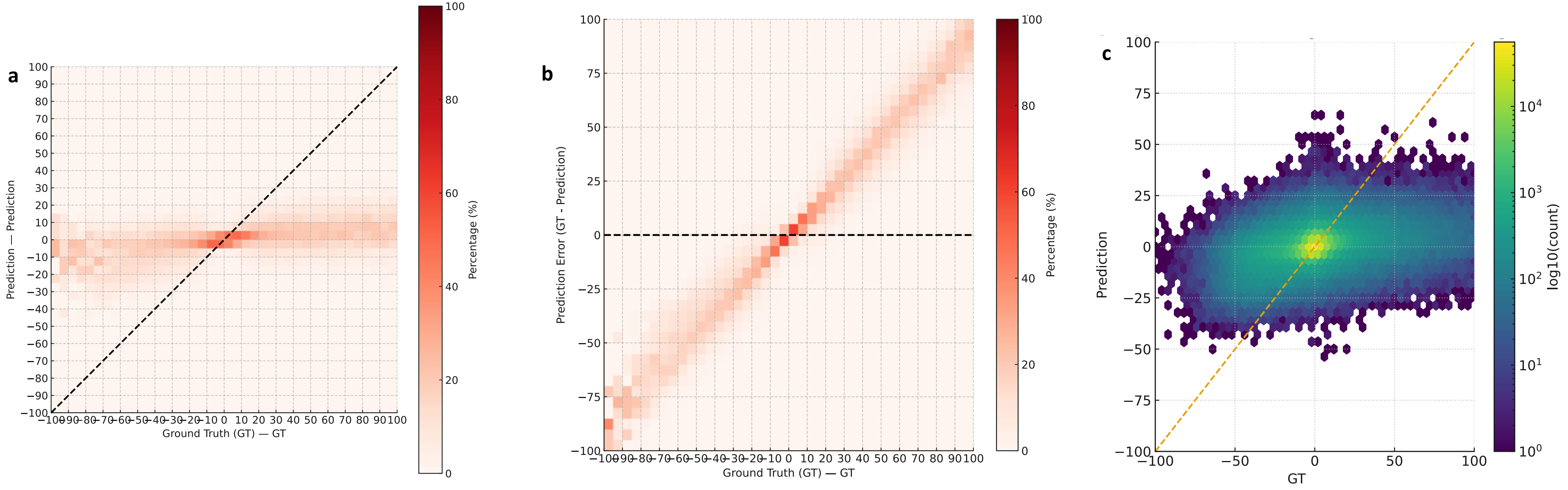


***Supplementary Figure 5.***

*(a) Binned concordance between observed and predicted Bliss values by Linear regression, (bin size = 5), color intensity indicating the percentage of samples within each bin.*
*(b) Binned error distribution (observed – predicted Bliss values by Linear regression) showing deviation across the observed range.*
*(c) Hex-bin plot comparing observed versus predicted Bliss synergy scores by Linear regression (log count).*

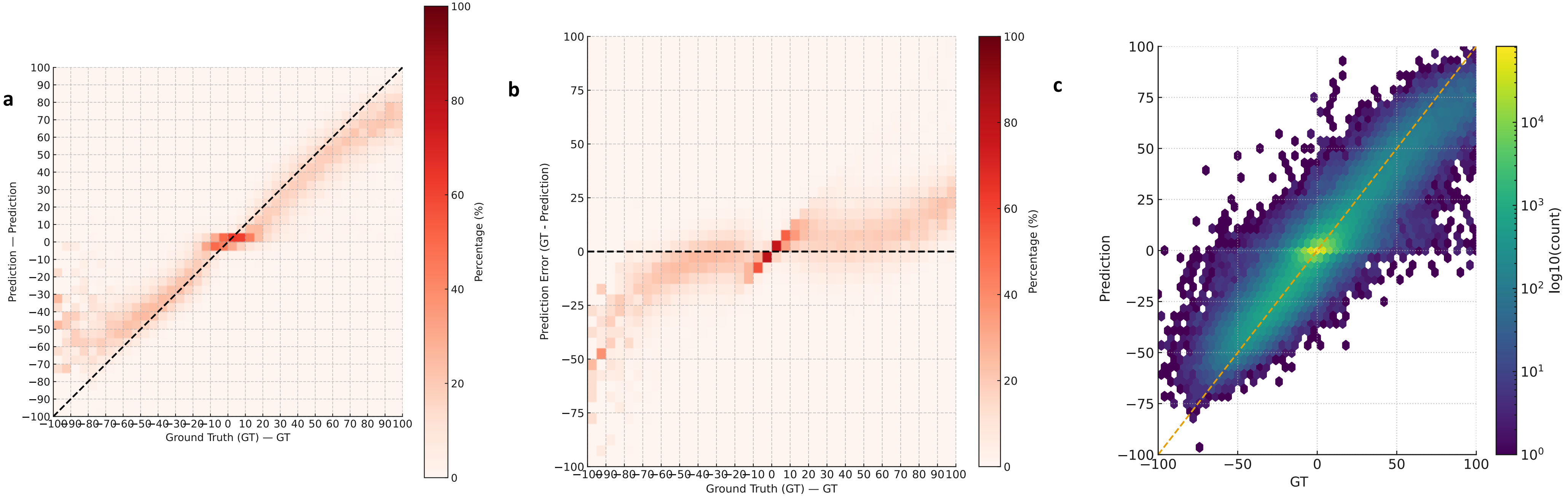


***Supplementary Figure 6.***

*(a) Binned concordance between observed and predicted Bliss values by MLP, (bin size = 5), color intensity indicating the percentage of samples within each bin.*
*(b) Binned error distribution (observed – predicted Bliss values by MLP) showing deviation across the observed range.*
*(c) Hex-bin plot comparing observed versus predicted Bliss synergy scores by MLP (log count).*

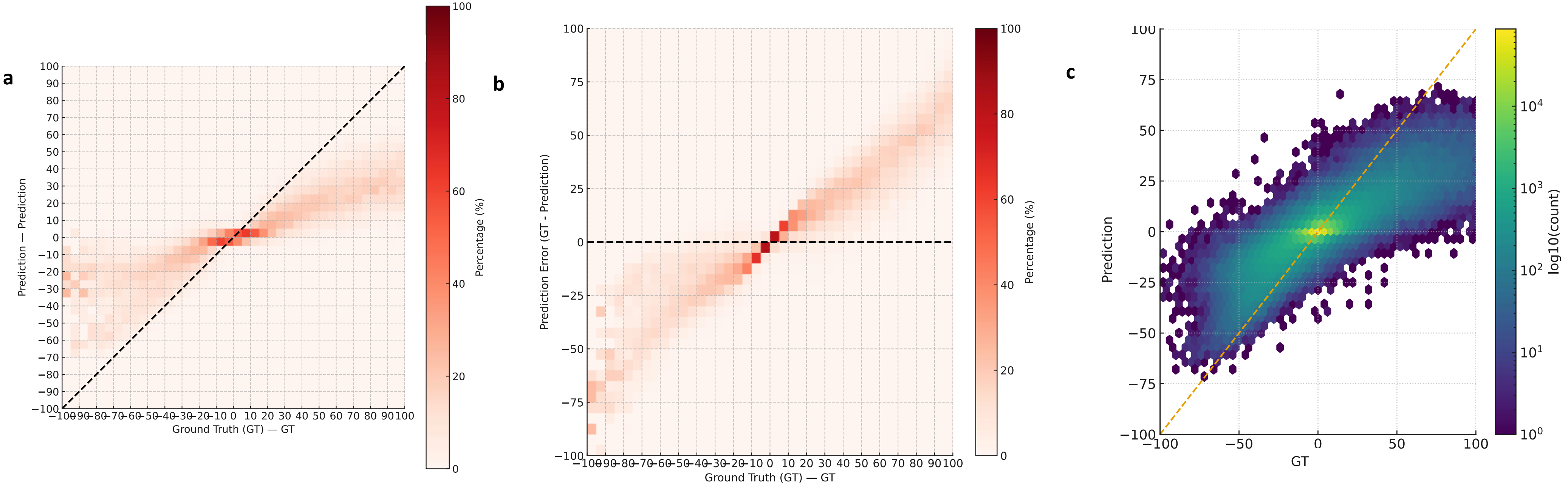


***Supplementary Figure 7.***

*(a) Binned concordance between observed and predicted Bliss values by Random Forest, (bin size = 5), color intensity indicating the percentage of samples within each bin.*
*(b) Binned error distribution (observed – predicted Bliss values by Random Forest) showing deviation across the observed range.*
*(c) Hex-bin plot comparing observed versus predicted Bliss synergy scores by Random Forest (log count).*

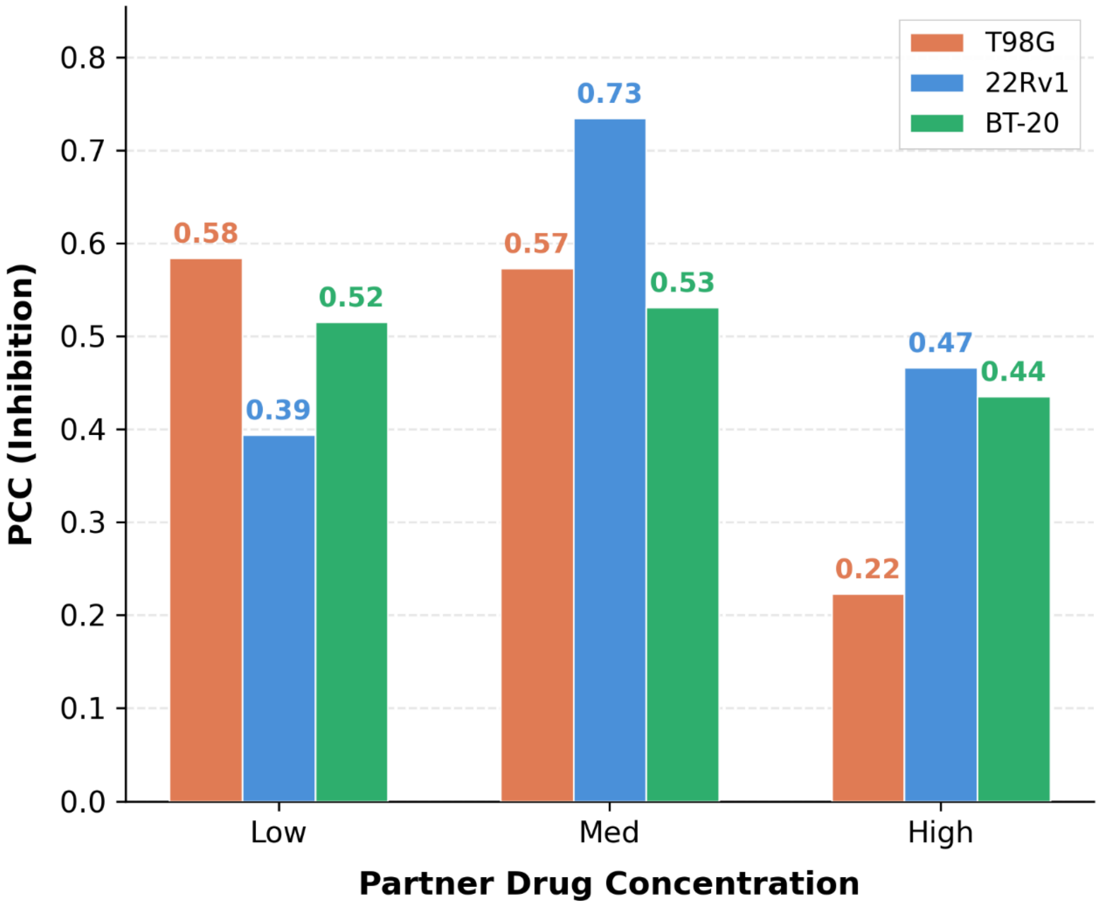


***Supplementary Figure 8 – PCC between predicted and observed inhibition stratified by partner drug concentration level (low: ≤0.5 μM; medium: 0.5–5.0 μM; high: >5.0 μM), computed separately for each cell line (T98G, 22Rv1, BT-20).***
