## Supplementary File 1:comparison of machine learning models, Supplementary File 2: MoA classes, Supplementary File 3: Prospective in vitro validation for "Uncertainty-Aware Deep Learning for Multi-Metric and Dose-Specific Prediction of Drug Synergy": Supplementary_Tables.docx

Professor Fatemeh Vafaee

School of Biotechnology and Biomolecular Sciences

UNSW SYDNEY NSW 2052 AUSTRALIA

T: +61 (2) 9065 2699

E:

**Supplementary Table 1.** Overview of 16 distinct molecular representation strategies (Page 2)

**Supplementary Table 2**. Drug representation ablation study for predictive performance (Page 4)

**Supplementary Table 1.** Overview of 16 distinct molecular representation strategies, encompassing expert-defined structural fingerprints, learned embeddings, physicochemical descriptors, and functional profiles that collectively capture chemical structure, bioactivity, and systems-level context.

| **No** | **Category / Family** | **What it captures** | **Typical data type & size** | **Similarity /distance measure** | **Representative members / examples** |
| --- | --- | --- | --- | --- | --- |
| **1** | **Key‑based structural fingerprints** | Presence/absence of expert‑defined SMARTS or fragment keys (fixed dictionary). Highly interpretable. | Binary vector (e.g. 166–5 000 bits) | Tanimoto,  Dice on bit sets | MACCS (166 bits), PubChem (881 bits), Klekota‑Roth (4860 bits), EState, Substructure (CDK / Open Babel FP4) |
| **2** | **Hashed topology fingerprints (path‑ or circular‑based)** | Local topological environments hashed into bit positions; variable collision; low‑level structure. | Binary vector (typically 1 024–4 096 bits) | Tanimoto,  Dice | ECFP / Morgan, RDKit Path FP, Daylight‑style FP |
| **3** | **Atom‑pair & torsion fingerprints** | Counts of atom‑type pairs at topological distances; or sequences of four bonded atoms (“topological torsions”). Capture medium‑range topology. | Count / binary vector (variable length or hashed to 1–4 k bits) | Tanimoto (binary or count), Cosine | Atom Pair FP, Topological Torsion FP |
| **4** | **Pharmacophore /**  **reduced‑graph fingerprints** | Encodes abstract pharmacophore features (H‑bond donors/acceptors, aromatic, hydrophobes) and their topology; reduced graphs compress rings/linkers. | Binary vector (hundreds–thousands of bits) | Tanimoto | ErG (Extended Reduced Graph), Topological Pharmacophore Triplets, Pharmacophore Keys |
| **5** | **Physicochemical /**  **global descriptor vectors** | Hand‑crafted numerical properties: constitutional, electronic, geometric, hydrophobic, etc.; summarize whole molecule. | Continuous vector (hundreds–thousands float, e.g. RDKit 200+, Dragon >3 000) | Euclidean, Cosine, Mahalanobis, Pearson | RDKit / Mordred descriptors, Dragon descriptors |
| **6** | **2‑D shape /**  **scaffold encodings** | Coarse representation of ring‑linker frameworks or shape signatures without full 3‑D conformers. | Strings / hashed bits | Levenshtein (scaffolds), Tanimoto | Bemis–Murcko scaffold strings, Scaffold keys |
| **7** | **Sequence‑based discrete encodings (SMILES)** | Treat SMILES as text: one‑hot tokens, k‑mer / n‑gram counts, unsupervised word embeddings. Capture syntax motifs. | Sparse counts or dense vectors (e.g. 300‑dim Mol2Vec) | Cosine, Euclidean | Bag‑of‑n‑grams, Mol2Vec, SMILES2Vec |
| **8** | **Self‑supervised SMILES language‑model embeddings** | Contextual embeddings learned by transformers or masked‑LMs on large SMILES corpora; encode global chemistry & subtle semantics. | Dense vector (hundreds–thousands dim) | Cosine, Euclidean | ChemBERTa, SMILES Transformer, MolT5, MoLFormer |
| **9** | **Graph‑structured (2‑D) learned representations** | Full molecular graph (atoms/bonds) enabling message passing / neural embeddings; captures arbitrary substructures. | Node‑feature matrix + edge list (variable); learned fixed vectors after pooling | GNN embeddings compared by Cosine / Euclidean; kernel methods | Message Passing Neural Networks (MPNN), GraphConv, GIN |
| **10** | **3‑D geometric descriptors (hand‑crafted)** | Geometry of a conformer: pairwise Coulombic interactions, distances, local atomic environments. | Matrices / vectors (e.g. Coulomb matrix n×n, Bag‑of‑Bonds, SOAP vector) | Frobenius norm, RMSD, Cosine, Gaussian kernels | Coulomb Matrix, Bag‑of‑Bonds, SOAP |
| **11** | **3‑D equivariant neural embeddings** | Learn from 3‑D coordinates with rotational/translation equivariance; capture quantum‑level interactions. | Learned dense vector (hundreds dims) | Cosine / task‑specific loss | SchNet, DimeNet / DimeNet++, PhysNet, EGNN |
| **12** | **3‑D shape & pharmacophore alignment descriptors** | Conformer shape overlaps or pharmacophore alignment for virtual screening. | Real‑valued shape/pharmacophore feature vectors | Shape / color Tanimoto, Gaussian overlap | ROCS Shape/Color, Ultrafast Shape Recognition (USR), SHAEP |
| **13** | **Bioactivity /**  **target‑profile fingerprints** | Experimental response profiles across assays or targets reflect functional similarity beyond structure. | Continuous or binary vector (assays × responses) | Pearson, Cosine, Tanimoto (binary) | HTSFP (High‑Throughput Screening Fingerprints), Target inhibition profiles |
| **14** | **Knowledge‑based /**  **omics‑linked features** | External knowledge: drug–target gene sets, pathways, side‑effect vectors; integrate systems biology context. | Binary / count vectors (genes, pathways) | Jaccard, Cosine | Target gene binary vectors, Pathway membership, Side‑effect (e.g., SIDER) |
| **15** | **Image‑based (2‑D rendering) representations** | Rasterized 2‑D depictions processed by CNNs; capture visual layout of substructures. | Image tensor (e.g. 224×224×3) | Learned CNN embedding (Cosine) | CNN on RDKit images, Vision Transformers over 2‑D depictions |
| **16** | **Hybrid multimodal models** | Joint embedding of text (natural language or SMILES) + graph/3‑D modalities; leverage complementary information. | Multiple encoders; fused dense vector | Cosine / task‑specific | MolT5 (text–SMILES), graph‑text alignment (e.g., MoMu/MolCA) |

**Supplementary Table 2** Results of the ablation study quantifying the contribution of each drug representation to predictive performance across synergy measurements and evaluation metrics.

| Input Drug Features | Synergy Measurement | Performance Metric | Value |
| --- | --- | --- | --- |
| Chemberta | ZIP | MAE | 6.238492803193105 |
| Chemberta | Loewe | MAE | 4.664775085850207 |
| Chemberta | Bliss | MAE | 8.884453108838443 |
| Chemberta | HSA | MAE | 5.340179484 |
| Morgan-PubChem | ZIP | MAE | 6.130076541331025 |
| Morgan-PubChem | Loewe | MAE | 4.605825842401649 |
| Morgan-PubChem | Bliss | MAE | 8.733015494 |
| Morgan-PubChem | HSA | MAE | 5.335403700969333 |
| Morgan | ZIP | MAE | 6.161484734376237 |
| Morgan | Loewe | MAE | 4.627652246345203 |
| Morgan | Bliss | MAE | 8.726120358403165 |
| Morgan | HSA | MAE | 5.326292324996855 |
| PubChem | ZIP | MAE | 6.398809742676099 |
| PubChem | Loewe | MAE | 4.718590865225017 |
| PubChem | Bliss | MAE | 8.805984333442535 |
| PubChem | HSA | MAE | 5.433441848330955 |
| RDKit-Chemberta | ZIP | MAE | 6.510318540568121 |
| RDKit-Chemberta | Loewe | MAE | 4.733677356 |
| RDKit-Chemberta | Bliss | MAE | 8.794982479699703 |
| RDKit-Chemberta | HSA | MAE | 5.395819086602245 |
| RDKit-Chemberta-Morgan-PubChem | ZIP | MAE | 6.130076541331025 |
| RDKit-Chemberta-Morgan-PubChem | Loewe | MAE | 4.605825842401649 |
| RDKit-Chemberta-Morgan-PubChem | Bliss | MAE | 8.733015494 |
| RDKit-Chemberta-Morgan-PubChem | HSA | MAE | 5.335403700969333 |
| RDKit | ZIP | MAE | 6.222547540577302 |
| RDKit | Loewe | MAE | 4.656751530771261 |
| RDKit | Bliss | MAE | 8.957658109745008 |
| RDKit | HSA | MAE | 5.404925430731974 |
| RDKit-Morgan | ZIP | MAE | 6.307949994224713 |
| RDKit-Morgan | Loewe | MAE | 4.678861703423138 |
| RDKit-Morgan | Bliss | MAE | 8.810125342628776 |
| RDKit-Morgan | HSA | MAE | 5.365817480153674 |
| Chemberta | ZIP | MSE | 93.95728677165995 |
| Chemberta | Loewe | MSE | 44.95246613563758 |
| Chemberta | Bliss | MSE | 152.5237838678744 |
| Chemberta | HSA | MSE | 63.44304192420236 |
| Morgan-PubChem | ZIP | MSE | 92.26964371801554 |
| Morgan-PubChem | Loewe | MSE | 44.46219240053394 |
| Morgan-PubChem | Bliss | MSE | 152.2653367532283 |
| Morgan-PubChem | HSA | MSE | 63.26184958755129 |
| Morgan | ZIP | MSE | 92.92417340322895 |
| Morgan | Loewe | MSE | 44.726019808212975 |
| Morgan | Bliss | MSE | 150.71549675616325 |
| Morgan | HSA | MSE | 63.50720299098178 |
| PubChem | ZIP | MSE | 103.67078771082755 |
| PubChem | Loewe | MSE | 47.22643934038412 |
| PubChem | Bliss | MSE | 153.3045635108018 |
| PubChem | HSA | MSE | 66.55499557671311 |
| RDKit-Chemberta | ZIP | MSE | 108.66848822189108 |
| RDKit-Chemberta | Loewe | MSE | 49.02110267652446 |
| RDKit-Chemberta | Bliss | MSE | 156.95529390261552 |
| RDKit-Chemberta | HSA | MSE | 66.39045009 |
| RDKit-Chemberta-Morgan-PubChem | ZIP | MSE | 92.26964371801554 |
| RDKit-Chemberta-Morgan-PubChem | Loewe | MSE | 44.46219240053394 |
| RDKit-Chemberta-Morgan-PubChem | Bliss | MSE | 152.2653367532283 |
| RDKit-Chemberta-Morgan-PubChem | HSA | MSE | 63.26184958755129 |
| RDKit | ZIP | MSE | 94.52904921911608 |
| RDKit | Loewe | MSE | 45.27463338759316 |
| RDKit | Bliss | MSE | 155.6390845265861 |
| RDKit | HSA | MSE | 65.29703934135127 |
| RDKit-Morgan | ZIP | MSE | 99.76873095 |
| RDKit-Morgan | Loewe | MSE | 46.294371194658645 |
| RDKit-Morgan | Bliss | MSE | 153.1533271796862 |
| RDKit-Morgan | HSA | MSE | 65.21595354323553 |
| Chemberta | ZIP | RMSE | 9.693156697983374 |
| Chemberta | Loewe | RMSE | 6.704660031324301 |
| Chemberta | Bliss | RMSE | 12.350051978347071 |
| Chemberta | HSA | RMSE | 7.9651140559443565 |
| Morgan-PubChem | ZIP | RMSE | 9.605708913 |
| Morgan-PubChem | Loewe | RMSE | 6.667997630513521 |
| Morgan-PubChem | Bliss | RMSE | 12.339584140206197 |
| Morgan-PubChem | HSA | RMSE | 7.953731802591239 |
| Morgan | ZIP | RMSE | 9.639718533402775 |
| Morgan | Loewe | RMSE | 6.687751476259638 |
| Morgan | Bliss | RMSE | 12.276623996692383 |
| Morgan | HSA | RMSE | 7.969140668289259 |
| PubChem | ZIP | RMSE | 10.181885272916189 |
| PubChem | Loewe | RMSE | 6.872149542929353 |
| PubChem | Bliss | RMSE | 12.38162200645787 |
| PubChem | HSA | RMSE | 8.158124513435249 |
| RDKit-Chemberta | ZIP | RMSE | 10.424417884078279 |
| RDKit-Chemberta | Loewe | RMSE | 7.001507171782691 |
| RDKit-Chemberta | Bliss | RMSE | 12.528179991627496 |
| RDKit-Chemberta | HSA | RMSE | 8.148033510800499 |
| RDKit-Chemberta-Morgan-PubChem | ZIP | RMSE | 9.605708913 |
| RDKit-Chemberta-Morgan-PubChem | Loewe | RMSE | 6.667997630513521 |
| RDKit-Chemberta-Morgan-PubChem | Bliss | RMSE | 12.339584140206197 |
| RDKit-Chemberta-Morgan-PubChem | HSA | RMSE | 7.953731802591239 |
| RDKit | ZIP | RMSE | 9.722605063413615 |
| RDKit | Loewe | RMSE | 6.728642759694793 |
| RDKit | Bliss | RMSE | 12.475539448319903 |
| RDKit | HSA | RMSE | 8.080658348262922 |
| RDKit-Morgan | ZIP | RMSE | 9.988429854 |
| RDKit-Morgan | Loewe | RMSE | 6.803996707425617 |
| RDKit-Morgan | Bliss | RMSE | 12.375513208739514 |
| RDKit-Morgan | HSA | RMSE | 8.075639512957196 |
| Chemberta | ZIP | PCC | 0.8479875238745817 |
| Chemberta | Loewe | PCC | 0.7677186391385866 |
| Chemberta | Bliss | PCC | 0.5945700678093746 |
| Chemberta | HSA | PCC | 0.7860910395508683 |
| Morgan-PubChem | ZIP | PCC | 0.8510537790244346 |
| Morgan-PubChem | Loewe | PCC | 0.7699472078682355 |
| Morgan-PubChem | Bliss | PCC | 0.5959768915754906 |
| Morgan-PubChem | HSA | PCC | 0.7877665121954682 |
| Morgan | ZIP | PCC | 0.8499367438285369 |
| Morgan | Loewe | PCC | 0.7693666432166798 |
| Morgan | Bliss | PCC | 0.6011541932190027 |
| Morgan | HSA | PCC | 0.7860469156694018 |
| PubChem | ZIP | PCC | 0.8310554090958214 |
| PubChem | Loewe | PCC | 0.7546219316755102 |
| PubChem | Bliss | PCC | 0.5938688616111973 |
| PubChem | HSA | PCC | 0.7752682705805233 |
| RDKit-Chemberta | ZIP | PCC | 0.8215448060817865 |
| RDKit-Chemberta | Loewe | PCC | 0.7412939821887133 |
| RDKit-Chemberta | Bliss | PCC | 0.587841303 |
| RDKit-Chemberta | HSA | PCC | 0.7759567380691348 |
| RDKit-Chemberta-Morgan-PubChem | ZIP | PCC | 0.8510537790244346 |
| RDKit-Chemberta-Morgan-PubChem | Loewe | PCC | 0.7699472078682355 |
| RDKit-Chemberta-Morgan-PubChem | Bliss | PCC | 0.5959768915754906 |
| RDKit-Chemberta-Morgan-PubChem | HSA | PCC | 0.7877665121954682 |
| RDKit | ZIP | PCC | 0.8468517596658265 |
| RDKit | Loewe | PCC | 0.7647935307825297 |
| RDKit | Bliss | PCC | 0.5890645824272698 |
| RDKit | HSA | PCC | 0.7801780918194393 |
| RDKit-Morgan | ZIP | PCC | 0.8378006107276509 |
| RDKit-Morgan | Loewe | PCC | 0.7597596697432023 |
| RDKit-Morgan | Bliss | PCC | 0.5935012794298818 |
| RDKit-Morgan | HSA | PCC | 0.7800439874378179 |
| Chemberta | ZIP | R2 | 0.718688544 |
| Chemberta | Loewe | R2 | 0.5857862088499057 |
| Chemberta | Bliss | R2 | 0.3346511135978677 |
| Chemberta | HSA | R2 | 0.6148460651374524 |
| Morgan-PubChem | ZIP | R2 | 0.7237414068016439 |
| Morgan-PubChem | Loewe | R2 | 0.5903038284596027 |
| Morgan-PubChem | Bliss | R2 | 0.3357785279299996 |
| Morgan-PubChem | HSA | R2 | 0.6159460587586831 |
| Morgan | ZIP | R2 | 0.7217817216575654 |
| Morgan | Loewe | R2 | 0.5878727949671511 |
| Morgan | Bliss | R2 | 0.34253933788369184 |
| Morgan | HSA | R2 | 0.6144565521729796 |
| PubChem | ZIP | R2 | 0.6896059764109969 |
| PubChem | Loewe | R2 | 0.564832719 |
| PubChem | Bliss | R2 | 0.3312451473100315 |
| PubChem | HSA | R2 | 0.5959538248220151 |
| RDKit-Chemberta | ZIP | R2 | 0.674642683 |
| RDKit-Chemberta | Loewe | R2 | 0.548295822 |
| RDKit-Chemberta | Bliss | R2 | 0.31531970054264846 |
| RDKit-Chemberta | HSA | R2 | 0.5969527577001459 |
| RDKit-Chemberta-Morgan-PubChem | ZIP | R2 | 0.7237414068016439 |
| RDKit-Chemberta-Morgan-PubChem | Loewe | R2 | 0.5903038284596027 |
| RDKit-Chemberta-Morgan-PubChem | Bliss | R2 | 0.3357785279299996 |
| RDKit-Chemberta-Morgan-PubChem | HSA | R2 | 0.6159460587586831 |
| RDKit | ZIP | R2 | 0.7169766664163206 |
| RDKit | Loewe | R2 | 0.5828176037812922 |
| RDKit | Bliss | R2 | 0.3210613522405361 |
| RDKit | HSA | R2 | 0.6035907031815577 |
| RDKit-Morgan | ZIP | R2 | 0.7012888730713855 |
| RDKit-Morgan | Loewe | R2 | 0.5734212458202146 |
| RDKit-Morgan | Bliss | R2 | 0.3319048799886968 |
| RDKit-Morgan | HSA | R2 | 0.6040829638496867 |
